# Early origin of sugar sensing in jawed vertebrates

**DOI:** 10.64898/2026.08.28.746343

**Authors:** Qiaoyi Liang, Yasuka Toda, Pauline Affatato, Gentoku Kakizaki, Hitomi Kaname, Kayoko Suzuki, Tae Kuramoto, Meng-Ching Ko, Maxime Policarpo, Julia F. Cramer, Akihiro Itoigawa, Keisuke Furumitsu, Keren R. Sadanandan, Shigehiro Kuraku, Atsuko Yamaguchi, Takashi Hayakawa, Glenn Cockburn, Vincent Van Meir, Ryan K. Tisdale, Sabine Moritz, Ryan M. Carney, Karen Hissmann, Nathaniel S. R. Ng, Borja Reh, Jessica G. H. Lee, Jean-Baptiste Raina, Pablo Oteiza, Naotaka Tsutsumi, Atsuko Yamashita, Yoshiro Ishimaru, Hidenori Nishihara, Maude W. Baldwin

## Abstract

Sweet taste guides animals to consume carbohydrate-rich foods, and many different vertebrate groups, from fish to mammals, rely on sugar-rich fruits or nectar produced by flowering plants (angiosperms). Although the genes encoding T1R2-T1R3, the receptor pair that mammals use to sense sugars, exist in the genomes of many vertebrates, their functions are unclear—and whether sugar sensing arose once early in vertebrate evolution or independently in different lineages after angiosperms evolved is currently unknown. Here, we combined ancestral reconstruction and receptor functional profiling to examine the evolutionary history of T1R taste receptors—including recently-described non-canonical receptors—across all major vertebrate clades. Our results pinpoint the origin of sugar sensing to before the emergence of angiosperms and uncover a myriad of alternative T1R-based sugar-sensing mechanisms, suggesting multiple independent T1R trajectories and revealing uncharted sensory diversity across vertebrates.

---

Sugars are an essential source of energy for many species, and animals all across the tree of life—from mites (*1*, *2*), to butterflies (*3–5*), to hummingbirds (*6–8*), to humans (*9–11*)—consume sugar-rich foods such as fruits (*12*) and nectar (*13*). Although sugars are strongly appetitive for many species, the specific receptors underlying sugar detection have been investigated only in a limited number of species. Sugar sensing has convergently evolved in different clades, with insects relying on ion channels as gustatory receptors (*14*) and vertebrates using the taste receptor type 1 (T1R), a small family of class C G protein-coupled receptors (GPCRs)—in particular, in the form of the T1R2-T1R3 heterodimer—to taste sweet (*15*).

The evolutionary history of sugar detection in vertebrates is not well understood. In mammals, sugar sensing appears to be widespread and likely ancient, with sugar responses documented in T1R2-T1R3 across multiple clades, including primates (*10*, *16*, *17*), rodents (*15*), bats (*18*, *19*), sirenians (*20*), pigs (*21*, *22*) and horses (*21*, *23*)—with secondary losses in certain carnivorous lineages, such as felids (*24*, *25*). Outside of mammals, however, functional responses of T1R2-T1R3 have been observed in only three species: a nectar-feeding gecko (*26*), a frugivorous tortoise and the American alligator (*27*). Although orthologs of the mammalian T1R2 are also present in coelacanth and amphibians (*28–30*), receptors from these species have not responded to sugars in *in vitro* tests (*27*). Bird genomes lack *T1R2* (perhaps due to the carnivorous diet of their theropod ancestors) but multiple lineages including hummingbirds (*8*), woodpeckers (*31*), manakins (*32*), and songbirds (*33*) convergently regained sugar sensing by repurposing the ancestral T1R1-T1R3 savory (umami) taste receptor.

In fish, T1R taste receptors have been shown to be sugar-sensitive in only two species, the herbivorous grass carp (*34*) and the carnivorous seabream (*35*); receptors from other species, such as zebrafish and medaka, have been documented not to respond to sugars (*36*). Interestingly, recent phylogenetic studies have confirmed that the primary fish *T1R* receptor genes in most ray-finned fish genomes encode a receptor clade previously named T1R2, which may in fact not be an ortholog to the mammalian T1R2, but may be more closely related to the umami receptor subunit T1R1: only two non-teleost fish lineages, polypterids and bowfins, possess clear orthologs of the mammalian sugar-sensitive T1R2 (*28–30*, *37*). These studies have also uncovered additional *T1R* gene family members (called non-canonical or ‘novel’ *T1R*) in non-mammalian clades, such as *T1R4*, *T1R5* and *T1R8* in lungfish as well as *T1R4* and *T1R6* in cartilaginous fish, suggesting potential diversity in the mechanisms vertebrates use to sense sugars and amino acids (*29*, *37*).

Sugar-rich fruits and nectar derived from flowering plants (angiosperms) are consumed by many vertebrate lineages, including several clades of specialized birds (*38*, *39*), mammals (*39*, *40*) and non-avian reptiles (*41–43*), as well as frugivorous fish (*44*). Interestingly, a recent study suggests that flowering plants likely originated during the Late Jurassic (150-133 million years ago (Ma)) (*45*), and therefore much later than the emergence of early amniotes (∼318 Ma) (*46*) and tetrapods (∼372 Ma) (*47*). This raises a fundamental question regarding the evolutionary timing of the emergence of sweet taste: given the late appearance of angiosperms, convergent evolution of sugar sensing might plausibly have occurred only after fruits and nectar appeared, with sugar detection then independently evolving in mammals, birds, non-avian reptiles and different clades of fish. Alternatively, the ability to sense sugars may represent an ancient trait shared by most taxa, preceding angiosperms and perhaps later facilitating fruit consumption. When the T1R2-based sugar response first evolved, as well as the extent to which non-canonical T1Rs play a role in sugar sensing, remain unclear.

Here, we used functional profiling of receptors across all major clades to characterize the evolutionary history of T1R taste receptors in vertebrates. Using ancestral sequence reconstruction, we reveal that T1R-mediated sugar sensing evolved far earlier than the emergence of angiosperms, both in the canonical T1R2-T1R3 heterodimers as well as in other non-canonical T1R pairs. Using a series of receptor heterodimers reconstructed at key time-points in vertebrate history, we chart the functional specialization of T1R2-T1R3 and T1R1-T1R3 pairs, documenting the evolution of distinct functions in the canonical sweet and umami taste receptors of mammals. These results provide unexpected insights into multiple molecular mechanisms underlying vertebrate sugar detection, revealing an unexplored landscape of taste diversity across vertebrates.

## Widespread fruit consumption across vertebrates

Although mammals and birds harbor the greatest proportions of fruit-taking species (33% of mammals (2,218 of 6,723 species); and 33.4% of birds (3,719 of 11,131 species)), prevalent fruit consumption has been documented across eight main clades of vertebrates (Fig. 1A-B; fig. S1A; table S1). Interestingly, flowering plants—the producers of fleshy and often sweet fruits (*12*)—emerged later than the amniote and tetrapod ancestors (Fig. 1A). Given this widespread fruit consumption across vertebrates, we tracked the origin of mammalian sugar sensing by first investigating whether diverse fruit-taking lineages across non-avian reptiles (belonging to amniotes) can detect dietary sugars.

**Fig. 1.**
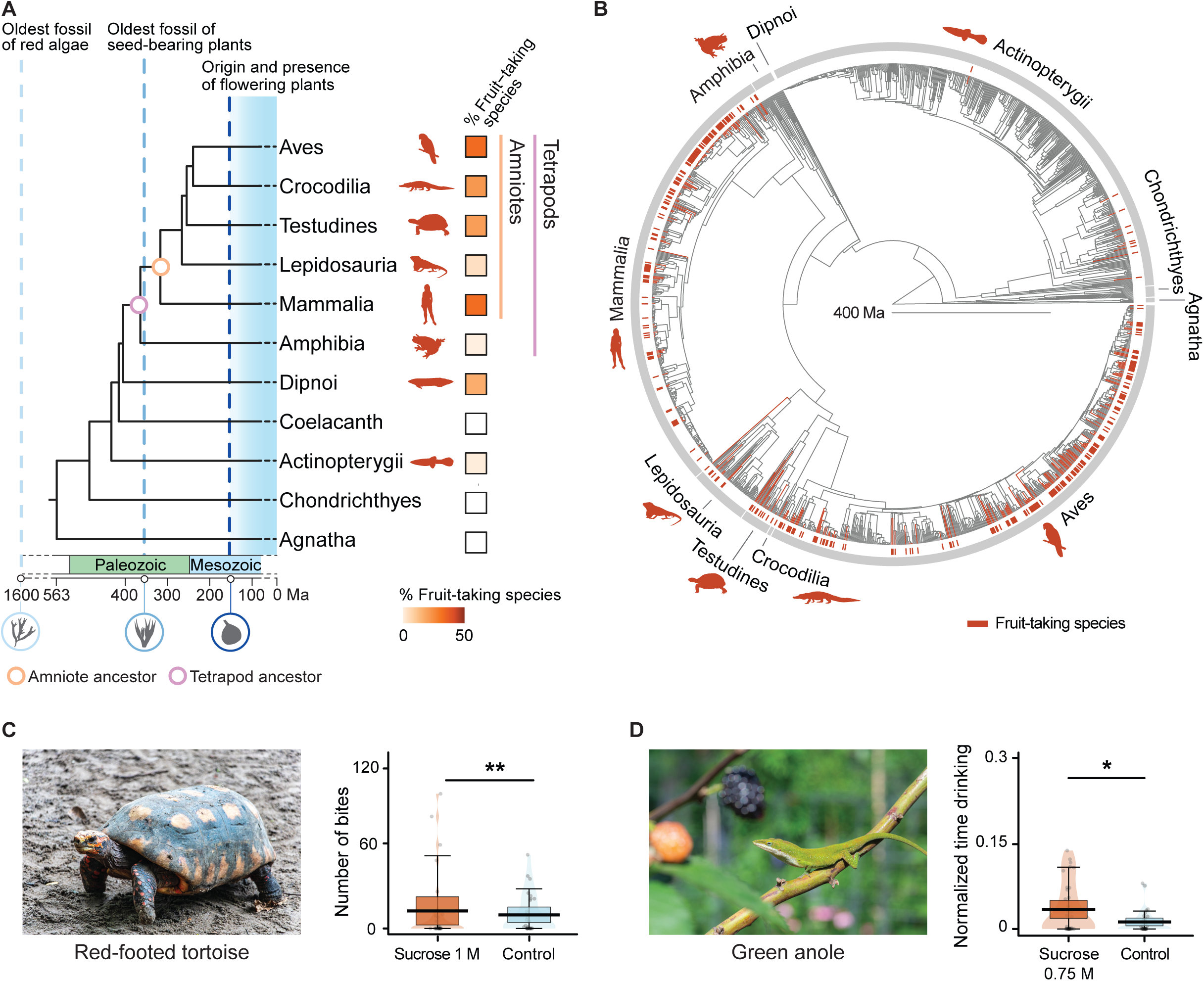
Widespread fruit consumption across vertebrates and preferences for sugar in two non-avian reptiles. (**A, B**) Widespread consumption of fruit. (A) Prevalence of fruit-taking behavior across eight main clades of vertebrates (%: number of fruit-taking species divided by total species count per clade; fig. S1A; table S1; see Methods), and timing of origin of major lineages relative to the evolution of plants—amniotes evolved after seed-bearing plant (spermatophyte) emergence but before the origin of flowering plants (angiosperms). Ma: million years ago. (B) Distribution of fruit consumption across 1,428 vertebrates (with high-quality genome assemblies, see Methods); 318 species are known to take fruits (orange). (**C, D**) Behavioral preferences for sucrose in two non-avian reptiles. (C) The frugivorous red-footed tortoises (*Chelonoidis carbonarius*) preferred agar cubes flavored with sucrose (1 M) over unflavored control cubes within 5 minutes after the first consumption (generalized linear mixed model; ** *p* < 0.01; 7 trials, 5 individuals; fig. S2A; table S3). (D) Green anoles (*Anolis carolinensis*) displayed feeding preferences for 0.75 M sucrose solutions versus water controls (5 minutes after the first drinking bout; linear-mixed model; * *p* < 0.05; 34 trials, 5 individuals; fig. S2B; table S3). See Methods for image credits.

## Behavioral preferences for sugar in two non-avian reptiles

Fruits are frequently consumed by three distinct clades of non-avian reptiles (Fig. 1A-B; fig. S1A). To investigate whether behavioral preferences for sugar exist in these fruit-taking clades (similar to the sugar preference demonstrated in a nectarivorous gecko (*26*)), we conducted brief-access two-choice tests with two fruit-taking non-avian reptiles—red-footed tortoises (*Chelonoidis carbonarius*) and green anoles (*Anolis carolinensis*) (Fig. 1C-D). We found that red-footed tortoises, which are important seed dispersers in savanna areas and surrounding forests in South America (*48*, *49*), showed a significant preference for agar cubes flavored with 1 M sucrose over control (unflavored) agar cubes; this preference was evident even in the first five minutes of the trial, and persisted over the course of 2.5 hours (Fig. 1C; fig. S2A; table S3). In addition, green anoles, carnivores which also opportunistically consume fruits and nectar (*50–53*), displayed an immediate preference for sucrose solutions (0.75 M, also to 0.5 and 0.3 M) over water controls in brief-access presentations (evident in five minutes after the first drinking bout, and remaining consistent over the entire hour; Fig. 1D; fig. S2B; table S3). Therefore, three phylogenetically distinct non-avian reptiles (turtles, anoles, and geckos (*26*)) display preferences for sugar in brief-access trials, suggesting a potentially shared role for sensory (rather than only post-ingestive) detection of sugar.

## Functional responses of T1Rs to sugars and amino acids in non-avian reptiles

Next, to examine if behavioral responses to sugar are reflected in T1R responses *in vitro*, we cloned T1Rs from the oral tissues of four representative non-avian reptiles, and assessed their functional responses in an established cell-based luminescence assay (*54*, *55*). Recently, several additional T1R clades were discovered across jawed vertebrates (*29*, *30*). Therefore, we examined receptors from the canonical T1R clades found in mammals, T1R1, T1R2, and T1R3, which have orthologs in most tetrapods and lobe-finned fish, as well as additional non-canonical T1R clades (including T1R2B, T1R4-8, and T1R3B-C), some of which were present in non-avian reptiles such as anoles (T1R4 and T1R7) (Fig. 2A).

**Fig. 2.**
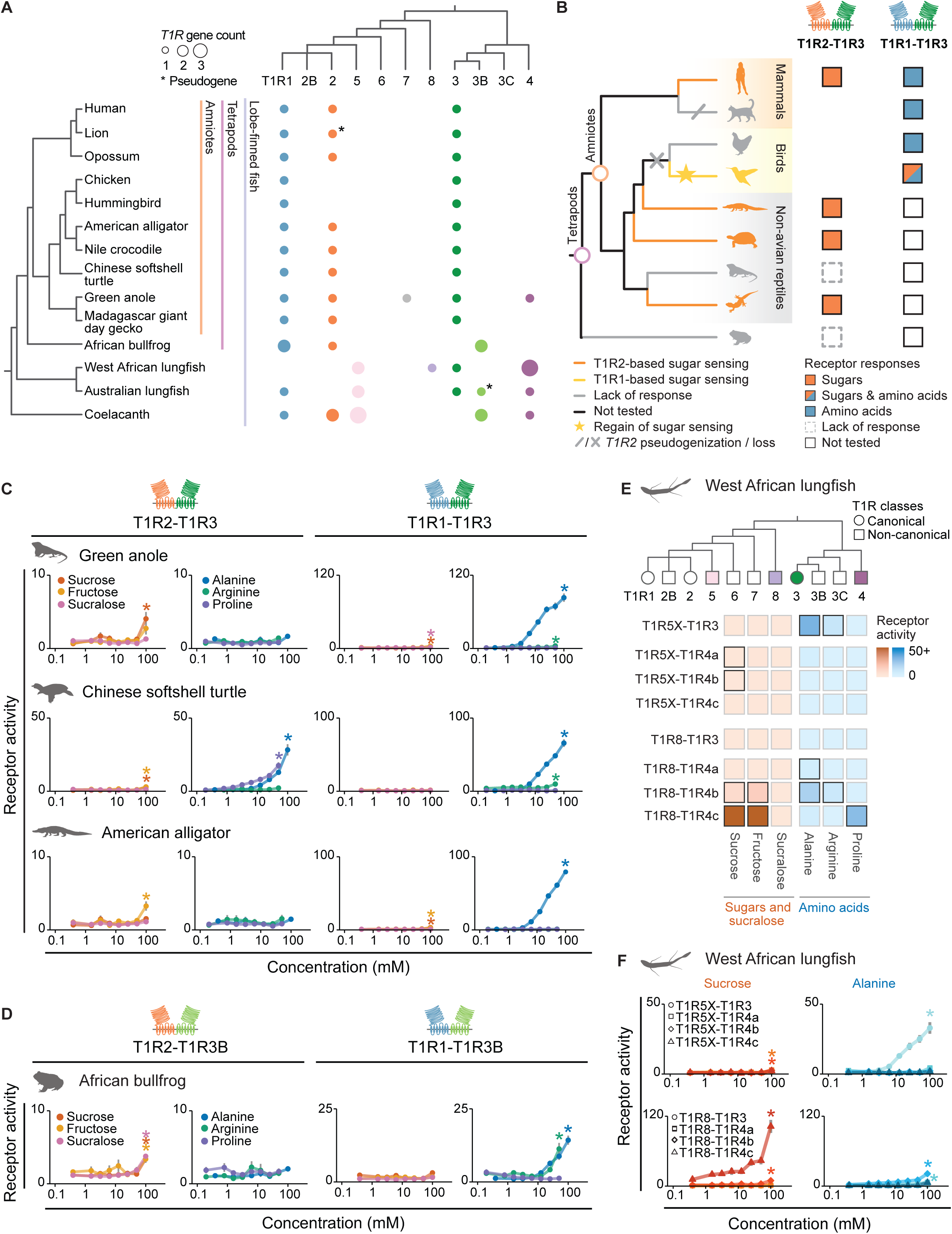
Lobe-finned fish possess multiple canonical and alternative sugar-responsive T1Rs. (**A**) Distribution of *T1R*s among lobe-finned fish (asterisks: pseudogenes). (**B**) Previous studies identified that T1R2-T1R3 (orange-green receptor schematics) responds to sugars in many mammals and several non-avian reptiles (orange branches and squares), whereas nectarivores like hummingbirds (yellow branches) independently regained sugar sensing via T1R1-T1R3 (blue-green schematics; orange-blue squares) following an early loss of *T1R2*. (**C-F**) Heterodimeric receptor responses of non-avian reptiles, amphibians, and lungfish. (C) Green anole, Chinese softshell turtle (*Pelodiscus sinensis*) and American alligator (*Alligator mississippiensis*) possess a sugar-sensing T1R2-T1R3 receptor; unexpectedly, anole and American alligator show responses to sucrose as well as to alanine in the canonical umami receptor T1R1-T1R3. (D) Dose-dependent responses of two African bullfrog (*Pyxicephalus adspersus*) functional receptor pairs: one T1R2-T1R3B pair responds to two sugars and sucralose, whereas one T1R1-T1R3B pair responds only to amino acids (responses of an additional four (out of six total) receptor pairs are shown in fig. S8). (E) West African lungfish (*Protopterus annectens*) have four T1R pairs responsive to sugars and four receptor pairs responsive to three amino acids (all 12 pairs in fig. S9). (F) Dose-dependent responses of eight T1R receptor pairs of West African lungfish. Colored circles and squares in the gene tree illustration in (E) indicate canonical and non-canonical genes present in the lungfish genome. Receptor activity is calculated as the area under the curve (AUC) (x10^3^, relative light units (RLU)) (n = 5-9; mean ± s.e.m; significance (* *p* < 0.05) assessed with one-tailed Welch’s *t*-tests (with the Benjamini-Hochberg (BH) multiple testing correction) comparing between the responses to the highest and the lowest concentrations (with the criteria that the response to the highest concentration was larger than the control cell response (cells transfected with G-protein and photoprotein but without the receptors) or baseline response; ligand concentration series see Methods). Colored cells in the heatmap (E) represent the mean responses of transfected cells to sucrose, fructose, sucralose (all 100 mM, orange colors) and alanine (100 mM) or arginine and proline (50 mM) (blue colors) (dark gray outline: significant responses; light gray outline: non-significant responses; significance assessed using dose-dependent response data in (F) and fig. S9). Silhouette credits: see Methods.

*T1R2* is lost in all bird genomes, but non-avian reptiles possess a T1R2-T1R3 pair, orthologs of the sweet taste receptor in mammals (Fig. 2B). Previous studies have shown variable responses of T1R2-T1R3 to sugar across several non-avian reptiles—the receptors of nectar-taking geckos, desert tortoises and American alligators (*Alligator mississippiensis*) respond to natural sugars, whereas those of bearded dragons, Aldabra giant tortoises and saltwater crocodiles were not responsive to three natural sugars tested (although these receptors could also be nonfunctional, as their amino acid responses were not tested) (*26*, *27*) (Fig. 2B). Here, we demonstrate that T1R2-T1R3 heterodimers of the green anole and the Chinese softshell turtle (*Pelodiscus sinensis*) respond to sucrose (and the latter also responds strongly to several amino acids) (Fig. 2C; fig. S3-5). We additionally tested two crocodilians, confirming the reported response of the American alligator’s T1R2-T1R3 to fructose, and demonstrating a lack of both sugar and amino acid responses in the receptor pair of the Nile crocodile (*Crocodylus niloticus*) (Fig. 2C; fig. S3 and S5B-C). Hence, the function of T1R2-T1R3 as a sugar-sensing receptor pair is likely shared by non-avian reptiles and mammals, suggesting the potential presence of T1R2-based sugar detection in the last common ancestor of extant amniotes.

The lack of responses in two crocodilian T1R2-T1R3 pairs and the unusually restricted receptor response (only to fructose) in the alligator made us curious whether the reduced functionality was due to relaxed selection (perhaps due to their predominantly carnivorous diet, as has been shown for felids (*24*, *25*, *30*)). Using RELAX (*56*), we examined patterns of sequence evolution and detected a signal of relaxation in *T1R2* genes of six crocodilians (but not in *T1R1* or *T1R3*) (fig. S6).

Next, we examined the entire *T1R* repertoire in non-avian reptile genomes and tested other T1R heterodimer combinations. In addition to *T1R2* and *T1R3*, *T1R1* is present in the genomes of almost all bony vertebrates we examined; in mammals, T1R1-T1R3 functions as the umami receptor and responds to amino acids and 5′-ribonucleotides (*57*, *58*). In the four non-avian reptiles we examined, T1R1-T1R3 responds to a wide array of amino acids (Fig. 2C; fig. S3-5). Surprisingly, we also observed responses to several natural sugars by anole and alligator T1R1-T1R3 receptor pairs (Fig. 2C; fig. S4B and S5B), revealing that this type of bifunctional response of the umami receptor (to sugars as well as amino acids) is not unique to specific bird clades, as previously assumed (*59*). Like anoles, American alligators also consume various types of fruits in the wild, whereas only limited fruit consumption has been recorded in crocodiles (*42*). Whether anoles and alligators might taste sugars using either or both T1R2-T1R3 and T1R1-T1R3 taste receptors *in vivo* remains to be explored.

In addition to the canonical T1R1, T1R2, and T1R3 receptors, genes belonging to two non-canonical T1R clades—*T1R4* and *T1R7*—were also observed in the genomes of green anoles (but not crocodilians or softshelled turtles) and were amplified from oral tissues. Using *in situ* hybridization, we identified expression of all five *T1R* genes (and their downstream effector genes (*PLCB2* (phospholipase C beta 2) and *TRPM5* (transient receptor potential cation channel subfamily M member 5)) in putative taste buds of the anole tongue (fig. S4A), suggesting a role for all five T1Rs in taste sensing in anoles. However, in our functional experiments, most tested combinations of receptors (T1R1 and T1R2 paired with T1R4; T1R7 paired with T1R3 and T1R4) appeared nonfunctional, as they did not respond to any of the five sugars, to sucralose (an artificial sweetener and a structural analog of sucrose) or to the 17 amino acids used in an extended ligand panel (with an exception of a small response to trehalose by one receptor pair T1R7-T1R4, see fig. S4B).

## Canonical and alternative sugar-sensing receptors across lobe-finned fish

The widespread sugar responses of T1R2-T1R3 from multiple non-avian reptile species— including species from diverse dietary guilds (such as frugivorous tortoises, nectarivorous lizards and carnivorous crocodilians)—indicate that many non-avian reptiles and mammals may share the same T1R2-T1R3 for sugar sensing, suggesting a possible ancient origin of T1R2-based sweet taste. The mammalian sweet receptor subunit T1R2 is not only present in non-avian reptiles, but also in most amphibians, coelacanth (*Latimeria chalumnae*) (Fig. 2A) and several early-branching ray-finned fish species (*29*). We first examined the functional responses of the canonical and non-canonical receptors from African bullfrog (*Pyxicephalus adspersus*) (with five total receptors forming six different heterodimers) and coelacanth (nine total receptors forming 12 heterodimers). In both species, we coexpressed T1R2 together with T1R3B (a duplication of T1R3 in the bony vertebrate ancestor which was only retained in coelacanth, amphibians, and ray-finned fish (*29*): coelacanth and most amphibians (except axolotl) have lost T1R3, but have two copies of T1R3B) (Fig. 2A). Whereas three coelacanth T1R2-T1R3B pairs appear to be nonfunctional and one pair responds to two amino acids but not to sugars (formed by two copies each of T1R2 and T1R3B; fig. S7), one of the two T1R2-T1R3B pairs from the African bullfrog shows responses to sucrose, fructose and sucralose (Fig. 2D; fig. S8). Additionally, one bullfrog T1R1-T1R3B receptor pair responds strongly to amino acids (Fig. 2D) and another responds weakly to sucrose. Functional responses to six tested ligands were not observed for the three other receptor pairs (fig. S8). Thus, the African bullfrog possesses two sugar-sensing receptor pairs, formed by T1R1 or T1R2 paired with T1R3B, sharing two canonical receptors with the amniote lineage.

Next, we aimed to determine whether non-canonical T1Rs identified in coelacanth and in lungfish (i.e. T1R4, T1R5, and T1R8) can mediate sugar sensing (Fig. 2A). West African lungfish (*Protopterus annectens*) possesses one T1R3 and six members from recently described non-canonical T1R clades. By functionally examining 12 possible heterodimeric combinations, we discovered that four heterodimers containing T1R5 or T1R8 respond to sugars (Fig. 2E-F; fig. S9). Two receptor pairs respond to both sugars and amino acids, among which T1R8-T1R4c exhibits an extremely high sucrose response—higher than other vertebrate receptor responses documented so far—and a relatively small response to proline (Fig. 2E-F; fig. S9). All lungfish are omnivorous, including fruits, seeds, and other plant materials in their diet (*60–66*) (unlike the piscivorous coelacanth (*67–69*), from which none of the nine T1R5 receptor pairs showed functional response *in vitro* (all 18 pairs in fig. S7)). Thus, these retained T1R4, T1R5, and T1R8 members in lungfish may be involved in detecting dietary carbohydrates from the plant materials they consume. Considering the widespread presence of T1R4 and T1R8 across jawed vertebrates (*29*), these sugar-sensing non-canonical T1Rs might have originated before the divergence between lobe-finned and ray-finned fish.

## Diverse sugar-sensing mechanisms across ray-finned fish

Among ray-finned fish, 215 species of 12 orders belonging to teleosts consume fruits in the wild (Fig. 3A; fig. S1B), and some are important seed dispersers in ecosystems such as Amazon floodplain forests (*70*). Interestingly, the fish-specific T1R duplication, here called T1R2B (also known as T1R1B (*30*); depicted here as a polytomy as it is unclear whether it is closer to T1R1 or T1R2), varies extensively in copy number across fish (in contrast to many other T1R clades in vertebrates) and is present in both teleost and non-teleost lineages (Fig. 3B). The origin of this receptor clade predates the origin and diversification of flowering plants (*45*, *71*), and thus, whether phylogenetically distinct herbivorous or fruit-eating teleosts, such as grass carps (*Ctenopharyngodon idella*) (*34*), pacus (*Piaractus brachypomus*) and guppies (*Poecilia reticulata*), convergently obtained sugar sensing via the fish-specific T1R2B, independent of the mammalian sweet receptor, is currently unclear.

**Fig. 3.**
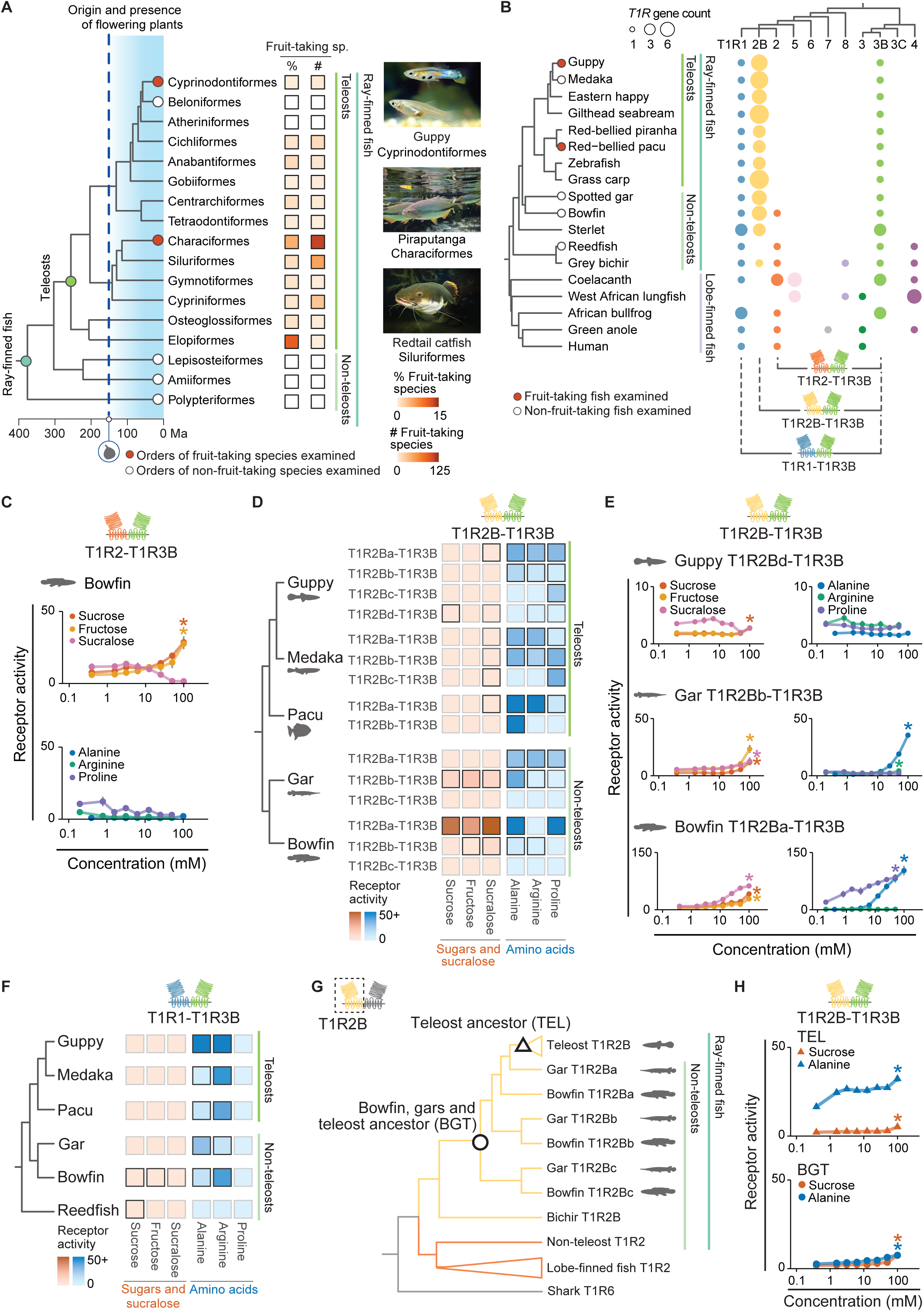
Widespread presence of dietary fruit and T1R-mediated sugar sensing in ray-finned fish. (**A**) Fruits are consumed across 12 orders of ray-finned fish (215 of 2,564 species exampled; 12 of 84 orders; #: number of species; %: number of fruit-taking species divided by total species count per clade; fig. S1B; table S2; see Methods). Fruit-eating behavior is observed mostly in teleosts; the common ancestor of teleosts evolved before the origin and radiation of flowering plants. (**B**) *T1R2B* (yellow), the dominant *T1R* clade and a clade unique to ray-finned fish, may be more closely related to *T1R1* (blue) than to *T1R2* (orange). *T1R2* orthologs are present in 3 non-teleosts. (**C**) Dose-dependent T1R2-T1R3B responses of bowfin (*Amia calva*) show higher responses to sucrose and fructose than to amino acids. (**D, E**) Two to four pairs of T1R2B-T1R3B of five ray-finned fish respond to various ligands (fig. S10-11). Guppies (*Poecilia reticulata*) (a fruit-eating species), spotted gar (*Lepisosteus oculatus*) and bowfin possess T1R2B-T1R3B responsive to both sugars and amino acids. (**F**) Summary of dose-dependent T1R1-T1R3B responses (primarily an amino acid sensor in ray-finned fish, except in bowfin and reedfish, in which they also respond to sucrose). (**G, H**) Ancestral sequence reconstruction of T1R2B from two ancestral nodes and ancestral T1R2B-T1R3B responses. T1R2B-T1R3B reconstructions from the ancestors of all teleosts (TEL, triangles) and of bowfin, gars and teleosts (BGT, circles) show dose-dependent responses to sucrose and alanine (fig. S14). Receptor activity is measured as the AUC (x10^3^ RLU; n = 6-9; mean ± s.e.m; significance (\**p* < 0.05): one-tailed Welch’s *t*-tests with BH corrections, as in Fig. 2; same ligand concentration series as in Fig. 2). Same heatmap colors (D, F) as in Fig. 2E; additionally, the medium gray outline indicates a significant response compared to the lowest documented response of this receptor pair (see Methods). Image credits: see Methods.

Although absent in teleosts, orthologs of mammalian T1R2 exist in three non-teleost species (Fig. 3B). Through receptor profiling, we found that bowfin (*Amia calva*) T1R2 paired with T1R3B is more responsive to sucrose and fructose than to amino acids (receptors from the related reedfish (*Erpetoichthys calabaricus*) were not functional in our assay; Fig. 3C; fig. S11B-C); however, the behavioral sugar preference in bowfin remains to be verified in future studies. Thus, it appears that T1R2-mediated sugar sensing is not restricted to tetrapods and might have been present in the last common ancestor of all bony vertebrates, when T1R2 might have first evolved.

Next, we examined whether fruit-taking teleosts (after the loss of T1R2) might have evolved sugar sensing. We functionally tested different possible T1R combinations from two fruit-eating teleosts, as well as the medaka (*Oryzias latipes*) (medaka receptors were previously shown to be sugar-insensitive (*36*)). We demonstrate that guppies, fruit-eating teleosts, possess one (out of the four total) T1R2B-T1R3B pairs responsive to sucrose (T1R2Bd-T1R3B, Fig. 3D-E; fig. S10A), which might be related to their preference for sugar-rich fruits (*72*) as well as their behavioral preference for sucrose in lab experiments (*73*). We also tested the three T1R2B-T1R3B pairs previously reported from medaka (*36*), observing consistent responses to amino acids in three T1R2B receptor pairs (Fig. 3D; fig. S10B). We observed that both guppy and medaka T1R2Bc-T1R3B pairs are primarily tuned to proline (Fig. 3D; fig. S10A-B). Unexpectedly, we did not document any sugar responses in the two tested T1R2B receptor pairs (which exhibit strong amino acid responses) of the heavily frugivorous red-bellied pacu, a representative of another distinct order (Characiformes, a dietarily-diverse clade containing the most fruit-eating species among ray-finned fish) (*44*) (Fig. 3D; fig. S10C). Although pacu receptors do not respond to sucrose, one receptor pair shows a small response to sucralose (Fig. 3D; fig. S10C); whether these receptors might be associated with fruit-eating behavior in pacus (or whether additional untested T1R2B copies exist in the genome, as a complete genome assembly for this species was not available at the time of testing) remains to be explored.

Together with previous research, these results suggest that sugar sensing mediated by T1R2B is present in at least three phylogenetically distinct teleost species (grass carp (*34*), seabream (*35*) and guppy), in contrast to the previously reported sugar-insensitivity in zebrafish and medaka (*36*)—and in these lineages, is often mediated by different T1R2B copies (as T1R2B duplications are often lineage-specific and not shared across species, our naming convention (ie. T1R2Ba, T1R2Bb) refers here to copies within a species, not to orthologs across different fish lineages). To examine whether this pattern potentially reflects a shared ancestral response to sugars across ray-finned fish, we tested T1R2B-based receptor pairs of non-teleosts to assess whether sugar sensing might have been present ancestrally. We identified sugar-responsive T1R2B-T1R3B pairs in two non-teleosts, the spotted gar (*Lepisosteus oculatus*) and the bowfin: interestingly, these heterodimers show robust sugar responses, and also strong responses to three amino acids (i.e. gar T1R2Bb-T1R3B, bowfin T1R2Ba-T1R3B and T1R2Bb-T1R3B; Fig. 3D-E; fig. S11A-B). Similar to teleosts, both gar and bowfin each possess a T1R2B-T1R3B exhibiting high sensitivity to proline (T1R2Ba-T1R3B; Fig. 3D; fig. S11A-B), suggesting that the perception of proline is widespread across ray-finned fish— although mediated by different T1R2B-T1R3B heterodimers.

Similar to the umami receptor T1R1-T1R3 of amniotes, ray-finned fish T1R1-T1R3B pairs are primarily responsible for amino acid detection (i.e. responses to alanine and arginine tested here); these receptors in bowfin and reedfish also respond weakly to sucrose (Fig. 3F; fig. S10-11). Thus, T1R1-T1R3B of ray-finned fish primarily functions as an amino acid sensor, but in non-teleosts, it can also be involved in sugar sensing.

Given the widespread presense of sugar responses of T1R2B-based receptor pairs—but not in the T1R1 pairs—across ray-finned fish, we wondered if sugar sensing mediated by T1R2B-T1R3B might represent an ancestral trait (potentially lost in later lineages). We reconstructed and functionally tested ancestral receptor pairs from two phylogenetic nodes (Fig. 3G; fig. S14A-B) and found that sugars and amino acids can elicit responses of T1R2B-T1R3B pairs reconstructed from the common ancestor of bowfin, gars and teleosts (BGT), as well as the ancestor of all teleosts (TEL) (Fig. 3G-H; fig. S14C). Interestingly, the ancestral heterodimer of all teleosts is more sensitive to alanine than to sucrose compared to reconstructions from the previous node, implying an increased propensity in amino acid sensing with a retained sucrose response in early teleosts after diverging from non-teleosts. Therefore, sugar sensing via T1R2B-T1R3B might have been present early in ray-finned fish, predating the origin of flowering plants. Subsequently, some teleost lineages might have lost the ability to sense sugars, whereas others might have retained the ancestral sugar-sensing function, thereby facilitating the detection or consumption of sugar-rich fruits in their diets.

## Early origin of sugar sensing in jawed vertebrates

*T1R* genes, unique to vertebrates, are absent from the genomes of living jawless fish (hagfish and lampreys), and first found in cartilaginous fish such as sharks and rays (*29*, *30*, *37*, *74*). Cartilaginous fish diverged from the other jawed vertebrates ∼450 Ma (70) and also possess lineage-specific *T1R6* and *T1R3C* as well as *T1R4* genes. To examine whether receptors from this lineage respond to sugars (as a previous study had identified amino acid responses in one receptor pair from the non-elasmobranch Australian ghostshark (*Callorhinchus milii*) (*29*)), we tested the four receptors encoded by *T1R*s mined from the genome of the great white shark (*Carcharodon carcharias*) as well as T1R6-T1R3C cloned from the red stingray (*Hemitrygon akajei*) oral tissue. We identified clear sucrose responses when pairing T1R6.1 (one of the ancient T1R6 orthologs shared across cartilaginous fish) with T1R3C in the stingray and great white shark; these heterodimers are also sensitive to proline and alanine (Fig. 4A; fig. S15A). Thus, we hypothesized that sugar detection via T1Rs might have evolved early in jawed vertebrates.

**Fig. 4.**
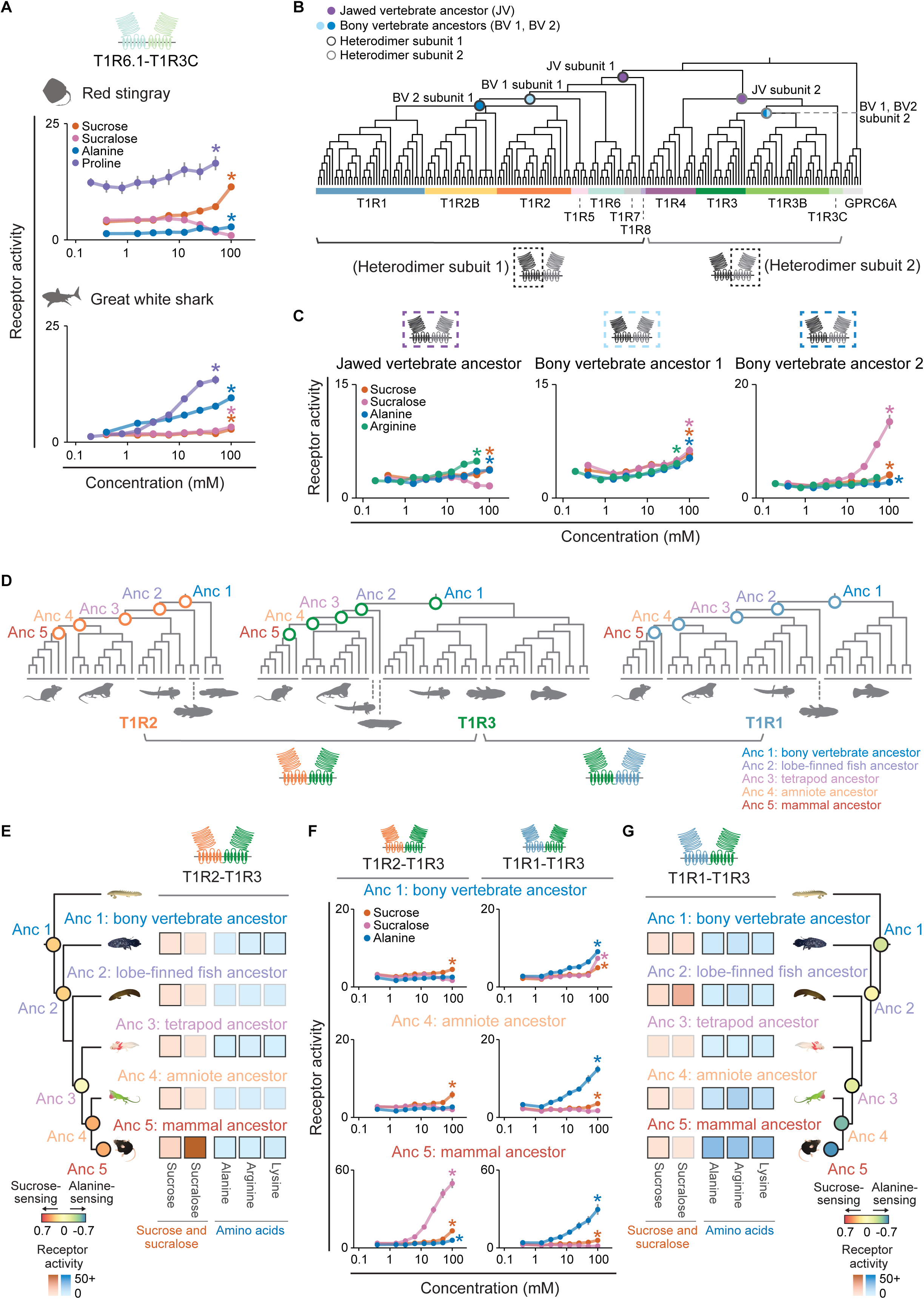
Early origin of sugar sensing in jawed vertebrates and subsequent functional specialization into sweet and umami receptors. (**A**) Functional responses of T1R6.1-T1R3C pairs of the red stingray (*Hemitrygon akajei*) and great white shark (*Carcharodon carcharias*). (**B, C**) Ancestral reconstruction and functional responses of early vertebrate nodes. (B) Five ancestral T1Rs from early vertebrates were reconstructed (using GPRC6A as an outgroup; fig. S17) and expressed in functional assays. (C) Dose-dependent responses of three ancestral receptor pairs to sucrose, sucralose and two amino acids, indicating an early emergence of a small response to sucrose in the heterodimer of the jawed vertebrate ancestor. (**D-G**) Ancestral reconstruction and dose-dependent responses of T1R1, T1R2, and T1R3 receptors. (D) Topologies used for ancestral reconstruction of each receptor across bony vertebrates. (E) Response summary of five ancestral T1R2-T1R3 receptor pairs; colored circles on nodes on the tree represent an index of relative sucrose to alanine response (the difference of average responses to 100 mM sucrose and 100 mM alanine divided by the sum of average responses to sucrose and alanine; see Methods)—a shift towards a predominant response to sucrose (orange circles) appears to have evolved subsequently in bony vertebrates, and is most prominent in the ancestor of mammals. (F) Dose-dependent responses of ancestral T1R2-T1R3 and T1R1-T1R3 from the bony vertebrate (Anc 1), amniote (Anc 4) and mammal ancestors (Anc 5). (G) Response summary of five ancestral T1R1-T1R3 receptor pairs; blue index circles indicate a heightened relative response to alanine (although sucrose responses are also retained in four out of five pairs). Receptor activity is calculated as AUC (x10^3^ RLU; mean ± s.e.m; significance (* *p* < 0.05) assessed with one-tailed Welch’s *t*-tests with BH corrections, as in Fig. 2 (n = 6-8; ligand concentration series see Methods). Same heatmap colors (E, G) as in Fig. 2E (derived from data in fig. S18).

To investigate whether T1R-mediated sugar sensing might have been present ancestrally, we inferred five ancestral sequences from the different T1R clades at the jawed vertebrate and bony vertebrate nodes in the phylogeny, and functionally tested these heterodimers (Fig. 4B; fig. S17). We found that a jawed vertebrate ancestral heterodimer shows robust responses to amino acids (jawed vertebrate ancestor (JV); Fig. 4C), consistent with the amino acid activation properties of other class C GPCR members (*75–78*). In addition, it also displays a small response to sucrose (Fig. 4C). This early emergence of sugar detection is also consistent with the sugar responses we documented in the receptor pairs from representative species across all extant jawed vertebrate clades.

Compared to the jawed vertebrate ancestor, two receptor pairs in the bony vertebrate ancestor show elevated responses to sucrose and sucralose (bony vertebrate ancestor 1 and 2 (BV 1 and BV 2); Fig. 4B-C). Thus, enhanced sugar sensing might have first occurred in the last common ancestor of all bony vertebrates, involving the common ancestor of the receptor clades T1R1, T1R2, T1R2B and T1R5 (Fig. 4B-C).

## Functional specialization of the sweet and umami receptor

When did the bifunctional ancestral receptor pairs become specialized into the dedicated sweet and umami receptors characterized in mammals? To address this question, we functionally examined ancestral T1R2-T1R3 and T1R1-T1R3 pairs inferred from five major phylogenetic nodes across bony vertebrates (Fig. 4D; fig. S17). Most ancestral heterodimers respond to both sucrose and amino acids (Fig. 4E-G; fig. S18), consistent with patterns observed in some receptor pairs from representatives of diverse vertebrate clades (Fig. 2-3). To assess the relative response of a given heterodimer to sucrose or alanine (as a representative sugar and amino acid), we calculated a response index (by taking the difference of average responses to 100 mM sucrose and 100 mM alanine divided by the sum of sucrose and alanine average responses). Notably, we observed a heightened relative sensitivity to sucrose in ancestral T1R2-T1R3 compared to T1R1-T1R3 across bony vertebrates, whereas ancestral T1R1-T1R3 pairs display enhanced relative responses to amino acids compared to sucrose (Fig. 4E-G; fig. S18). Ancestral T1R2-T1R3 and T1R1-T1R3 of all mammals show the highest degree of functional specialization into distinct sucrose- and alanine-sensing receptors (orange and blue circles on the trees in Fig. 4E, 4G). In addition, the ancestral T1R2-T1R3 of all mammals displays the highest response to sucralose among five tested ligands (Fig. 4E-F; fig. S18), in line with the strong affinity of sucralose binding observed in human and mouse sweet taste receptors (*10*, *15*). The mammalian ancestral T1R2 also shares most of the critical residues for sucralose binding and heterodimer interaction with human TAS1R2 (*79–81*) (fig. S19), indicating that humans might have inherited an enhanced sweet taste receptor from the last common ancestor of all extant mammals. Altogether, we revealed that after an acquisition of sugar detection in the ancestral heterodimeric receptors, specialized sweet and umami receptors may have evolved subsequently during vertebrate evolution.

## Discussion

Here, we aimed to understand when T1R-based sugar detection first arose in vertebrates. We focused first on the canonical sweet (T1R2-T1R3) and umami (T1R1-T1R3) receptors, and then extended our functional profiling to include the non-canonical T1R repertoire, revealing diversity in copy number as well as unexpected functional shifts in other heterodimers. Collectively, these results point to an earlier gain of sugar sensing than had previously been appreciated (*27*), and to a plethora of different sugar sensors in different lineages.

Tracking the origin of T1R2-T1R3 sugar responses revealed a clear pattern of consistent sugar responses across bony vertebrates. Present first unequivocally in three early-branching ray-finned fish species (but lost in teleosts), T1R2, when paired with T1R3B, responded to sugars in both non-teleost bowfins as well as African bullfrogs (Fig. 2D and 3C): as both species are carnivorous, these results suggest a possible ancient and ancestral role for sugar sensing in this receptor heterodimer (instead of a response restricted to specialized sugar-consuming lineages). Importantly, both extant and ancestral T1R2-T1R3 from reptiles as well as the mammalian ancestor were clearly responsive to sugars (Fig. 2C and 4F; fig. S18 and S22A). Therefore, given the indispensable role of T1R2 in sugar sensing, it seems likely that sugar responses in this canonical heterodimer may have evolved early, at least in the ancestor of all bony vertebrates, predating the emergence of angiosperms.

In birds, T1R2 appears to have been lost early, sometime after the split from the last common ancestor with living crocodilians (since *T1R2* does not even appear as a pseudogene in bird genomes). Interestingly, we observed strong signatures of relaxation in crocodilian *T1R2* genes. T1R2-T1R3 from two out of three tested crocodilian species showed no response *in vitro*, whereas the alligator receptor pair had a restricted response to fructose. Whether these selection signatures and receptor response profiles suggest a signal of an early shift towards carnivory in early archosaurs (*82*, *83*) together with the accompanying relaxed pressure on *T1R2*, followed perhaps by a regain of sugar-sensing in alligators, remains to be investigated. Alternatively, these may be separate events, with bird *T1R2* undergoing relaxed selection and loss later in evolution, perhaps in their carnivorous theropod ancestors (*84*, *85*). Among extant taxa, fruit consumption by American alligators—but not crocodiles—is frequently documented (*42*) and anecdotally, alligators are known to consume marshmallows provided by tourists (*86*). The repeated evolution of herbivorous crocodyliforms has also been inferred from the fossil record (*87*): sugars, therefore, may represent an important dietary component for at least some crocodilians, perhaps mediated by diverse detection mechanisms. However, which receptor pair may drive dietary sugar preferences in alligators (as both T1R2-T1R3 and T1R1-T1R3 in alligators respond to natural sugars) remains unclear.

In addition to T1R2-T1R3, we also functionally profiled other T1R heterodimers and discovered unexpected bifunctional responses (to amino acids and sugars). We started with the umami receptor (T1R1-T1R3) and, surprisingly, these heterodimers exhibited bifunctional responses in two non-avian reptiles (anoles and alligators) (Fig. 2C) - a response profile previously thought to be unique to ‘repurposed’ bird umami receptors in sugar-consuming lineages (*8*, *31–33*). Ancestral receptor profiling underscores this broad tuning: although the mammalian ancestral receptor T1R1-T1R3 has the highest relative amino-acid-to-sugar response, T1R1-T1R3 receptor pairs in three earlier nodes are not strictly specialized (Fig. 4F, 4G). This pattern of bifunctionality was also observed in many of the non-canonical pairs we tested, which is not surprising given that both umami and sweet are appetitive tastes important for feeding and survival. After an early loss of the mammalian ortholog T1R2, ray-finned fish also gained and expanded a lineage-specific clade, T1R2B, responsive to both sugars and amino acids when pairing with T1R3B. We observed the presence of bifunctionality in the ancestral T1R2B-T1R3B pair in early ray-finned fish and, subsequently, a functional shift to an enhanced amino acid detection (with retained sucrose response) in the ancestral pair of all teleosts (Fig. 3H). Consistent with this ancestral response, we observed broadly tuned T1R2B-T1R3B heterodimers retaining high sugar responses in non-teleost species (i.e. bowfins and gars); by contrast many teleost T1R2B-T1R3B pairs were more responsive to amino acids with low (or absent) responses to sugars or sucralose. Overall, however, most receptor pairs (including some extant and ancestral T1R2-T1R3 and many fish-specific T1R2B-T1R3B heterodimers) were broadly tuned.

As flowering plants clearly evolved after the emergence of the amniote ancestor (*45*, *46*), our results suggest that a low sugar response existed in an ancestrally bifunctional T1R pair far earlier than the time when angiosperms evolved. Later increases in response magnitudes, as well as a subsequent division of function (partitioning sweet and umami) in the lineages leading to mammals may reflect changes in the availability of fruits and nectar as dietary sources. Indeed, widespread fruit consumption has been documented across many tetrapod clades, and fruit is speculated to have been an important diet item for ancient mammals and reptiles alike. Fossil records indicate that large fleshy fruits were already present by the Late Cretaceous (∼74.6 Ma) (*88*) and many groups of extinct mammals may have consumed fruits, including multituberculates (*89*, *90*), gondwanatherians (*91*), marsupial relatives (Polydolopimorphia) (*92*) and pan-primates (Plesiadapiforms) (*93*). It is possible that this enhanced sugar sensing might have enabled early arboreal mammals to occupy available dietary niches during the drastic diversification of angiosperm fruit sizes (*40*), by fine-tuning their sweet taste sensitivity to include sweet foods in their diets—a process that may have convergently evolved in specialized nectar- and fruit-taking birds, which regained sugar sensitivity using T1R1-T1R3 (interestingly, data from extant birds suggests T1R1-T1R3 bifunctionality may not have been ancestral, but secondarily re-evolved in hummingbirds (*8*), woodpeckers (*31*), manakins (*32*) and songbirds (*33*)).

The ancestral function of T1Rs in vertebrates remains an interesting and unresolved question. In ancient vertebrates, T1Rs may have played a sensory role. *T1R* and their co-effector genes are expressed in taste buds and oral tissues in non-teleosts, such as bichirs (*29*), and in various cartilaginous fish (*94*, *95*), suggesting a possible sensory role in food assessment and feeding behavior. This early receptor response to sucrose indicates that dietary sugars may have been important sources of nutrients long before angiosperms originated. Indeed, highly specific and specialized receptors for individual sugars (e.g., fructose, galactose, glucose, maltose, ribose, and trehalose) are present in many bacteria, archaea and microalgae (*96*, *97*), allowing them to detect and move towards sugar-rich hotspots (*98*). In addition, macroalgae, a polyphyletic group of carbohydrate producers that evolved long before the emergence of seed plants, dominated the ancient ocean when jawed vertebrates first evolved (*99–101*) and were consumed by early Ediacaran animals (558 Ma) (*102*). Sugar-rich algae (including both macroalgae such as seaweed (*103*) and microalgae such as diatoms (*104*, *105*)) are still included in the diets of many species across extant vertebrates (*106–109*). Whether later clades of jawed vertebrates use T1R taste receptors to detect angiosperm-derived sugars (e.g., from fallen fruit, consumed by fruit-eating species such as pacus (*110*) and guppies (*111*), or from aquatic plants, consumed by herbivorous fish such as grass carp (*112*, *113*)), or other possible non-angiosperm sources of sugars (such as algae, or such as sugar-rich gymnosperm fleshy seeds (*114–117*) consumed by multiple dinosaur lineages in the past (*118–120*) as well as by extant amniotes (*121*, *122*)) remains to be tested.

In addition to expression in sensory organs, T1R expression and extraoral functions have also been documented in other tissues, and it is possible that ancestrally, T1Rs may have played other non-sensory roles before being co-opted to detect sugars and amino acids in taste buds. In mammals and teleosts, extraoral T1Rs suggest diverse functions, including nutrient sensing and hormone regulation in the digestive tract (*16*, *123–128*), glucose sensing in the brain (*34*, *129*, *130*), involvement in pathogen responses in the airway (*131*, *132*), and prey detection with modified sensory appendages (*133*). Further investigation into the “nonfunctional” T1R receptors (which do not respond to canonical taste compounds) and into T1R expression patterns in a wide panel of tissue types across jawed vertebrates might provide insight into the overlooked and potentially ancestral functions of this receptor family.

Overall, we demonstrate surprising functional diversity of the T1R receptor family and shed light on the origin and functional specialization of an important sensory modality in vertebrates. This work lays the foundation for future examinations of the underlying structural and molecular changes, and raises questions about the role of sugar sensing in vertebrate diets, as well as the sensory underpinnings of ancient plant-animal interactions.

## Acknowledgments

We thank Prof. H. Matsunami (Duke University) for providing the HEK293T cells, H. Fricke and P. Bartsch for the coelacanth tissue sampling, and N. Rattenborg for the Nile crocodile tissue collection. We also thank C. Reusch, L. Gaitanos, M. Hertel, S. Kuhn, A. Bakker, C. Baumgartner and D. Mendez Aranda for assistance in molecular experiments. We appreciate the animal caretaking by F. Lehmann and D. Witkowski in Seewiesen as well as the staff of the RepTopia in Singapore Zoo. Claude 3.5 Sonnet was used to assist in creating the heatmaps for the cell assay data in R.

## Funding

This work is supported by the Max Planck Society (to M.W.B.), KAKENHI grants from the Japan Society of Promotion of Science (18KK0166 to Y.I.; 20H02941, 22KK0079, 23H02168, 25H01362, 26K01695 to Y.T.; 19H03272 to H.N.), the International Collaborative Research Promotion Project from Meiji University (to Y.T.), the JST FOREST Program (Grant Number JPMJFR220C, Japan) (to Y.T.), the Lotte Shigemitsu Prize (to Y.I., Y.T.).

## Author contributions

Conceptualization: Q.L., Y.T., A.Ys., Y.I., H.N., M.W.B.

Formal analysis: Q.L., Y.T., P.A., G.K., M.P., K.R.S., S.K., A.Yg., T.H., A.Ys., H.N., M.W.B.

Funding acquisition: Y.T., A.Ys., Y.I., H.N., M.W.B.

Investigation: Q.L., Y.T., P.A., G.K., H.K., K.S. T.K., J.F.C., A.I., A.I., S.K., T.H., G.C., A.Yg., Y.I., H.N., M.W.B.

Methodology: Q.L., Y.T., P.A., M.K., M.P., K.S., A.I., N.T., A.Y., Y.I., H.N., M.W.B.

Project administration: Q.L., Y.T., P.O., V.V.M., N.S.R.N., B.R., J.G.H.L., A.Ys., Y.I., H.N., M.W.B.

Resources: R.K.T., S.M., R.M.C., K.H.

Visualization: Q.L., Y.T., M.K., M.P., P.O., A.Ys., H.N., M.W.B.

Writing – original draft: Q.L., Y.T., A.Ys., H.N., M.W.B.

Writing – review & editing: all authors

## Competing interests

Authors declare that they have no competing interests. R.K.T. is a current employee of F. Hoffmann-La Roche Ltd.

## Data and materials availability

All data and codes used in main text or supplementary materials are accessible at the Dryad depository (https://doi.org/10.5061/dryad.tqjq2bwfh).

## Supplementary Materials

*Materials and Methods*

*figs. S1 to S22*

*tables S1 to S4*

## Notes

### Competing Interest Statement

The authors have declared no competing interest.

