## Supplementary Materials for "Early origin of sugar sensing in jawed vertebrates"

**The PDF file includes:**

Materials and Methods
figs. S1 to S22

**Other Supplementary Material for this manuscript includes the following:**

Tables S1 to S4

#### Materials and Methods

*Diet data and taxonomic relationships of vertebrates used for visualizing fruit and seed consumption, and timing of the origin of particular clades*

Dietary data for mammals and birds were extracted from the EltonTrait database (*134*): species with >1% of fruits and seeds in their diet were considered fruit- and seed-taking species. Squamate species were scored as fruit- and seed-taking by searching for the keywords “fruit,” “berries”, “berry”, and “seed” in the “diets: comments” column of a global trait database for lizards (*52*); additional fleshy fruit-taking species mentioned in a subsequent publication (*41*) were also included. Lists of crocodilian and turtle species documented as consuming fruits and/or seeds were extracted from two reviews (*42*, *43*). We identified five out of 6,776 amphibians that take fruits and seeds from the AmphiBIO database (*135*). Data on coelacanth and lungfish were compiled from 11 publications: whereas coelacanths primarily consume fish (*67–69*), all lungfish are generalists, including invertebrate and vertebrate prey as well as plant material in their diet (*60–66*). In particular, Australian lungfish (*Neoceratodus forsteri*) have been observed feeding on fruits and seeds of *Eucalyptus* (*60*) and seeds have also been identified in the gut contents of West African lungfish (*Protopterus annectens*) (*64–66*). Lists of fruit- and seed-eating ray-finned fish species were compiled from references cited in two reviews (*44*, *110*) and by searching for keywords “fruit”, “seed” and “berries” in the diet items table of the FishBase database using the R package “rfishbase” (*136*, *137*) (2,564 of 34,635 species examined). Since the guts and feces of both frugivores and granivores can contain seeds, differentiating between the two diet types based on gut contents or fecal samples alone can be challenging. Therefore, we presented fruit- and seed-taking species together as ‘fruit- & seed-taking species’ and used the more restricted term ‘fruit-taking species’ to denote species in which fruits were specifically mentioned in the diet, or fruit remains were found in gut contents or feces. The lists of names of fruit- and seed-taking ray-finned fish and non-avian reptile species names were checked for synonyms to remove redundancy using FishBase and Reptile Database (*138*). Order and family information was obtained from the NCBI database (*139*), FishBase and Reptile Database.

To calculate the percentages of ‘fruit-taking species’ (Fig. 1A) and ‘fruit- & seed-taking species’ (fig. S1A), total counts of extant species per clade were taken from the Mammal Diversity Database (v2.3) (*140*) for mammals, from AviList (v2025) (*141*) for birds, from Reptile Database (05/2025) (*138*) for non-avian reptiles, from AmphibiaWeb (09/2025) (*142*) for amphibians, and from FishBase (04/2025) (*137*) for lungfish, coelacanth, ray-finned fish, cartilaginous fish and jawless fish (table S1). For Fig. 3A and fig. S1B, total species counts of fish per order were extracted from “rfishbase” package in R to calculate the percentages of ‘fruit- & seed-taking species’ and ‘fruit-taking species’ per order (table S2).

The vertebrate phylogenies shown Fig. 1A and fig. S1A were generated in TimeTree (v5) using representative species per clade (*143*) and were modified by adjusting branch lengths based on previous studies (*71*, *144–147*). The phylogenetic tree of 1,428 vertebrate species in Fig. 1B was modified from a previously published phylogeny (*30*); this dataset selected species that had genome assemblies with BUSCO (Benchmarking Universal Single-Copy Orthologs; v5.1.2) completeness (*148*) of greater than 80% (except for 5 jawless fish whose genome assembly completeness might not be reliably assessed with

BUSCO gene sets). In Fig. 3A and fig. S1B, the order-level phylogeny of ray-finned fish was modified from the backbone of Rabosky *et al.* (*149*) with minor adjustments(*71*, *150–152*).

In Fig. 1A and 3A, we indicated the estimated origin of flowering plants (angiosperms; Late Jurassic (150-133 Ma)) based on a recent study (*45*). The emergence of seed-bearing plants (spermatophytes, including gymnosperms and angiosperms) and red algae (rhodophytes) in Fig. 1A were approximated using the oldest undisputed fossil of each group (*101*, *153*).

*Silhouettes and animal images*

The silhouettes were downloaded and modified from PhyloPic (https://www.phylopic.org/) under the Universal Public Domain license CC0 1.0; several were from Mr. Fulano (*Xenohyla truncata*), Soledad Miranda-Rottmann (*Pelodiscus sinensis*), Milton Tan (*Lepisosteus oculatus*) and Roberto Diaz Sibaja (*Neoceratodus forsteri*). The illustrations in Fig. 4E, 4G and S22C were attributed to Daniel Ocampo Daza. The animal photographs in Fig. 1C-D and 3A were obtained from Rudolf Ernst (*Chelonoidis carbonarius*), Marianne Pfeil (*Anolis carolinensis*), Luciano Queiroz (*Brycon hilarii*) and Kletr (*Phractocephalus hemioliopterus*) and licensed via Adobe Stock Photo (https://stock.adobe.com), as well as from Deacon *et al.* (*154*) (*Poecilia reticulata*).

*Two-choice behavioral tests*

The experiments with five juvenile red-footed tortoises were conducted in the RepTopia (formerly 'Reptile Garden') exhibit section of the Singapore Zoo following protocols approved by the Mandai Wildlife Group (formerly known as Wildlife Reserves Singapore) in Singapore. Behavioral experiments with 15 captive-bred juvenile green anoles housed in the animal facility of the Max Planck Institute for Biological Intelligence, Seewiesen, Germany were performed following the regulations of the European Union and the German Animal Welfare Regulation Governing Experimental Animals (TierSchVersV). All animals had access to water *ad libitum* and were fed after the experiments.

After acclimatization to the filmed two-choice set-up in their home terrarium (fig. S2A right), five tortoises were given a pair of dishes with 10 g each of 1% orange-colored agar cubes, prepared with either 1 M sucrose solution or tap water. Trials were conducted once a day, and the stimulus presentation side was first randomized and then switched in the subsequent trials to avoid side biases. A USB video webcam was used to record the behaviors of five tortoises for up to 2.5 hours per trial at 10 frames per second (fps) and the numbers of bites from each dish were manually scored by a single observer (Q.L.) (blinded to which side the stimulus was present on) using the Behavioral Observation Research Interactive Software (BORIS v3.60) (*155*). To estimate the sucrose preference of tortoises, we applied generalized linear mixed models with a Poisson distribution for each time window (5, 10, 20, 60, 150 minutes) after the first bite using the lme4 package (*156*) in R (4.5.0) (*157*). In these models, we defined the number of bites as the response variable, the treatment choice as the fixed effect as well as the trial ID and individual ID as random effects.

After getting accustomed to the filmed two-choice test setup in its home terrarium (fig. S2B right), each anole was presented with two side-by-side feeders and its responses were recorded for an hour (using Flir Chameleon3 or BlackflyS camera with a Fujinon lens

YV2.8x2.8SA-2 at 30 fps). Each animal was presented with a pair of feeders, with one side filled with 1 mL of sugar solution (sucrose, 0.3, 0.5 and 0.75 M; glucose, 0.75 M; fructose, 0.75 M) and the other containing deionized water, as a control. Two trials were conducted each day with a 30-minute break in between. The presentation sides were randomized in the first trial and then swapped for the second trial to avoid a potential side bias. From the recorded videos, the amount of time spent drinking from each side of the paired feeders was scored as one of three behavioral classes in DeepEthogram (v0.1.4) (*158*) (three classes: "background", "drinking left", "drinking right"; drinking was scored as when the anole had its tongue or its whole mouth in contact with or submerged in the liquid; images were resized to 224x224 pixels; 8 batch sizes and other default hyperparameters were used), followed by manual confirmation of predictions by two experimenters (P.A. and Q.L.) blinded to the stimulus presentation side). To test whether and when anoles displayed behavioral preference towards any sweet stimulus over water, we quantified the proportional drinking time of each side within a series of time windows after the first consumption from either side (time spent drinking divided by 5, 10, 20, 60 minutes). We applied linear mixed models and defined the normalized drinking time (normalized using square-root and z-scale transformations) as the response variable and the choice for stimulus or water as the fixed effect using lme4 in R. To account for multiple testing and within-individual variation, we specified the trial ID and individual ID as random effects.

*Animal tissue collection*

Green anole, axolotl (*Ambystoma mexicanum*), spotted gar (*Lepisosteus oculatus*) and red-bellied pacu (*Piaractus brachypomus*) were acquired from the pet trade and euthanized following protocols approved by the Institutional Animal Care and Use Committee at Harvard University (IACUC); tissue samples were accessioned in the Museum of Comparative Zoology (MCZ) (green anoles: R-193454, R-193455, R-193456; axolotls: 4260, 4264; spotted gar: 171800; red-bellied pacus: 171786, 171788). Tissues from crocodilians were collected as part of other studies: tissues from three American alligators (*Alligator mississippiensis*) were from R.M.C. and S.M. (subsequently accessioned into the MCZ Cryogenic collection: 4260, 4261, 4262); tissues from a Nile crocodile (*Crocodylus niloticus*) individual were obtained from N. Rattenborg and R.K.T.. Coelacanth (*Latimeria chalumnae*) tissues were from two individuals: one individual caught off Comoros in 1991 and later sampled by H. Fricke, P. Bartsch and K.H. (accessioned in Zoological Collection of the University of Tübingen; inventory number: SZ 10378); another individual caught off Tanzania in 2005 and transferred from the Tanzania Fisheries Research Institute to Tokyo Institute of Technology in 2007 for other projects (specimen number: TCC 023) (*159*, *160*). The Chinese softshell turtle (*Pelodiscus sinensis*) and two male red stingrays (*Hemitrygon akajei*) were purchased from commercial sources and tissues were collected following the guideline Rules for Animal Experiments approved by the Animal Experiment Committee of the National Institute of Genetics (Approval ID: R5-14). The oral sample for *in situ* hybridization was collected from a green anole from the animal facility of the Max Planck Institute for Biological Intelligence, Seewiesen, Germany, following the regulations of the European Union and approved protocols of the German Animal Welfare Regulation Governing Experimental Animals (TierSchVersV).

*T1R sequence curation*

We curated most taste receptor sequences from publically available vertebrate genomic and transcriptomic data (table S4), as described previously (*29*). In brief, nucleotide and amino acid sequences of *T1R*s of human, chicken, spotted gar, zebrafish and medaka as well as *T1R*s and related class C G protein-coupled receptor (*GPCR*) sequences from Atlantic stingray were used as initial queries to search for *T1R* sequences of other species genomes using NCBI tblastn applications (*161*, *162*). We constructed phylogenetic trees using candidate *T1R* sequences (thresholded with the E-value of $1x10^{-40}$ for the exon 3 and 6) and the gene sequences of the class C GPCR group 6 member A (*GPRC6A*) as the outgroup using RAxML (v8.2.12) (*163*) with the JTT + CAT model to categorize the candidate sequences into the corresponding *T1R* clades. *T1R* exon-intron boundaries were comprehensively annotated either with predictions from AUGUSTUS (v3.2.3) (*164*) or manually curated by comparing the target sequences to the queries and to *T1R*s of close relatives of the target species; splice sites were also examined. Additionally, functionally validated *T1R* sequences and cloned *T1R*s of target species were also used for subsequent iterations of the tblastn search. For the receptor naming, we used numbers to denote ancient T1R duplicates and lower case letters to refer to copies within a species (not to orthologs across different lineages). For West African lungfish *T1R5X*, “X” indicates its ambiguous phylogenetic position reported previously (*29*)).

*T1R cloning and plasmid construction*

The *T1R* sequences of green anole, American alligator, Nile crocodile, Chinese softshell turtle, axolotl, spotted gar, and red stingray were amplified with primers designed on the sequences of the target species and their close relatives (for Nile crocodile, no genome was available at the time of cloning) from cDNA libraries synthesized (SMARTer® PCR cDNA Synthesis Kit and Advantage® GC 2 Polymerase Mix & Kit (TaKaRa Bio); SuperScript III First-Strand Synthesis System and Phusion™ Hot Start II DNA Polymerases (Thermo Fisher Scientific); Tks Gflex DNA Polymerase (TaKaRa Bio)) from RNA extracted from oral tissues (RNeasy Kit and RNeasy Plus Kit (QIAGEN)). Genomic sequences of the regions surrounding the 5' and 3' ends of *T1R* coding sequences were determined using PCR with the genomic DNA (extracted using the DNeasy Blood & Tissue Kit (QIAGEN) and the NucleoSpin Tissue Kit (Macherey-Nagel)) together with species-specific primers to determine the full-length sequences of *T1R* genes. For the coelacanth, *T1R* exons were amplified from the genomic DNA extracted from muscle tissues using a phenol-chloroform extraction protocol. The amplified sequences above were cloned into an expression vector pEAK10 (Edge Biosystems) using the In-Fusion® HD Cloning Kit (TaKaRa Bio).

To obtain red-bellied pacu *T1R*s, we used a combination of a target capture sequencing approach and PCR amplification. For target capture sequencing, we first extracted DNA from the lip tissue of one individual (MagAttract HMW DNA Kit (QIAGEN)) and used the TruSeq Library Prep Kit (Illumina) to make a DNA library. We designed customized oligonucleotides based on *T1R* exonic predictions and their upstream and downstream intronic sequences of various jawed vertebrates and conducted hybridization and enrichment following the myBaits® manual (v4.01; Arbor Biosciences). Target capture fragments were pooled and sequenced on an Illumina MiSeq® with 250 bp paired-end reads and then analyzed using a custom pipeline. Using primers designed based on target capture sequencing results and *T1R* sequences of pacu close relatives, pacu *T1R* sequences were amplified from cDNA libraries generated with oral tissues (SMARTer® PCR cDNA Synthesis

Kit and Advantage® GC 2 Polymerase Mix & Kit (TaKaRa Bio); RNeasy Kit (QIAGEN)) and the full-length sequences were cloned into an expression vector pEAK10 (Edge Biosystems) following the manufacturer's protocol (In-Fusion® HD Cloning Kit (TaKaRa Bio)).

The curated sequences of African bullfrog (*Pyxicephalus adspersus*), guppy (*Poecilia reticulata*), white shark (*Carcharodon carcharias*), West African lungfish, reedfish (*Erpetoichthys calabaricus*) and bowfin (*Amia calva*) were synthesized (Azenta Life Sciences) and subcloned into the expression vector (using In-Fusion® HD Cloning kit (TaKaRa Bio) or by Azenta Life Sciences). All plasmids were verified using Sanger sequencing.

*Amniote T1R selection tests*

To test if we could detect signals of relaxed selection on crocodilian *T1R2*, we aligned six crocodilian *T1R2* genes with a set of intact and pseudogenized *T1R2*s from other amniotes, including 35 mammals (21 intact genes and 14 pseudogenes), five lepidosaurians (five intact genes) and five turtles (four intact genes and one pseudogene) extracted from a previous study (*30*). Coding sequences were translated using EMBOSS transeq (*165*), by first removing frameshifts and internal stop codons from pseudogenes. Protein sequences were then aligned using MUSCLE (v5) (*166*) and backtranslated to a coding sequence alignment using trimAl "-backtrans". This alignment and the species tree were used as input in RELAX with the synonymous rate variation option (*56*) to assess if crocodilian branches had a different selection regime than intact genes from other amniotes. RELAX uses likelihood ratio tests (LRT) to compare between a null model, under which background branches (i.e. intact amniote *T1R2* branches) and test branches (i.e. crocodilian branches) have the same dN/dS (ω, the ratio of non-synonymous to synonymous substitutions) distribution, and a model under which a parameter K is introduced to test whether selection has been relaxed or intensified along the test branches ($K < 1$, relaxed selection; $K > 1$, intensified selection; in the null model, $K = 1$). We considered RELAX results to be significant if the LRT $p$-value was $< 0.05$. In addition, to ensure convergence of results, RELAX was run ten times independently. The same alignment procedure and RELAX analysis were applied to *T1R1* and *T1R3* genes separately, with the addition of 34 avian sequences.

We then assessed if the relaxation signal we observed with RELAX on crocodilian *T1R2* genes could be indicative of complete relaxation of selection (i.e., neutral evolution) by using two independent approaches. First, using the same codon alignment as above and the PAML package (*167*), we compared three nested branch models: (i) a single ω shared across all branches; (ii) two ω values, one for intact genes and one shared by crocodilian genes and pseudogenes; and (iii) three ω values, one for intact genes, one for crocodilian genes, and one for pseudogenes. Pairs of models were compared using LRTs. As a second approach, we tested whether the pattern of amino acid substitution accumulation in crocodilian *T1R2* genes was more consistent with functional constraint or neutral evolution. Amino acid substitutions were reconstructed across branches of the species tree using phangorn (LG substitution model, four gamma rate categories) (*168*). The predicted deleteriousness of each substitution was scored using MutPred2 (MP2) (*169*), which assigns values from 0 (non-deleterious) to 1 (highly deleterious). We compared MP2 score distributions across four groups: (i) the crocodilian most recent common ancestor (MRCA) branch, (ii) extant crocodilian terminal branches, (iii) ten intact amniote *T1R2* genes, and (iv) ten amniote *T1R2* pseudogenes. As a neutral reference, we performed 100 neutral evolutionary simulations in which the human *TAS1R2* coding sequence evolved without selection, using a transition/transversion ratio of 2.59 (equal to the observed ratio on the *T1R2* alignment), until accumulating a number of amino acid substitutions equal to the mean that was observed per branch in the empirical dataset (65 substitutions). Simulated sequences were also scored with MP2. All pairwise comparisons of MP2 score distributions were performed using Kolmogorov–Smirnov tests, with *p*-values corrected for multiple testing using the false discovery rate (FDR) method.

*Fish T1R phylogenetic analysis and ancestral reconstruction*

To infer ancestral T1R2B-T1R3B receptor sequences within ray-finned fish, we first aligned 133 full-length *T1R2* and 55 *T1R3* nucleotide sequences of 54 ray-finned fishes, nine lobe-finned fishes as well as *T1R6*s and *T1R3C* of Australian ghostshark (*Callorhinchus milii*) as outgroups separately using the TranslatorX webserver (*170*) with the MAFFT alignment algorithm. We used PhyML Smart Model Selection (*171*) to identify GTR+G+I model as the optimal model, which was used for phylogenetic tree inference in RAxML (v8.2.12) (*163*) (100 replicates were performed). Inferred trees were modified based on known species relationships (*71*, *149*, *172*) and branches were collapsed into a polytomy in Mesquite (v4.0.1) (*173*) if the bootstrap support values < 50. We then performed codon-based marginal reconstructions separately using *T1R2* and *T1R3* nucleotide alignments and their corresponding tree topologies with the Yang codon model in FastML (*174*).

Six inferred ancestral *T1R2B* and *T1R3B* sequences (from the ancestors of all ray-finned fishes, of bowfin, gars and teleosts, and of all teleosts) were manually checked for insertions and deletions by aligning with wild-type sequences before codon optimization for expression in HEK293T cells (GenSmart™ codon optimization, GenScript) and synthesis into pEAK10 vectors (Azenta Life Sciences).

*Jawed vertebrate T1R phylogenetic analysis*

For ancestral reconstruction within jawed vertebrates, we first inferred phylogenetic trees of *T1R*s of jawed vertebrates using two sets of full-length *T1R* amino acid sequences (a full dataset: 190 *T1R*s of 44 species from 11 *T1R* clades; a reduced dataset: 185 *T1R*s of 51 species from eight clades; described below) with seven outgroup *GPRC6A*s (the closest relatives of *T1R*s (*78*)). These two sequence sets were aligned using MAFFT (v7.427) (*175*) with the ginsi option (*175*) followed by manual curation, and phylogenetic trees were constructed using RAxML (v8.2.12) (*163*) with the JTT + CAT model and 1,000 bootstrap replicates.

*Jawed vertebrate T1R ancestral reconstruction*

To assess whether the difference in sequence selection might affect ancestral receptor functions across jawed vertebrates, we inferred ancestral sequences using FastML (v3.1) (*174*) with the JTT + G model, using the two sets of *T1R* amino acid alignments and topologies as inputs.

First, we evaluated the influence of the number of *T1R* clades included in the sequence alignment by comparing ancestral receptor functions inferred from two alignment sets and their corresponding species tree topologies. Compared to the reduced dataset (185 *T1R*s and seven *GPRC6A*s), the full dataset included 14 recently-described non-canonical *T1R*s (*T1R5*s of coelacanth, lungfishes and amphibians; *T1R7*s of axolotl and lizards; *T1R4*s of bony vertebrates; *T1R8*s of bichir and West African lungfish; in total: 190 *T1R*s and seven *GPRC6A*s) (see fig. S16 and S20). To comply with the FastML sequence limit (maximum 200 taxa), we excluded sequences of eight recently-diverged bony vertebrate species or species without a full receptor repertoire in the full dataset (i.e. *Erinaceus europaeus*, *Salvator merianae*, *Podocnemis expansa*, *Spea multiplicata*, *Oophaga pumilio*, *Neoceratodus forsteri*, *Pygocentrus nattereri* and *Atractosteus spatula*), as receptors of closely related species were already included (therefore, excluding sequences of these species would be unlikely to affect the ancestral reconstruction of the deeper nodes), and added receptors of *Pelodiscus sinensis* (a species for which we generated functional data) and *Scyliorhinus torazame* (to increase the representation of cartilaginous fish) in the full dataset. For the species tree topologies, we modified the maximum-likelihood trees constructed with the full dataset and with the reduced dataset corresponding to known species relationships (*145*, *176–180*). Due to low bootstrap support values (see also (*29*, *30*)), we depicted *T1R1*, *T1R2* and *T1R2B* as a polytomy using Mesquite (v4.0.1) (*173*) (fig. S17 and S21).

We reconstructed and functionally tested ancestral heterodimers inferred from six phylogenetic junctions (the ancestors of all jawed vertebrates, all bony vertebrates, all lobe-finned fish, all tetrapods, all amniotes and all mammals) from the full and reduced dataset separately. We found that the ancestral heterodimers of all jawed vertebrates and all bony vertebrates inferred from both sets (the species tree with the full alignment, and the species tree with the reduced alignment) showed consistent sucrose responses, suggesting a very early origin of sugar sensing in vertebrates (Fig. 4C; fig. S22A; the ancestral receptors of all T1Rs were also inferred from both sets, but we did not observe any response elicited by these two homodimers to 17 amino acids or two sugars or sucralose). We also observed heightened specialized responses to amino acids in the ancestral T1R1-T1R3 and to sucrose in the T1R2-T1R3 of all mammals inferred from both sets (Fig. 4E-G; fig. S22A). The ancestral T1R2-T1R3 pairs of the lobe-finned fish and amniote ancestors (from both sets) showed small responses to sugars and had no amino acid responses. Whereas the ancestral T1R2-T1R3 from all tetrapods inferred from the full set responded to three sugars (with no amino acid response), this was not the case for the tetrapod ancestral pair inferred from the reduced set, which did not respond to any of the six ligands tested, suggesting the possibility that this receptor pair (inferred from the reduced set) may not be fully functional in the heterologous system (Fig. 4E; fig. S18 and S22B-C). We additionally inferred and tested the ancestral T1R1-T1R3 and T1R2-T1R3 from the ancestor of all reptiles (using the reduced alignment), which demonstrated that sucrose responses are present in ancestral receptor pairs (fig. S22A-B), consistent with those observed in the receptor pairs of extant non-avian reptiles (Fig. 2C). In summary, although sequence selection in general plays a role in ancestral sequence reconstruction, the reconstructions from the two sets of alignments and their respective topologies in our study did not result in major differences in receptor responses.

To further compare our ancestral receptor responses with those reported by Feng *et al.* (*27*), we synthesized and functionally examined the three T1R2-T1R3 receptor pairs reported in that study (from the ancestors of all tetrapods, all amniotes and all reptiles (sauropsids)) inferred using two alignments of 34 *T1R2* and 38 *T1R3* amino acid sequences and the species tree topologies. Among these ancestral pairs (inferred by Feng *et al.* (*27*)), we detected responses to three sugars, sucralose and three amino acids by T1R2-T1R3 from the ancestor of all reptiles and responses to sucralose and two amino acids by the T1R2-T1R3 pair from the ancestor of all tetrapods (fig. S22B). For the ancestral T1R2-T1R3 of all amniotes, we did not observe any response to six ligands tested in the pair from (*27*). Feng *et al.* did not observe any ancestral receptor response to three sugars (*27*), and this discrepancy between our results and the reported lack of responses may be due to technical issues, as amino acid responses (as control) had not been previously examined.

All ancestral sequences (marginal reconstructions) inferred in this study were manually checked for insertions and deletions using the alignment of extant sequences. The amino acid sequences of vertebrate ancestral *T1R*s from our study and from a previous study (*27*) were back-translated and codon-optimized (GenScript) before direct synthesis into the pEAK10 vectors (Azenta Life Sciences).

*Functional cell assays*

To functionally characterize extant and ancestral T1R receptors, we used an established luminescence-based cell assay (*54*, *55*). We transiently transfected human HEK293T cells (from the Matsunami Laboratory, Duke University, USA) with plasmids of target T1Rs, a rat G-protein (Gα15i2) and a calcium-dependent photoprotein (mt-apoclytin-II) using the Lipofectamine 2000 reagent (Invitrogen). Transfected cells were cultured for 48 hours before measuring their ligand-evoked responses on 96-well CellBIND surface plates (Corning) in a FlexStation 3 microplate reader (Molecular Devices). After removing the media, transfected cells were loaded with coelenterazine (Promega) and the loading buffer (10 mM 4-(2-hydroxyethyl)-1-piperazineethanesulfonic acid (HEPES), 130 mM NaCl, 10 mM glucose, 5 mM KCl, 2 mM $CaCl_2$, and 1.2 mM $MgCl_2$, 0.1% bovine serum albumin (BSA) (Sigma-Aldrich); pH 7.4) and then incubated at 27°C in the dark for 4 hours. Upon ligand solution addition, the luminescence intensity over time in each well was measured and the area under the curve (AUC) was calculated as the receptor activity (relative light units, RLU). The control cell response was also measured by the ligand response of cells transfected with the G-protein and photoprotein but not transfected with receptor plasmids. The baseline activity of the transfected (non-control) cells was defined by the response to the loading buffer without any ligand addition (baseline response).

For dose-dependent response data, concentrations of sucrose, fructose, sucralose and alanine were tested at 100, 50, 25, 12.5, 6.25, 3.125, 1.562, 0.391 mM and those of arginine and proline were at 50, 25, 12.5, 6.25, 3.125, 1.562, 0.781, 0.195 mM. To detect the dose-dependent changes of the red stingray T1R6.1-T1R3C to proline, this pair was tested with 18 additional lower concentrations of proline solutions (0.39, 0.098, 0.049, 0.024, 0.012, 0.006, 0.003, 0.0015, 0.0007, 0.00038, 0.00019, 0.0001, 0.00005, 0.000023, 0.000012, 0.000006, 0.000003, 0.000001 mM). Five ancestral T1R1-T1R3 pairs within bony vertebrates inferred from the full dataset were tested with 4 additional low concentrations (0.049, 0.012, 0.0031, 0.00019 mM) of arginine and proline to examine whether there was a significant decrease of responses to lower concentrations. For the ligand panel tested in fig.

S4B, S5, S7C, S8C, S15B and S22A-B, sugars and sucralose were tested at 100 mM and amino acids at 50 mM (Sigma-Aldrich).

For dose-dependent response data, we applied one-tailed Welch's *t*-tests followed by FDR (via the Benjamini-Hochberg procedure (BH) to correct for multiple ligand testing for each receptor pair) to compare the response elicited by the highest concentration to that by the lowest concentration for each ligand (provided that the response to the highest concentration was larger than the control cell response or baseline response to confirm that the observed luminescience changes were not caused by non-specific responses). For receptor pairs with the highest response elicited not by the highest concentration of a ligand (i.e. bowfin T1R1-T1R3B and T1R2-T1R3B, spotted gar T1R2Bb-T1R3B, guppy T1R2Bd-T1R3B, medaka T1R2Ba-T1R3B and T1R2Bb-T1R3B, and red stingray T1R6.1-T1R3C), we conducted additional one-tailed Welch's *t*-tests and BH corrections to compare the highest response to any concentration of a ligand to the lowest mean response to any concentration of any ligand in this receptor pair (and also confirmed that that the highest response was also larger than control cell or baseline response). Our results suggest that some ligands (e.g. sucralose) may act as inverse agonists for certain receptors (e.g. bowfin T1R2-T1R3B, guppy T1R2Bd-T1R3B and red stingray T1R6.1-T1R3C) or alternatively, some of these receptor pairs might be initially activated by lower concentrations than what we have measured and decreased within the eight-point concentration series used for most receptors (similar to the observations of the human sweet receptor to several artificial sweeteners above 10 mM (*181–183*)), which remains to be verified in future studies. For data presented in fig. S4B, S5, S7C, S8C, S15B and S22A-B, we used one-tailed Welch's *t*-tests and BH multiple testing corrections to compare the ligand-evoked responses to the no ligand baseline response (and confirmed that the receptor ligand response was larger than the control cell response or baseline response). It is important to note that when we refer to a receptor as nonfunctional (inferred from the non-significant response to canonical sugars and amino acids tested in this study), this may not imply a true absence of response in a receptor pair (as it could reflect a technical issue, for instance related to poor expression in the heterologous system).

To assess the ancestral receptor's relative response sensitivity to sucrose and alanine, we used the dose-dependent response data to calculate a response index for each receptor (the difference of average responses to 100 mM sucrose and 100 mM alanine divided by the sum of average responses to sucrose and alanine). Cell assay data were analyzed and visualized in R (v4.5.0) (*157*).

*In situ hybridization*

Five *T1R*, phospholipase C beta 2 (*PLCβ2*) and transient receptor potential cation channel subfamily M member 5 (*TRPM5*) genes of green anoles were used to design digoxigenin-labeled probes and synthesized for single-color *in situ* hybridization. One anole tongue tissue embedded in the optimal cutting temperature compound was sectioned sagittally (10 μm thick) and placed on MAS-coated glass slides (Matsunami Glass). We performed *in situ* hybridization as described previously (*184*, *185*). In short, after fixation with 4% paraformaldehyde in phosphate-buffered saline, the sections were prehybridized (58 °C, 1 h), hybridized (58 °C, two overnight sessions), washed (58 °C, 0.2× saline–sodium citrate) and developed with a single probe (nitroblue tetrazolium/ 5-bromo-4-chloro-3-indolyl phosphate; NBT-BCIP). Stained sections were imaged using a fluorescence microscope (DM6 B; Leica) equipped with a cooled CCD digital camera (DFC7000 T; Leica).

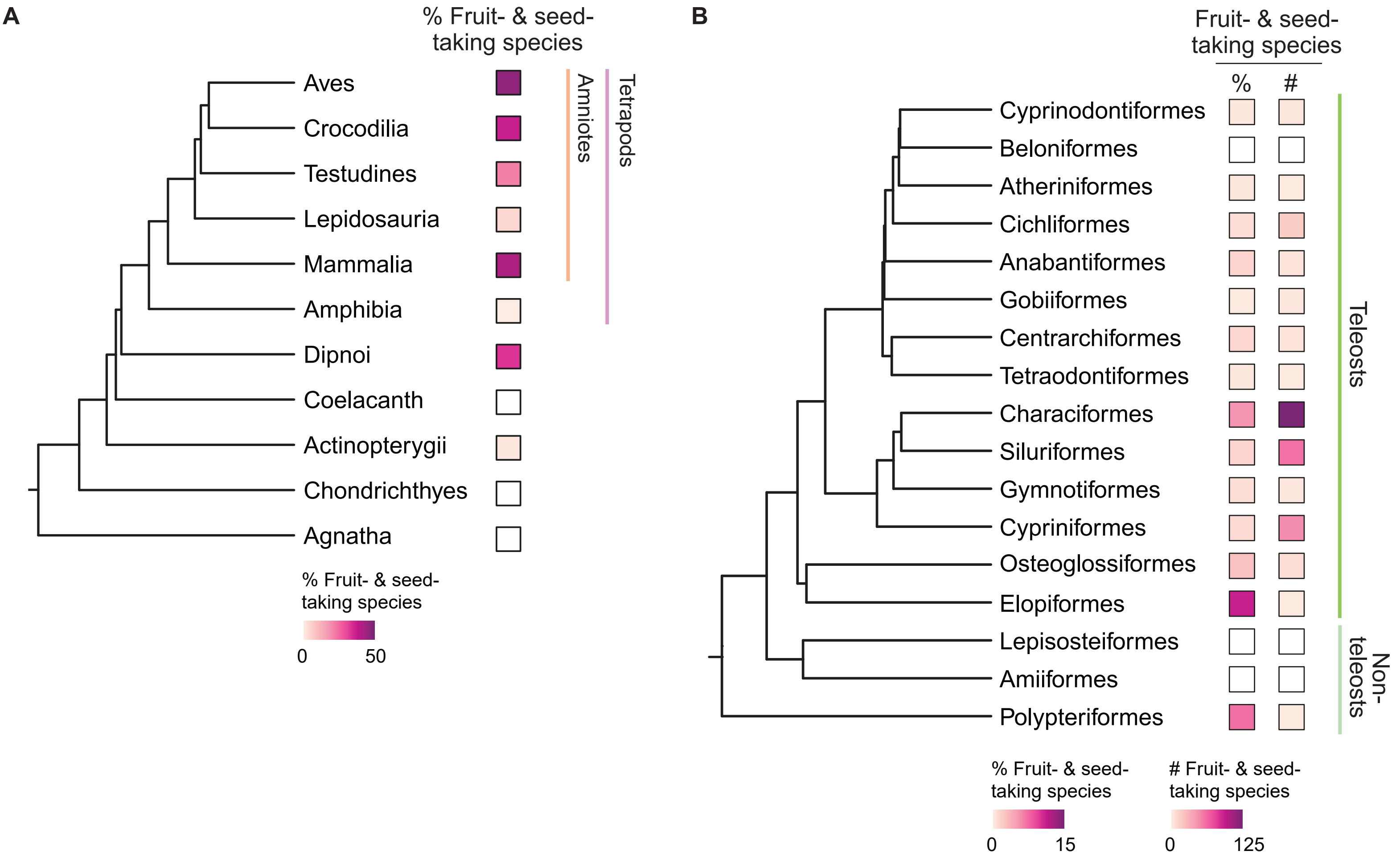
A
% Fruit- & seed-taking species
Aves
Crocodilia
Testudines
Lepidosauria
Mammalia
Amphibia
Dipnoi
Coelacanth
Actinopterygii
Chondrichthyes
Agnatha
Amniotes
Tetrapods
% Fruit- & seed-taking species
0
50
B
Fruit- & seed-taking species
%
#
Cyprinodontiformes
Beloniformes
Atheriniformes
Cichliformes
Anabantiformes
Gobiiformes
Centrarchiformes
Tetraodontiformes
Characiformes
Siluriformes
Gymnotiformes
Cypriniformes
Osteoglossiformes
Elopiformes
Lepisosteiformes
Amiiformes
Polypteriformes
Teleosts
Non-teleosts
% Fruit- & seed-taking species
0
15
### Fruit- & seed-taking species
0
125

**fig. S1. Fruit- and seed-taking species are widespread across vertebrates.** (**A**) Fruit- and seed-taking are present across eight major clades of vertebrates (%: number of fruit- & seed-taking species (pink) divided by total species count per clade). (**B**) Fruits and seeds are consumed across 14 orders of ray-finned fish (274 of 2,564 species exampled; 14 of 84 orders; #: number of species; %: number of fruit- & seed-taking species (pink) divided by total species count per order). Related to Fig. 1A-B and 3A; table S1-2.

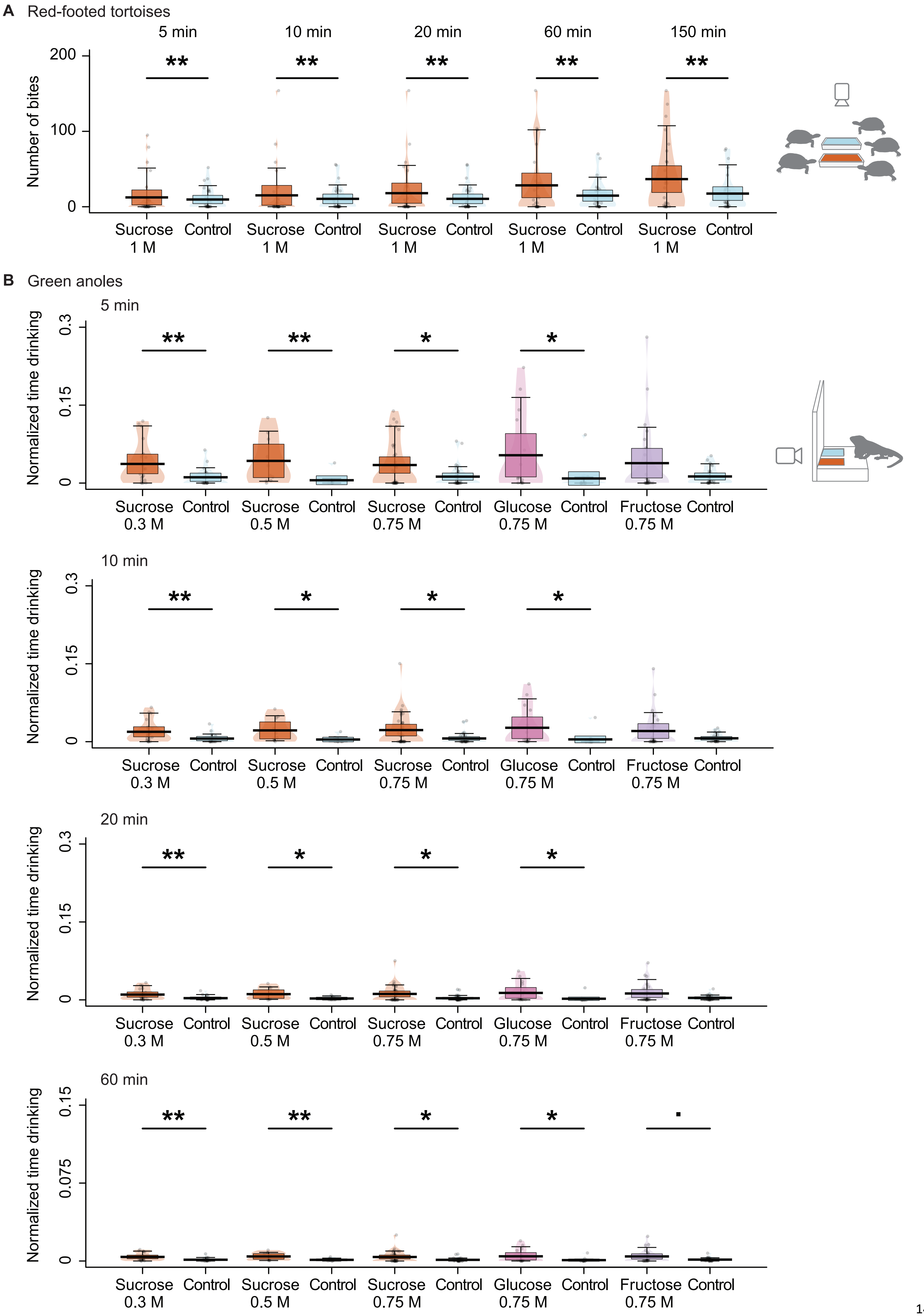
A Red-footed tortoises
5 min
10 min
20 min
60 min
150 min
Number of bites
200
100
0
**
Sucrose 1 M
Control
B Green anoles
5 min
10 min
20 min
60 min
Normalized time drinking
0.3
0.15
0.075
0
*
·
Sucrose 0.3 M
Sucrose 0.5 M
Sucrose 0.75 M
Glucose 0.75 M
Fructose 0.75 M
Control

**fig. S2. Two non-avian reptiles show behavioral preferences for sugar.** (**A**) Red-footed tortoises (*Chelonoidis carbonarius*) were significantly attracted to agar cubes containing 1 M sucrose (orange) over plain agar cubes with water as control (blue) in the 5 minutes after the first bite; this preference remained over the entire trial (up to 150 minutes). Five animals were tested and filmed as a group. Generalized linear-mixed models with a Poisson distribution; ** $p \leq 0.01$; 7 trials, 5 individuals. (**B**) Green anoles (*Anolis carolinensis*) significantly preferred 0.3 M, 0. 5 M and 0.75 M sucrose solutions (orange) as well as 0.75 M glucose solution (pink) over water control (blue) in a 5-minute time window after the first drinking bout. These preferences remained consistent over the entire hour (60 minutes). A trend ($p = 0.051$; not significant) towards consumption of 0.75 M fructose (purple) preference was only observed after the 60-minute exposure. Linear-mixed models; · $0.05 < p \leq 0.1$, * $0.01 < p \leq 0.05$, ** $p \leq 0.01$; 0.3 M sucrose, 20 trials, 3 individuals; 0.5 M sucrose, 10 trials, 2 individuals; 0.75 M sucrose, 34 trials, 6 individuals; 0.75 M glucose, 15 trials, 3 individuals; 0.75 M fructose, 25 trials, 4 individuals. Related to Fig. 1C-D; table S3.

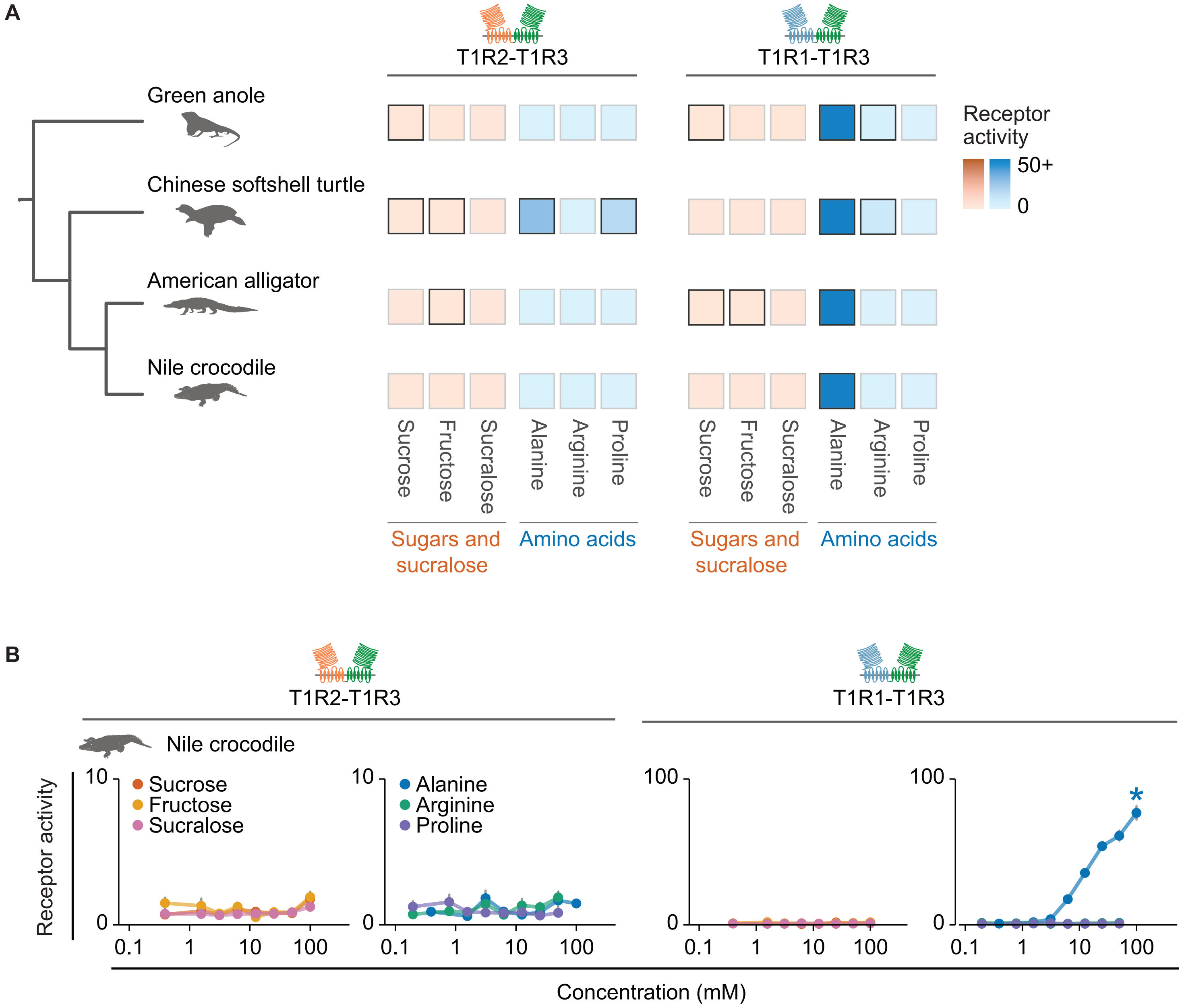
A
T1R2-T1R3
T1R1-T1R3
Green anole
Chinese softshell turtle
American alligator
Nile crocodile
Receptor activity
50+
0
Sucrose
Fructose
Sucralose
Alanine
Arginine
Proline
Sugars and sucralose
Amino acids
B
T1R2-T1R3
T1R1-T1R3
Nile crocodile
Receptor activity
Sucrose
Fructose
Sucralose
Alanine
Arginine
Proline
Concentration (mM)

**fig. S3. Summary and responses of non-avian reptile T1R2-T1R3 and T1R1-T1R3 receptor pairs.** Summary (**A**) of dose-dependent responses of T1R2-T1R3 and T1R1-T1R3 of four non-avian reptiles (Fig. 2C) and (**B**) dose-dependent responses of Nile crocodile (*Crocodylus niloticus*) receptor pairs. Receptor activity is measured as the area under the curve (AUC) (x$10^3$ relative light units, RLU) (n= 5-9; mean ± s.e.m; significance (* $p < 0.05$) assessed with one-tailed Welch's *t*-tests (with the Benjamini-Hochberg (BH) multiple testing corrections) comparing responses to the highest and lowest concentrations (provided that the response to the highest concentrations was larger than control cell response (cells transfected with G-protein and photoprotein but without the receptors) or baseline response). Colored cells in the heatmap (A) show the response of transfected cells to the highest concentration of each ligand (dark gray outline: significant response from dose-dependent response data in Fig. 2C and fig. S3B; light gray outline: no significant response). Concentrations tested were 100, 50, 25, 12.5, 6.25, 3.125, 1.562, 0.391 mM for sucrose, fructose, sucralose and alanine and 50, 25, 12.5, 6.25, 3.125, 1.562, 0.781, 0.195 mM for arginine and proline.

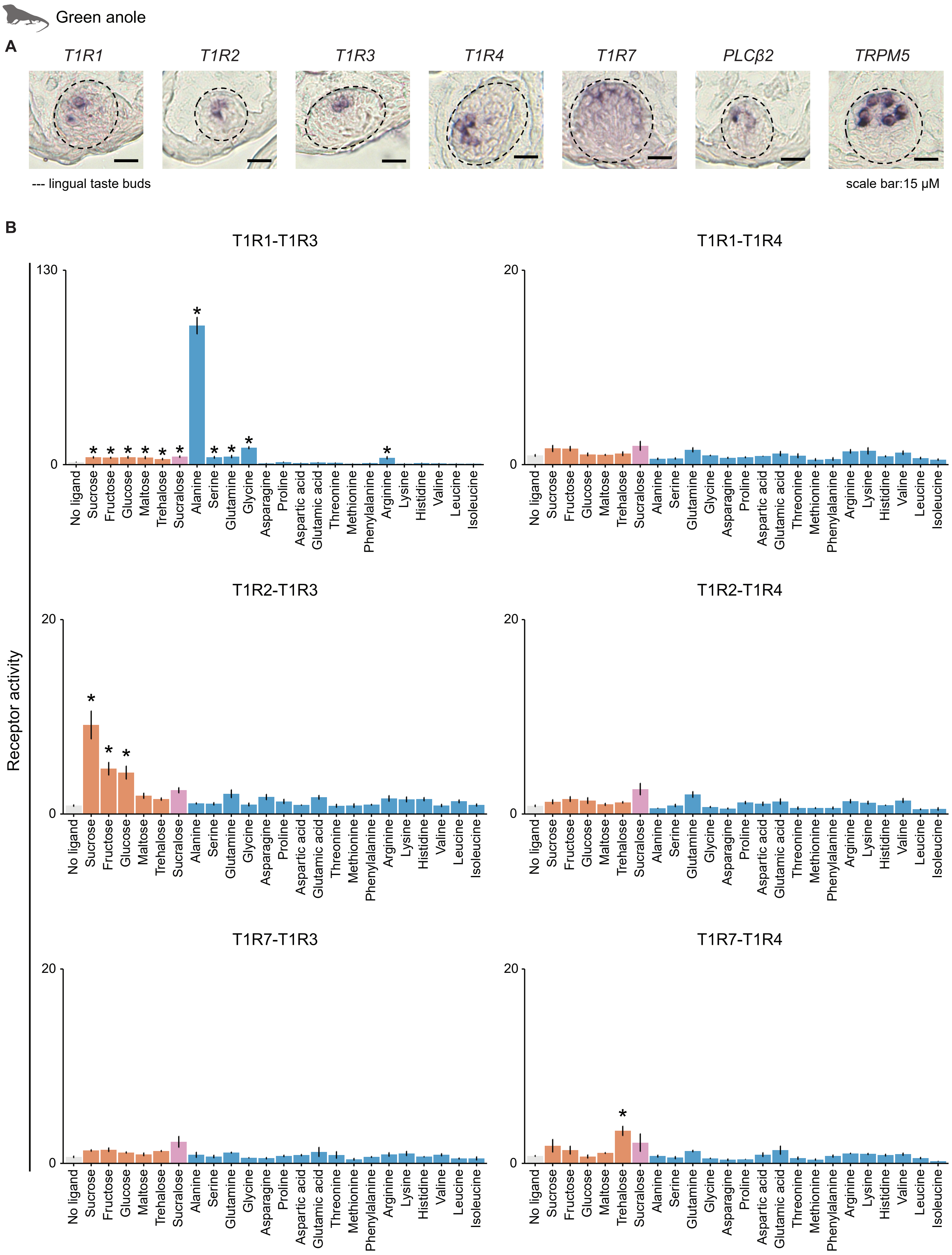
Green anole
A
T1R1
T1R2
T1R3
T1R4
T1R7
PLCβ2
TRPM5
--- lingual taste buds
scale bar:15 μM
B
Receptor activity
T1R1-T1R3
130
0
T1R1-T1R4
20
0
T1R2-T1R3
20
0
T1R2-T1R4
20
0
T1R7-T1R3
20
0
T1R7-T1R4
20
0
No ligand
Sucrose
Fructose
Glucose
Maltose
Trehalose
Sucralose
Alanine
Serine
Glutamine
Glycine
Asparagine
Proline
Aspartic acid
Glutamic acid
Threonine
Methionine
Phenylalanine
Arginine
Lysine
Histidine
Valine
Leucine
Isoleucine

**fig. S4. Lingual expression of anole *T1R*s and their downstream effector genes as well as extended ligand panel responses of six anole receptor pairs.** (**A**) Five *T1R* genes and two taste bud marker genes (phospholipase C beta 2 (*PLCβ2*) and transient receptor potential cation channel subfamily M member 5 (*TRPM5*)) are expressed in putative taste buds in sagittal sections of the anole tongue. Dashed circles, lingual taste buds; scale bar, 15 μm. (**B**) Functional characterizations of six green anole receptor pairs reveal that its T1R1-T1R3 is primarily an amino acid sensor (which retains responses to various sugars and sucralose), whereas T1R2-T1R3 has responses specifically to sugars and sucralose. Anole T1R7-T1R4 appears to only respond weakly to trehalose and three other heterodimers T1R1-T1R4, T1R2-T1R4 and T1R7-T1R3 did not respond to five sugars, sucralose or 17 amino acids tested. Receptor activity is calculated as AUC (x$10^3$ RLU; mean ± s.e.m; n = 3-11; significance (* $p < 0.05$) assessed with one-tailed Welch's *t*-tests and BH corrections comparing any ligand response to the no ligand baseline response (provided that the mean ligand response was larger than the control cell response or the baseline response). Sugars and sucralose were tested at 100 mM and amino acids at 50 mM.

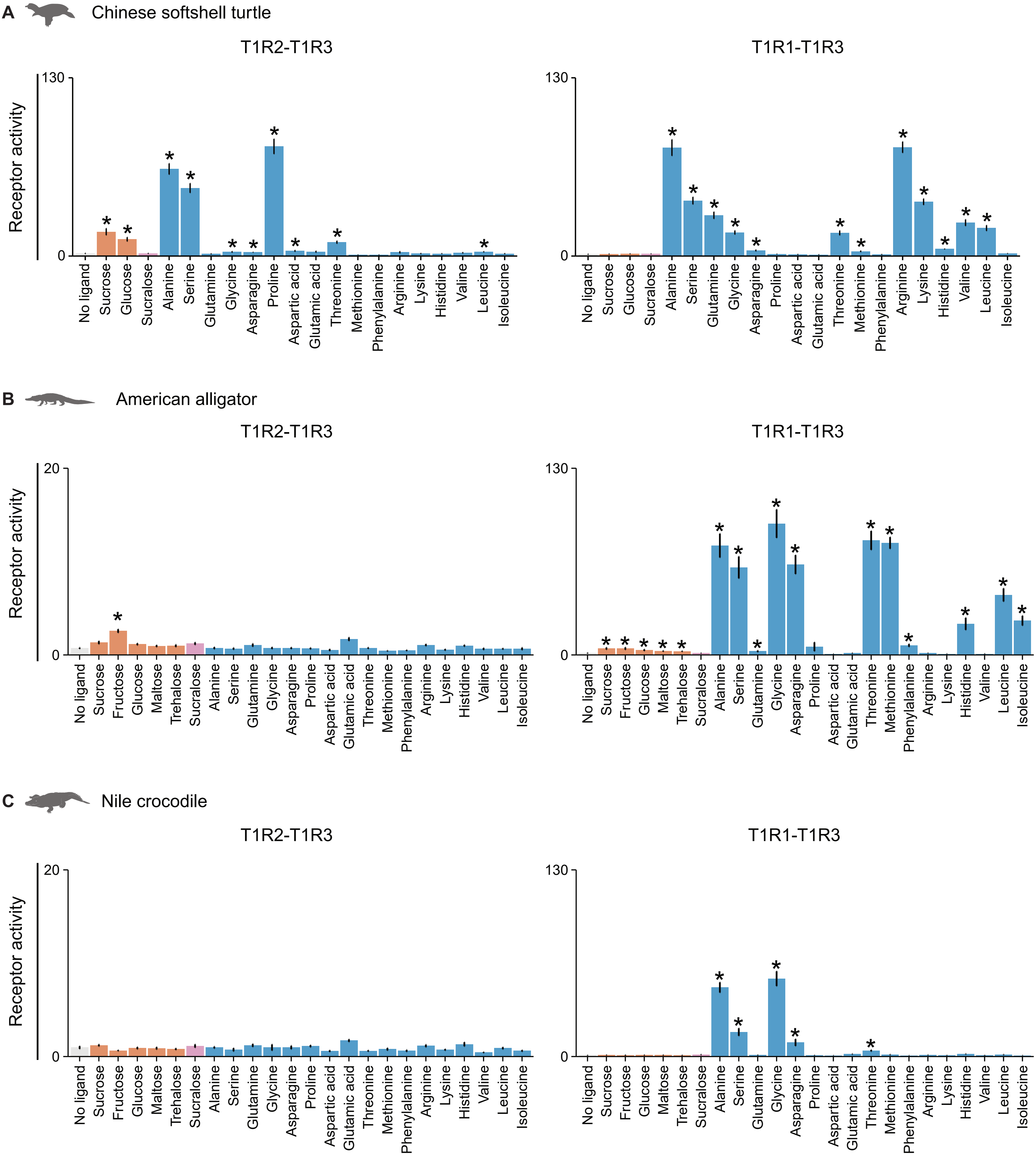
A
Chinese softshell turtle
T1R2-T1R3
T1R1-T1R3
Receptor activity
130
0
No ligand
Sucrose
Glucose
Sucralose
Alanine
Serine
Glutamine
Glycine
Asparagine
Proline
Aspartic acid
Glutamic acid
Threonine
Methionine
Phenylalanine
Arginine
Lysine
Histidine
Valine
Leucine
Isoleucine
B
American alligator
T1R2-T1R3
T1R1-T1R3
Receptor activity
20
130
0
No ligand
Sucrose
Fructose
Glucose
Maltose
Trehalose
Sucralose
Alanine
Serine
Glutamine
Glycine
Asparagine
Proline
Aspartic acid
Glutamic acid
Threonine
Methionine
Phenylalanine
Arginine
Lysine
Histidine
Valine
Leucine
Isoleucine
C
Nile crocodile
T1R2-T1R3
T1R1-T1R3
Receptor activity
20
130
0

**fig. S5. Extended ligand panel responses of turtle and crocodilian receptor pairs.** (**A**) Both Chinese softshell turtle T1R2-T1R3 and T1R1-T1R3 respond strongly to various amino acids but T1R2-T1R3 also shows responses to sucrose and glucose. (**B**) American alligator has a fructose-sensing T1R2-T1R3 pair, whereas its T1R1-T1R3 pair is primarily an amino acid sensor with responses to various sugars and sucralose. (**C**) Nile crocodile T1R2-T1R3 appeared nonfunctional and its T1R1-T1R3 only responds to amino acids. Receptor activity is calculated as AUC ($x10^3$ RLU; mean ± s.e.m; n = 5-12; significance (* $p < 0.05$) assessed with one-tailed Welch's *t*-tests and BH corrections as in fig. S4; similar ligand concentrations as fig. S4).

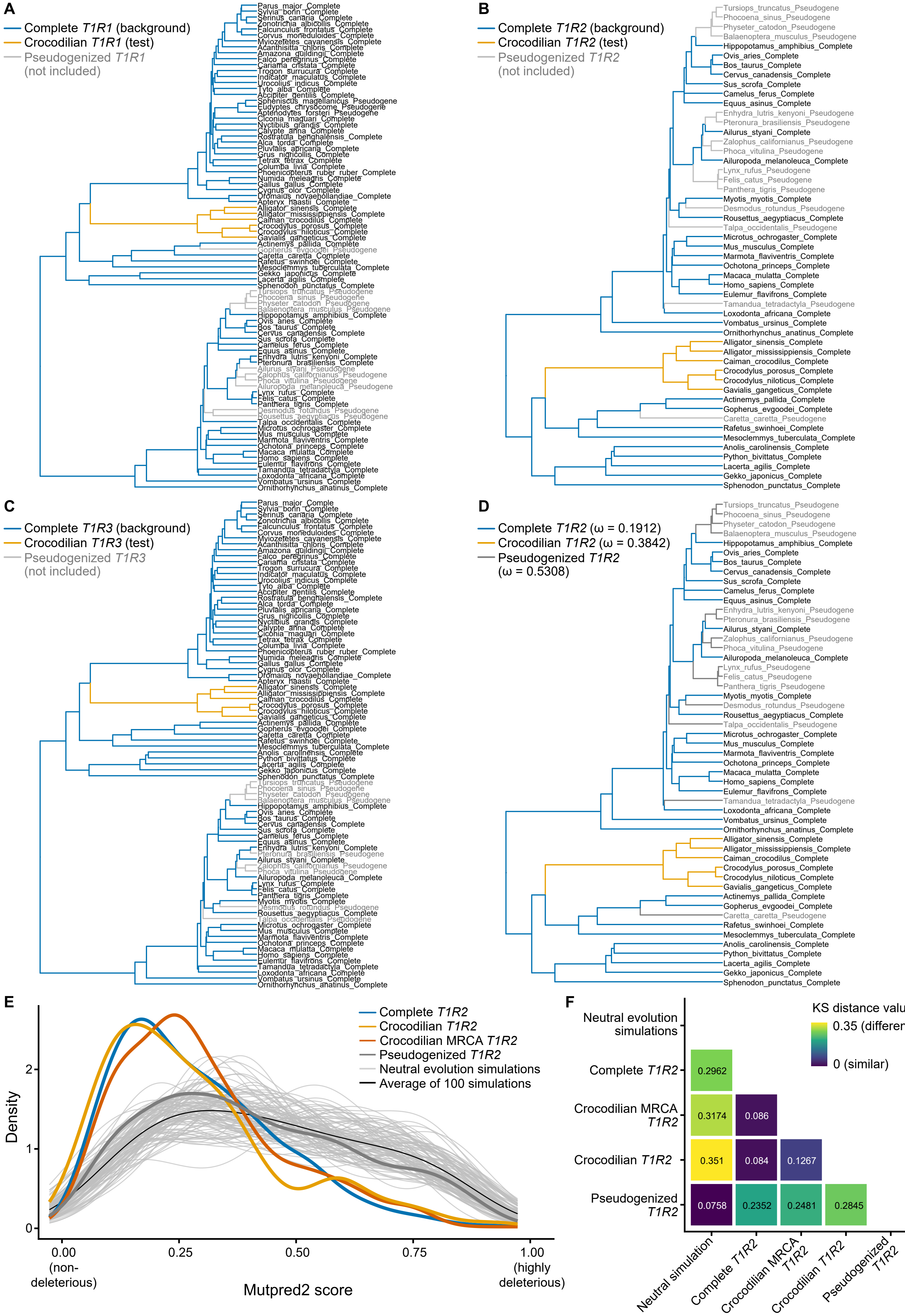
A
Complete T1R1 (background)
Crocodilian T1R1 (test)
Pseudogenized T1R1
(not included)
B
Complete T1R2 (background)
Crocodilian T1R2 (test)
Pseudogenized T1R2
(not included)
C
Complete T1R3 (background)
Crocodilian T1R3 (test)
Pseudogenized T1R3
(not included)
D
Complete T1R2 (ω = 0.1912)
Crocodilian T1R2 (ω = 0.3842)
Pseudogenized T1R2
(ω = 0.5308)
E
Complete T1R2
Crocodilian T1R2
Crocodilian MRCA T1R2
Pseudogenized T1R2
Neutral evolution simulations
Average of 100 simulations
Density
0
1
2
0.00
(non-
deleterious)
0.25
0.50
0.75
1.00
(highly
deleterious)
Mutpred2 score
F
KS distance values
0.35 (different)
0 (similar)
Neutral evolution
simulations
Complete T1R2
Crocodilian MRCA
T1R2
Crocodilian T1R2
Pseudogenized
T1R2
0.2962
0.3174
0.086
0.351
0.084
0.1267
0.0758
0.2352
0.2481
0.2845
Neutral simulation
Complete T1R2
Crocodilian MRCA
T1R2
Crocodilian T1R2
Pseudogenized
T1R2

**fig. S6. Crocodilian *T1R2*—but not *T1R1* or *T1R3*—is under relaxed selection but distinct from *T1R2* pseudogenes.** (**A-C**) Six crocodilian (test branches; orange) *T1R2* genes (B) likely have evolved under relaxed selection, whereas their *T1R1* (A) or *T1R3* (C) show no significant shift in selection intensity relative to complete genes of non-crocodilian amniotes (background branches; blue) (see Methods). Pseudogenes (gray) were not included in the relaxed selection tests. (**D**) PAML hypothesis testing reveals that crocodilian *T1R2*s have a significantly higher dN/dS ratio (ω, the ratio of non-synonymous to synonymous substitutions) than complete genes, coherent with the relaxation of selection detected with RELAX, but a significantly lower ω than pseudogenized *T1R2*s, indicating that they are not evolving under neutral evolution. (**E-F**) Distributions of Mutpred2 scores (from 0 to 1: non-deleterious to highly deleterious mutations) in (i) the crocodilian most recent common ancestor (MRCA, dark orange), (ii) six extant crocodilian *T1R2*s (light orange), (iii) ten complete amniote *T1R2*s (blue), (iv) ten amniote pseudogenes (dark gray) and (v) 100 neutral evolution simulations of the human *TAS1R2* gene (light gray, with their average in black). The distributions of extant crocodilian sequences or of their MRCA are similar to the distribution of complete *T1R2*s (assessed using Kolmogorov-Smirnov tests (KS); KS distance values in (F); purple and dark blue: similar distributions with relatively small KS values; green to yellow: dissimilar distributions with relatively large KS values), with an accumulation of non-deleterious mutations and few deleterious mutations. These distributions are very dissimilar to the observed or expected distributions under neutral evolution (distributions of pseudogenes and simulations, respectively), which are more uniform, with an increased accumulation of deleterious mutations. Therefore, crocodilian *T1R2* is under relaxed selection but to a lesser extent observed in pseudogenized *T1R2*. Neutral evolution simulations were performed using a transition/transversion ratio of 2.59, equal to the observed ratio on the *T1R2* alignment.

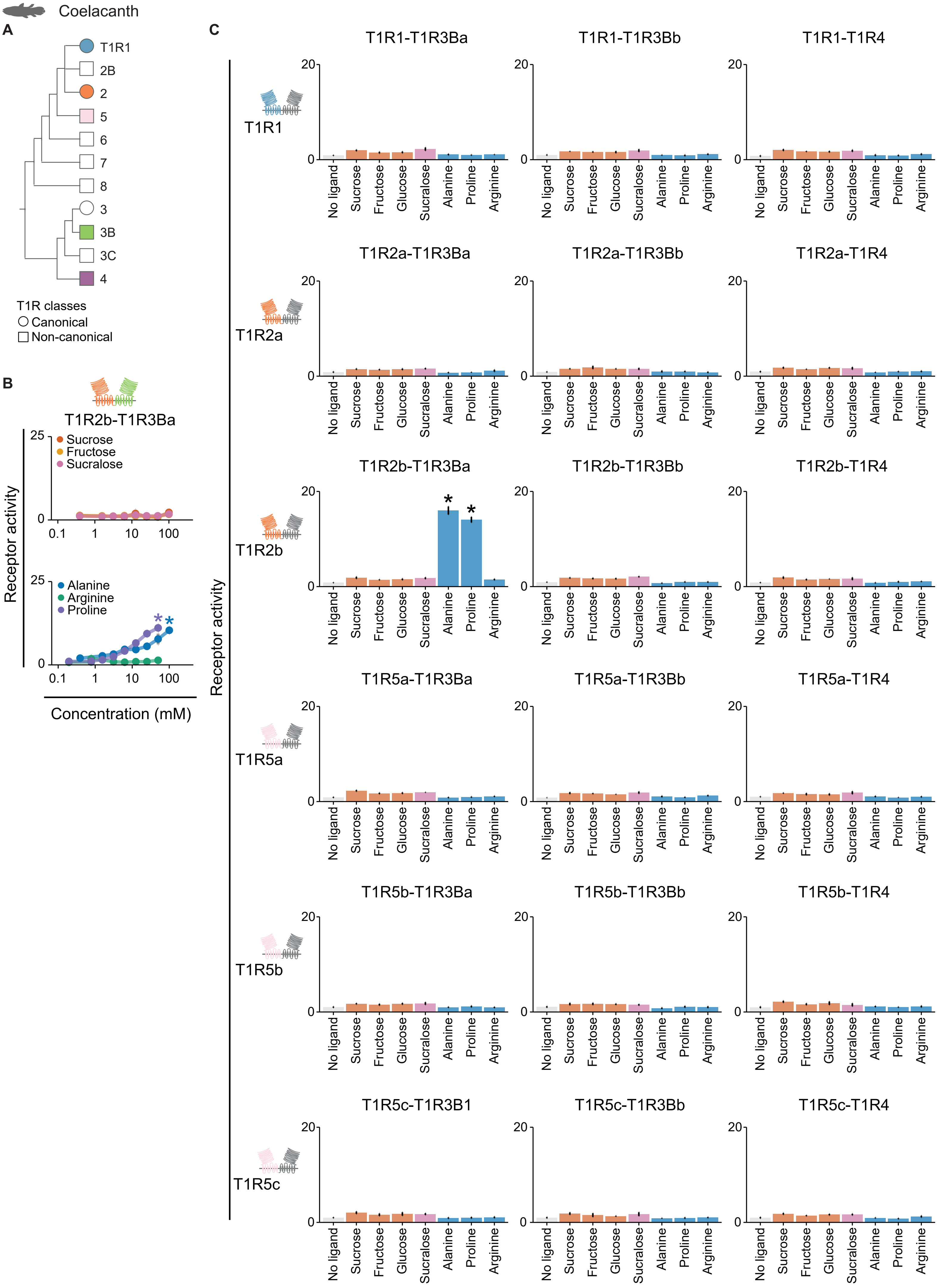

Coelacanth
A
T1R1
2B
2
5
6
7
8
3
3B
3C
4
T1R classes
Canonical
Non-canonical
B
T1R2b-T1R3Ba
Receptor activity
Sucrose
Fructose
Sucralose
Alanine
Arginine
Proline
Concentration (mM)
C
T1R1-T1R3Ba
T1R1-T1R3Bb
T1R1-T1R4
T1R1
T1R2a-T1R3Ba
T1R2a-T1R3Bb
T1R2a-T1R4
T1R2a
T1R2b-T1R3Ba
T1R2b-T1R3Bb
T1R2b-T1R4
T1R2b
T1R5a-T1R3Ba
T1R5a-T1R3Bb
T1R5a-T1R4
T1R5a
T1R5b-T1R3Ba
T1R5b-T1R3Bb
T1R5b-T1R4
T1R5b
T1R5c-T1R3B1
T1R5c-T1R3Bb
T1R5c-T1R4
T1R5c
Receptor activity
No ligand
Sucrose
Fructose
Glucose
Sucralose
Alanine
Proline
Arginine

**fig. S7. Receptor responses of 18 coelacanth (*Latimeria chalumnae*) receptor pairs.** (**A**) Coelacanth possesses nine T1R receptors, including canonical T1R1 and T1R2. (**B**) Coelacanth T1R2b-T1R3Ba shows dose-dependent responses to two amino acids, but not to sugars. (**C**) 17 out of 18 heterodimers of coelacanth did not respond to any of the six ligands tested, with the exception of an amino-acid responsive T1R2-T1R3B pair. Receptor activity is measured as AUC ($x10^3$ RLU; mean ± s.e.m; (B) n = 6; significance (* $p < 0.05$): one-tailed Welch's *t*-tests and BH corrections as in fig. S3; same ligands and concentrations as fig. S3; (C) n = 5-6; significance (* $p < 0.05$): one-tailed Welch's *t*-tests and BH corrections as in fig. S4; same ligand concentrations as fig. S4).

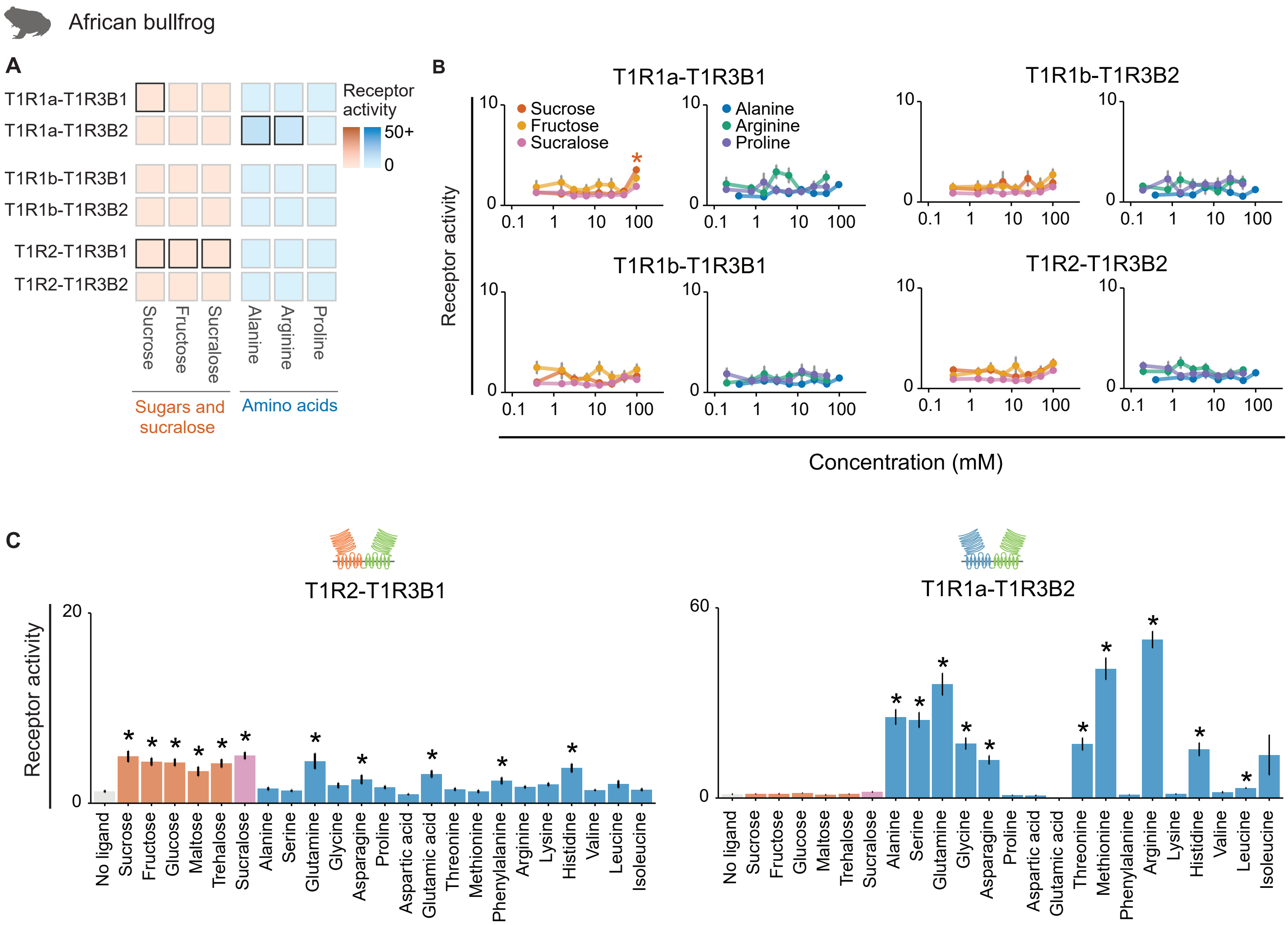

African bullfrog
A
T1R1a-T1R3B1
T1R1a-T1R3B2
T1R1b-T1R3B1
T1R1b-T1R3B2
T1R2-T1R3B1
T1R2-T1R3B2
Receptor activity
50+
0
Sucrose
Fructose
Sucralose
Alanine
Arginine
Proline
Sugars and sucralose
Amino acids
B
T1R1a-T1R3B1
T1R1b-T1R3B2
T1R1b-T1R3B1
T1R2-T1R3B2
Sucrose
Fructose
Sucralose
Alanine
Arginine
Proline
Receptor activity
Concentration (mM)
C
T1R2-T1R3B1
T1R1a-T1R3B2
Receptor activity
No ligand
Sucrose
Fructose
Glucose
Maltose
Trehalose
Sucralose
Alanine
Serine
Glutamine
Glycine
Asparagine
Proline
Aspartic acid
Glutamic acid
Threonine
Methionine
Phenylalanine
Arginine
Lysine
Histidine
Valine
Leucine
Isoleucine

**fig. S8. Functional responses of six African bullfrog (*Pyxicephalus adspersus*) receptor pairs.** (**A, B**) Dose-dependent response summary and data of African bullfrog receptor pairs as well as (**C**) full panel ligand responses of T1R2-T1R3B1 and T1R1a-T1R3B2 heterodimers (same pairs shown in Fig. 2D). Among six receptor pairs tested in African bullfrogs, two pairs (T1R1a-T1R3B1 and T1R2-T1R3B1) are weakly responsive to sugars, whereas T1R1a-T1R3B2 is specialized in amino acid detection. Three pairs (T1R1b-T1R3B1, T1R2-T1R3B2, and T1R1b-T1R3B2) did not respond to any of the six ligands tested. Receptor activity is measured as AUC (x$10^3$ RLU; mean ± s.e.m; (A, B): n = 6-9; significance (* $p < 0.05$): one-tailed Welch's *t*-tests and BH corrections as in fig. S3; same ligands and concentrations as fig. S3; same heatmap colors in (A) as in fig. S3; (C): n = 5-12; significance (* $p < 0.05$): one-tailed Welch's *t*-tests and BH corrections as in fig. S4; same ligands and concentrations as fig. S4).

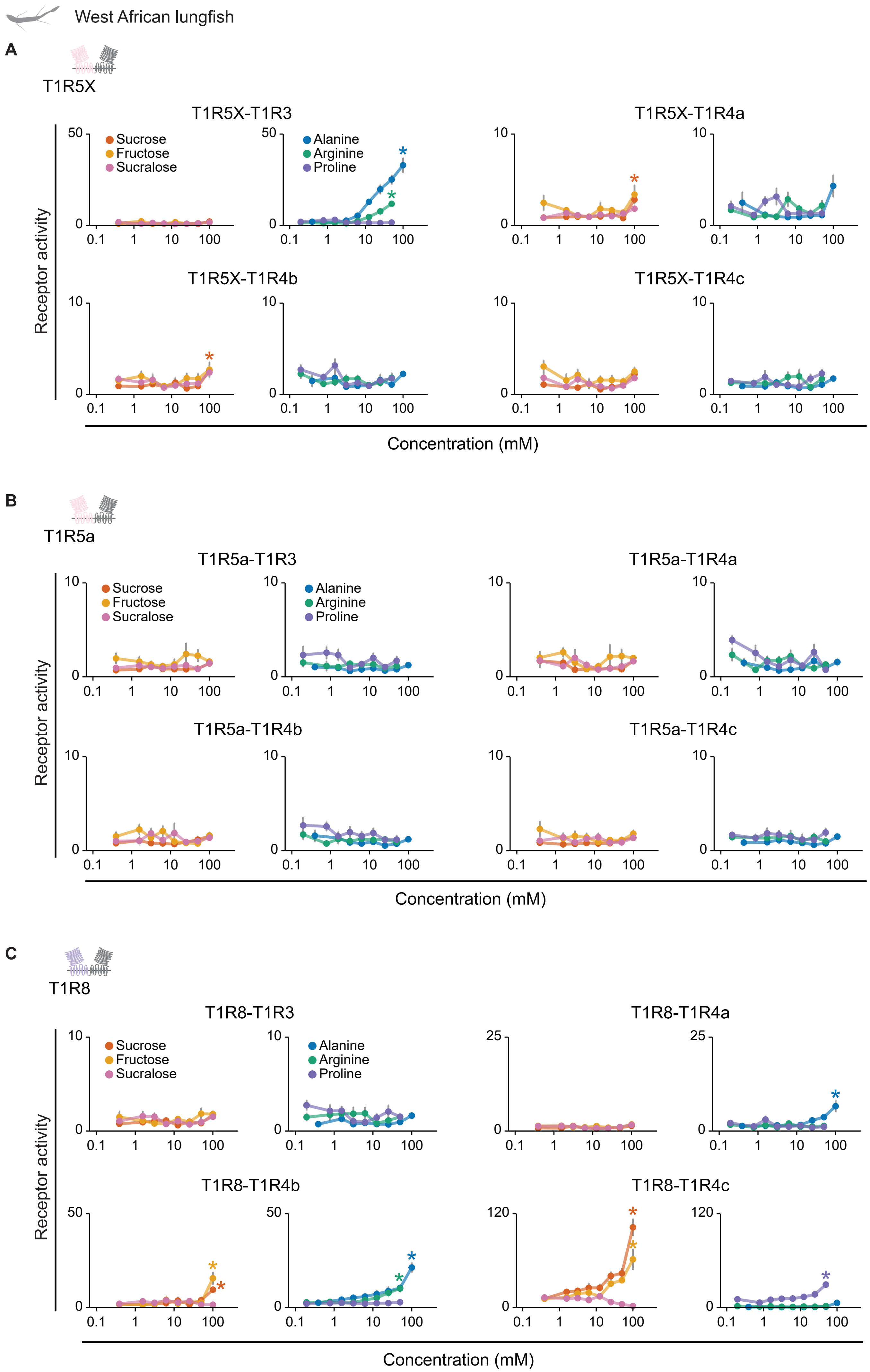
West African lungfish
A
T1R5X
T1R5X-T1R3
T1R5X-T1R4a
Sucrose
Fructose
Sucralose
Alanine
Arginine
Proline
T1R5X-T1R4b
T1R5X-T1R4c
Receptor activity
Concentration (mM)
B
T1R5a
T1R5a-T1R3
T1R5a-T1R4a
Sucrose
Fructose
Sucralose
Alanine
Arginine
Proline
T1R5a-T1R4b
T1R5a-T1R4c
Receptor activity
Concentration (mM)
C
T1R8
T1R8-T1R3
T1R8-T1R4a
Sucrose
Fructose
Sucralose
Alanine
Arginine
Proline
T1R8-T1R4b
T1R8-T1R4c
Receptor activity
Concentration (mM)

**fig. S9. Dose-dependent responses of 12 receptor pairs in West African lungfish (*Protopterus annectens*).** (**A-C**) Functional tests of 12 heterodimers formed by seven receptors of West African lungfish show that seven pairs are responsive to sugars or amino acids. Two T1R5X-based pairs (A) and two T1R8-based pairs (C) show sucrose responses, whereas T1R5a (B) did not form any receptor pair responsive to the six ligands tested. Among these, T1R8-T1R4c displays the highest relative response to sugars compared to other vertebrate receptor pairs tested so far. Receptor activity is measured as AUC (x$10^3$ RLU; mean ± s.e.m; n = 5-8; significance (* $p < 0.05$): one-tailed Welch's *t*-tests and BH corrections as in fig. S3; same ligands and concentrations as fig. S3).

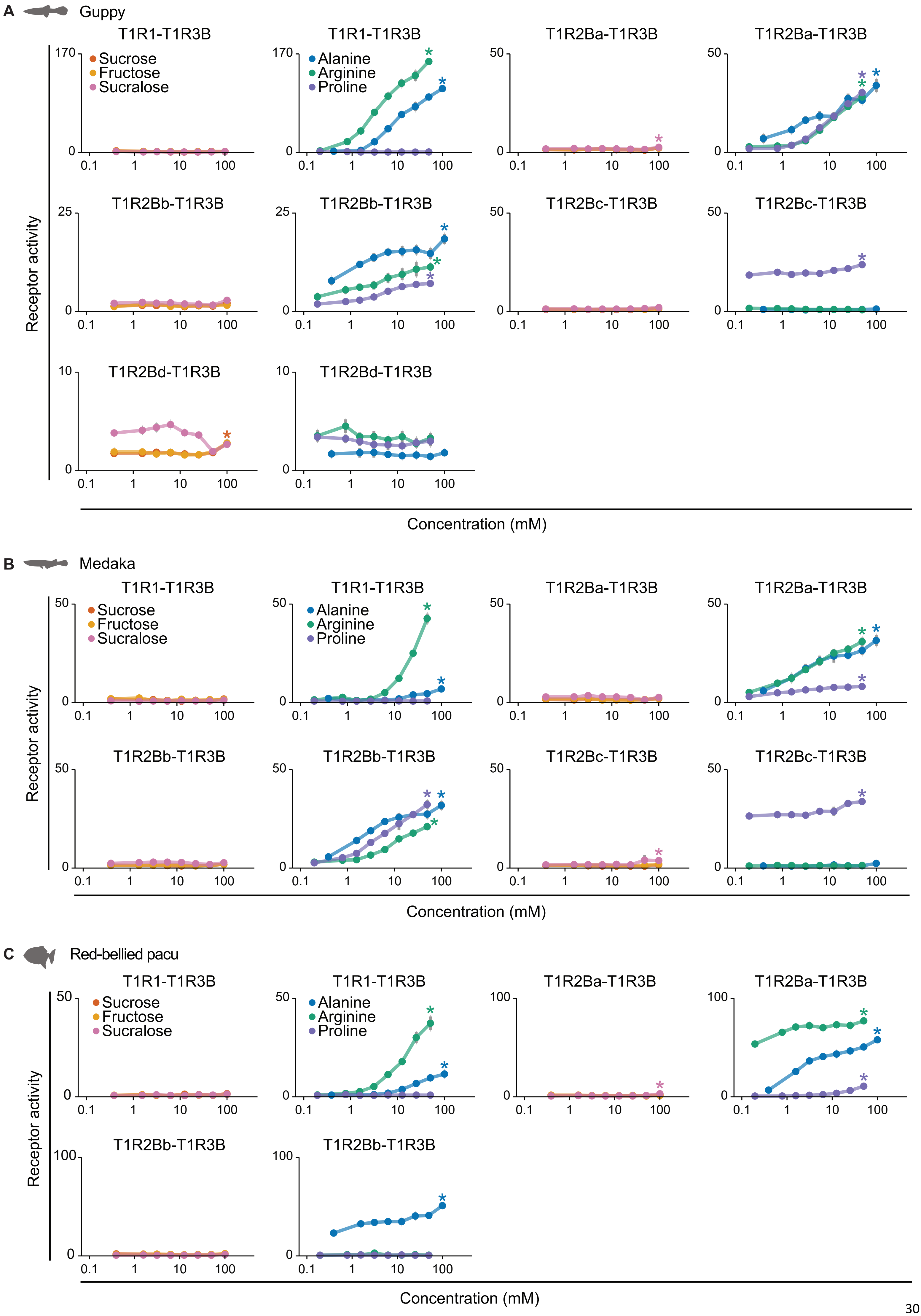
A
Guppy
T1R1-T1R3B
T1R2Ba-T1R3B
T1R2Bb-T1R3B
T1R2Bc-T1R3B
T1R2Bd-T1R3B
Sucrose
Fructose
Sucralose
Alanine
Arginine
Proline
Receptor activity
Concentration (mM)
B
Medaka
T1R1-T1R3B
T1R2Ba-T1R3B
T1R2Bb-T1R3B
T1R2Bc-T1R3B
Sucrose
Fructose
Sucralose
Alanine
Arginine
Proline
Receptor activity
Concentration (mM)
C
Red-bellied pacu
T1R1-T1R3B
T1R2Ba-T1R3B
T1R2Bb-T1R3B
Sucrose
Fructose
Sucralose
Alanine
Arginine
Proline
Receptor activity
Concentration (mM)

**fig. S10. Dose-dependent responses of receptor pairs of three teleosts.** (**A**) Fruit-eating guppy (*Poecilia reticulata*) possesses four T1R pairs responsive to amino acids, with one (T1R2Bc-T1R3B) specialized in proline detection; T1R2Bd-T1R3B responds to sucrose. (**B**) Four receptor pairs of medaka (*Oryzias latipes*) respond primarily to amino acids, of which T1R2Bc-T1R3B shows high sensitivity to proline and also a small response to sucralose. (**C**) Three receptor pairs of red-bellied pacu (*Piaractus brachypomus*) show robust responses to amino acids and T1R2Ba-T1R3B also responds to sucralose. Receptor activity is calculated as AUC (x$10^3$ RLU; mean ± s.e.m; n = 6-9; significance (* $p < 0.05$): one-tailed Welch's *t*-tests and BH corrections as in fig. S3; same ligands and concentrations as fig. S3).

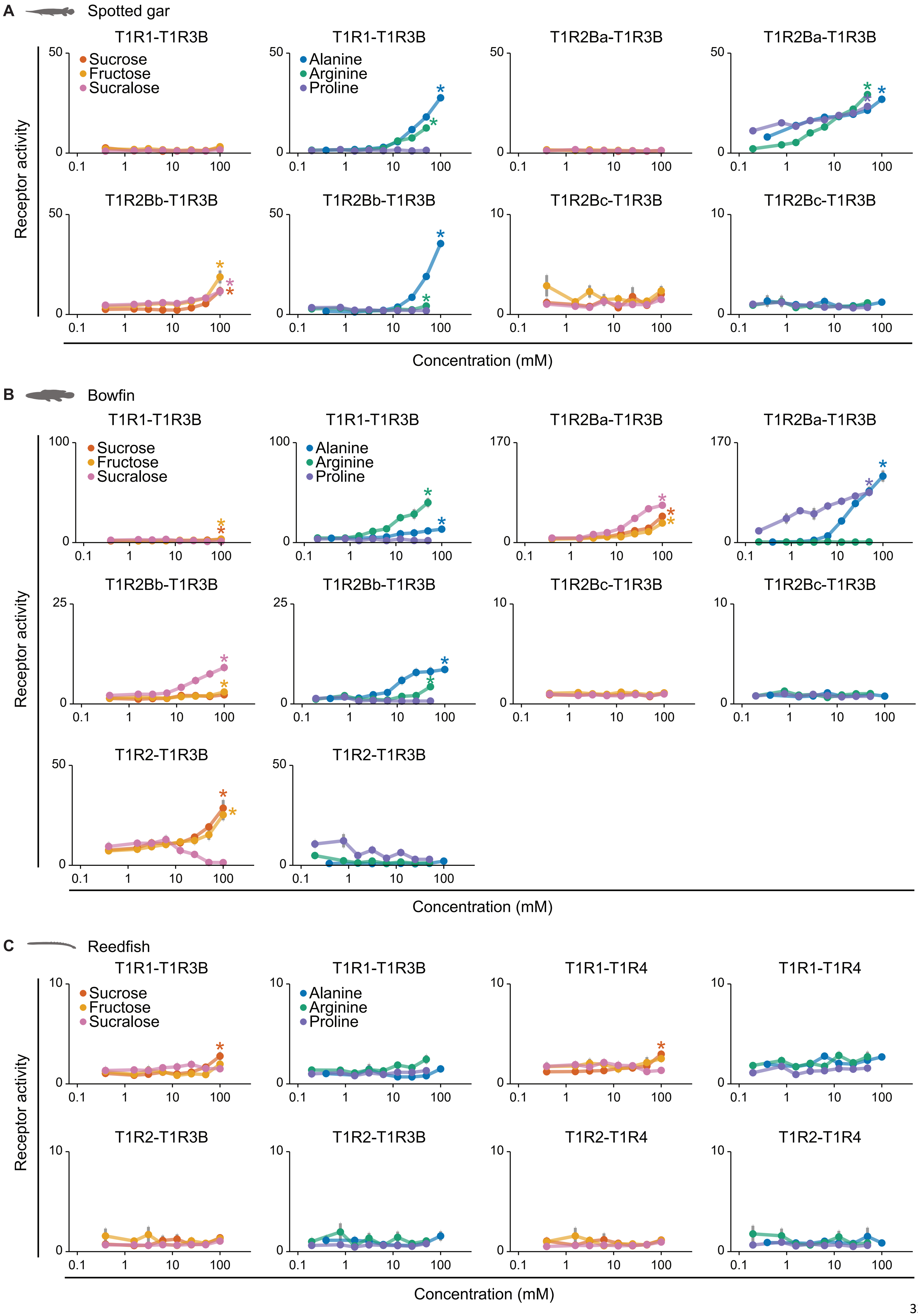
A
Spotted gar
B
Bowfin
C
Reedfish
T1R1-T1R3B
T1R2Ba-T1R3B
T1R2Bb-T1R3B
T1R2Bc-T1R3B
T1R2-T1R3B
T1R1-T1R4
T1R2-T1R4
Sucrose
Fructose
Sucralose
Alanine
Arginine
Proline
Receptor activity
Concentration (mM)

**fig. S11. Dose-dependent responses of receptor pairs of three non-teleosts.** (**A**) Spotted gar (*Lepisosteus oculatus*) possesses two receptor pairs specialized in amino acids, whereas T1R2Bb-T1R3B is responsive to both amino acids and sugars. (**B**) Bowfin (*Amia calva*) has four receptor pairs responsive to sugars: T1R1-T1R3B, T1R2Ba-T1R3B, T1R2Bb-T1R3B and T1R2-T1R3B (the latter is most sensitive to sugars than to amino acids). (**C**) Whereas reedfish (*Erpetoichthys calabaricus*) T1R1-T1R3B and T1R1-T1R4 show small responses to sucrose, its T1R2-based receptors appear nonfunctional. Receptor activity is measured as AUC (x$10^3$ RLU; mean ± s.e.m; n = 6-9; significance (* $p < 0.05$): one-tailed Welch's *t*-tests and BH corrections like in fig. S3; same ligands and concentrations as fig. S3).

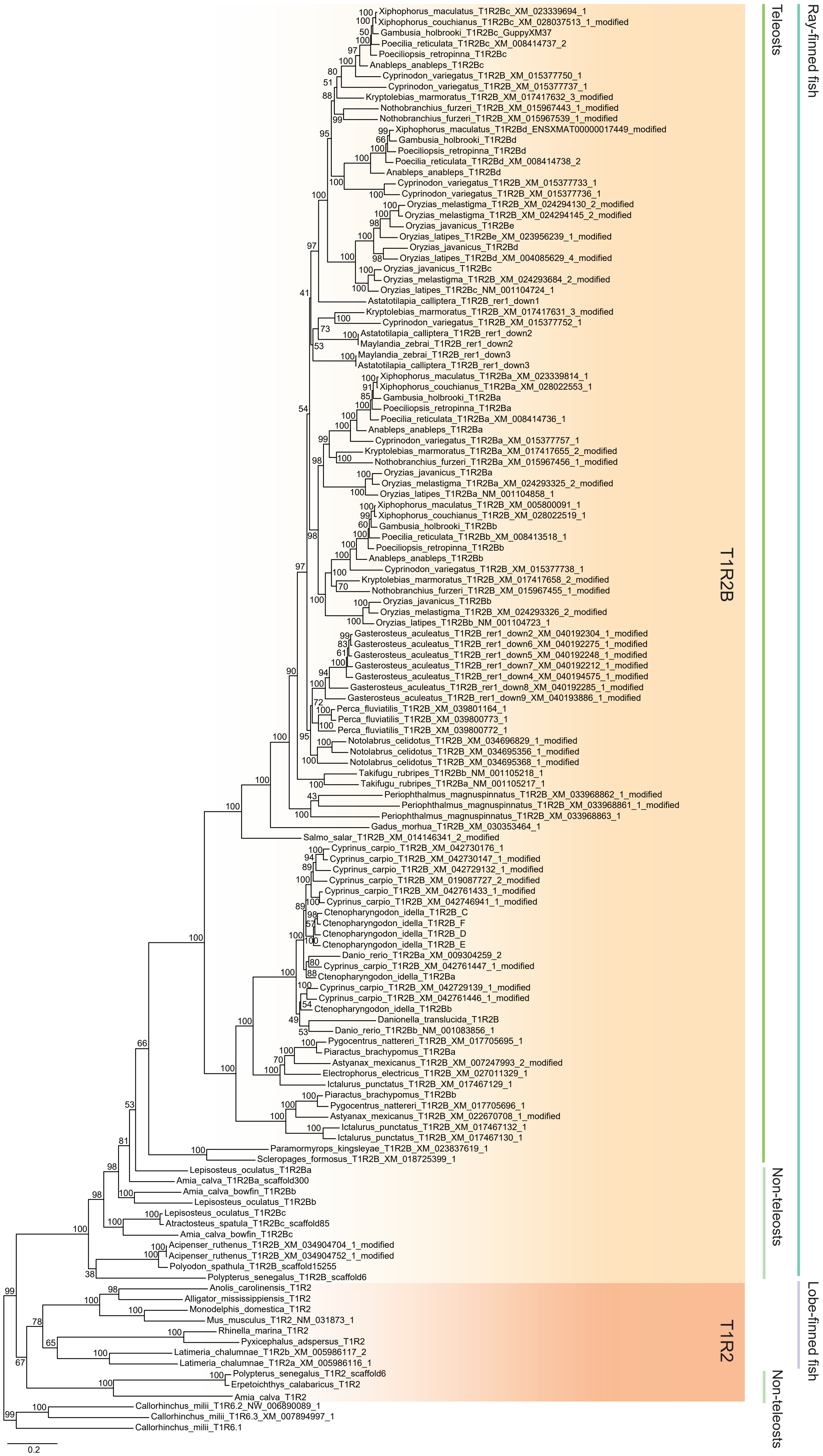
Ray-finned fish
Teleosts
Non-teleosts
Lobe-finned fish
Non-teleosts
T1R2B
T1R2
Xiphophorus_maculatus_T1R2Bc_XM_023339694_1
Xiphophorus_couchianus_T1R2Bc_XM_028037513_1_modified
Gambusia_holbrooki_T1R2Bc_GuppyXM37
Poecilia_reticulata_T1R2Bc_XM_008414737_2
Poeciliopsis_retropinna_T1R2Bc
Anableps_anableps_T1R2Bc
Cyprinodon_variegatus_T1R2B_XM_015377750_1
Cyprinodon_variegatus_T1R2B_XM_015377737_1
Kryptolebias_marmoratus_T1R2B_XM_017417632_3_modified
Nothobranchius_furzeri_T1R2B_XM_015967443_1_modified
Nothobranchius_furzeri_T1R2B_XM_015967539_1_modified
Xiphophorus_maculatus_T1R2Bd_ENSXMAT00000017449_modified
Gambusia_holbrooki_T1R2Bd
Poeciliopsis_retropinna_T1R2Bd
Poecilia_reticulata_T1R2Bd_XM_008414738_2
Anableps_anableps_T1R2Bd
Cyprinodon_variegatus_T1R2B_XM_015377733_1
Cyprinodon_variegatus_T1R2B_XM_015377736_1
Oryzias_melastigma_T1R2B_XM_024294130_2_modified
Oryzias_melastigma_T1R2B_XM_024294145_2_modified
Oryzias_javanicus_T1R2Be
Oryzias_latipes_T1R2Be_XM_023956239_1_modified
Oryzias_javanicus_T1R2Bd
Oryzias_latipes_T1R2Bd_XM_004085629_4_modified
Oryzias_javanicus_T1R2Bc
Oryzias_melastigma_T1R2B_XM_024293684_2_modified
Oryzias_latipes_T1R2Bc_NM_001104724_1
Astatotilapia_calliptera_T1R2B_rer1_down1
Kryptolebias_marmoratus_T1R2B_XM_017417631_3_modified
Cyprinodon_variegatus_T1R2B_XM_015377752_1
Astatotilapia_calliptera_T1R2B_rer1_down2
Maylandia_zebrai_T1R2B_rer1_down2
Maylandia_zebrai_T1R2B_rer1_down3
Astatotilapia_calliptera_T1R2B_rer1_down3
Xiphophorus_maculatus_T1R2Ba_XM_023339814_1
Xiphophorus_couchianus_T1R2Ba_XM_028022553_1
Gambusia_holbrooki_T1R2Ba
Poeciliopsis_retropinna_T1R2Ba
Poecilia_reticulata_T1R2Ba_XM_008414736_1
Anableps_anableps_T1R2Ba
Cyprinodon_variegatus_T1R2Ba_XM_015377757_1
Kryptolebias_marmoratus_T1R2Ba_XM_017417655_2_modified
Nothobranchius_furzeri_T1R2Ba_XM_015967456_1_modified
Oryzias_javanicus_T1R2Ba
Oryzias_melastigma_T1R2Ba_XM_024293325_2_modified
Oryzias_latipes_T1R2Ba_NM_001104858_1
Xiphophorus_maculatus_T1R2B_XM_005800091_1
Xiphophorus_couchianus_T1R2B_XM_028022519_1
Gambusia_holbrooki_T1R2Bb
Poecilia_reticulata_T1R2Bb_XM_008413518_1
Poeciliopsis_retropinna_T1R2Bb
Anableps_anableps_T1R2Bb
Cyprinodon_variegatus_T1R2B_XM_015377738_1
Kryptolebias_marmoratus_T1R2B_XM_017417658_2_modified
Nothobranchius_furzeri_T1R2B_XM_015967455_1_modified
Oryzias_javanicus_T1R2Bb
Oryzias_melastigma_T1R2B_XM_024293326_2_modified
Oryzias_latipes_T1R2Bb_NM_001104723_1
Gasterosteus_aculeatus_T1R2B_rer1_down2_XM_040192304_1_modified
Gasterosteus_aculeatus_T1R2B_rer1_down6_XM_040192275_1_modified
Gasterosteus_aculeatus_T1R2B_rer1_down5_XM_040192248_1_modified
Gasterosteus_aculeatus_T1R2B_rer1_down7_XM_040192212_1_modified
Gasterosteus_aculeatus_T1R2B_rer1_down4_XM_040194575_1_modified
Gasterosteus_aculeatus_T1R2B_rer1_down8_XM_040192285_1_modified
Gasterosteus_aculeatus_T1R2B_rer1_down9_XM_040193886_1_modified
Perca_fluviatilis_T1R2B_XM_039801164_1
Perca_fluviatilis_T1R2B_XM_039800773_1
Perca_fluviatilis_T1R2B_XM_039800772_1
Notolabrus_celidotus_T1R2B_XM_034696829_1_modified
Notolabrus_celidotus_T1R2B_XM_034695356_1_modified
Notolabrus_celidotus_T1R2B_XM_034695368_1_modified
Takifugu_rubripes_T1R2Bb_NM_001105218_1
Takifugu_rubripes_T1R2Ba_NM_001105217_1
Periophthalmus_magnuspinnatus_T1R2B_XM_033968862_1_modified
Periophthalmus_magnuspinnatus_T1R2B_XM_033968861_1_modified
Periophthalmus_magnuspinnatus_T1R2B_XM_033968863_1
Gadus_morhua_T1R2B_XM_030353464_1
Salmo_salar_T1R2B_XM_014146341_2_modified
Cyprinus_carpio_T1R2B_XM_042730176_1
Cyprinus_carpio_T1R2B_XM_042730147_1_modified
Cyprinus_carpio_T1R2B_XM_042729132_1_modified
Cyprinus_carpio_T1R2B_XM_019087727_2_modified
Cyprinus_carpio_T1R2B_XM_042761433_1_modified
Cyprinus_carpio_T1R2B_XM_042746941_1_modified
Ctenopharyngodon_idella_T1R2B_C
Ctenopharyngodon_idella_T1R2B_F
Ctenopharyngodon_idella_T1R2B_D
Ctenopharyngodon_idella_T1R2B_E
Danio_rerio_T1R2Ba_XM_009304259_2
Cyprinus_carpio_T1R2B_XM_042761447_1_modified
Ctenopharyngodon_idella_T1R2Ba
Cyprinus_carpio_T1R2B_XM_042729139_1_modified
Cyprinus_carpio_T1R2B_XM_042761446_1_modified
Ctenopharyngodon_idella_T1R2Bb
Danionella_translucida_T1R2B
Danio_rerio_T1R2Bb_NM_001083856_1
Pygocentrus_nattereri_T1R2B_XM_017705695_1
Piaractus_brachypomus_T1R2Ba
Astyanax_mexicanus_T1R2B_XM_007247993_2_modified
Electrophorus_electricus_T1R2B_XM_027011329_1
Ictalurus_punctatus_T1R2B_XM_017467129_1
Piaractus_brachypomus_T1R2Bb
Pygocentrus_nattereri_T1R2B_XM_017705696_1
Astyanax_mexicanus_T1R2B_XM_022670708_1_modified
Ictalurus_punctatus_T1R2B_XM_017467132_1
Ictalurus_punctatus_T1R2B_XM_017467130_1
Paramormyrops_kingsleyae_T1R2B_XM_023837619_1
Scleropages_formosus_T1R2B_XM_018725399_1
Lepisosteus_oculatus_T1R2Ba
Amia_calva_T1R2Ba_scaffold300
Amia_calva_bowfin_T1R2Bb
Lepisosteus_oculatus_T1R2Bb
Lepisosteus_oculatus_T1R2Bc
Atractosteus_spatula_T1R2Bc_scaffold85
Amia_calva_bowfin_T1R2Bc
Acipenser_ruthenus_T1R2B_XM_034904704_1_modified
Acipenser_ruthenus_T1R2B_XM_034904752_1_modified
Polyodon_spathula_T1R2B_scaffold15255
Polypterus_senegalus_T1R2B_scaffold6
Anolis_carolinensis_T1R2
Alligator_mississippiensis_T1R2
Monodelphis_domestica_T1R2
Mus_musculus_T1R2_NM_031873_1
Rhinella_marina_T1R2
Pyxicephalus_adspersus_T1R2
Latimeria_chalumnae_T1R2b_XM_005986117_2
Latimeria_chalumnae_T1R2a_XM_005986116_1
Polypterus_senegalus_T1R2_scaffold6
Erpetoichthys_calabaricus_T1R2
Amia_calva_T1R2
Callorhinchus_milii_T1R6.2_NW_006890089_1
Callorhinchus_milii_T1R6.3_XM_007894997_1
Callorhinchus_milii_T1R6.1
0.2

**fig. S12. Maximum-likelihood tree featuring ray-finned fish *T1R2B*, bony vertebrate *T1R2* and shark *T1R6*.** The maximum-likelihood tree was inferred with 133 *T1R* nucleotide sequences of 64 jawed vertebrate species (light orange shading, *T1R2B*; dark orange shading, *T1R2*; no shading, *T1R6*) and the GTR+G+I model using RAxML. Since T1R2B duplications are often lineage-specific and often not shared across species, the naming convention here (ie. T1R2Ba, T1R2Bb) refers to copies within a species, not to orthologs across different lineages. Node support values represent bootstrap values with 100 replicates.

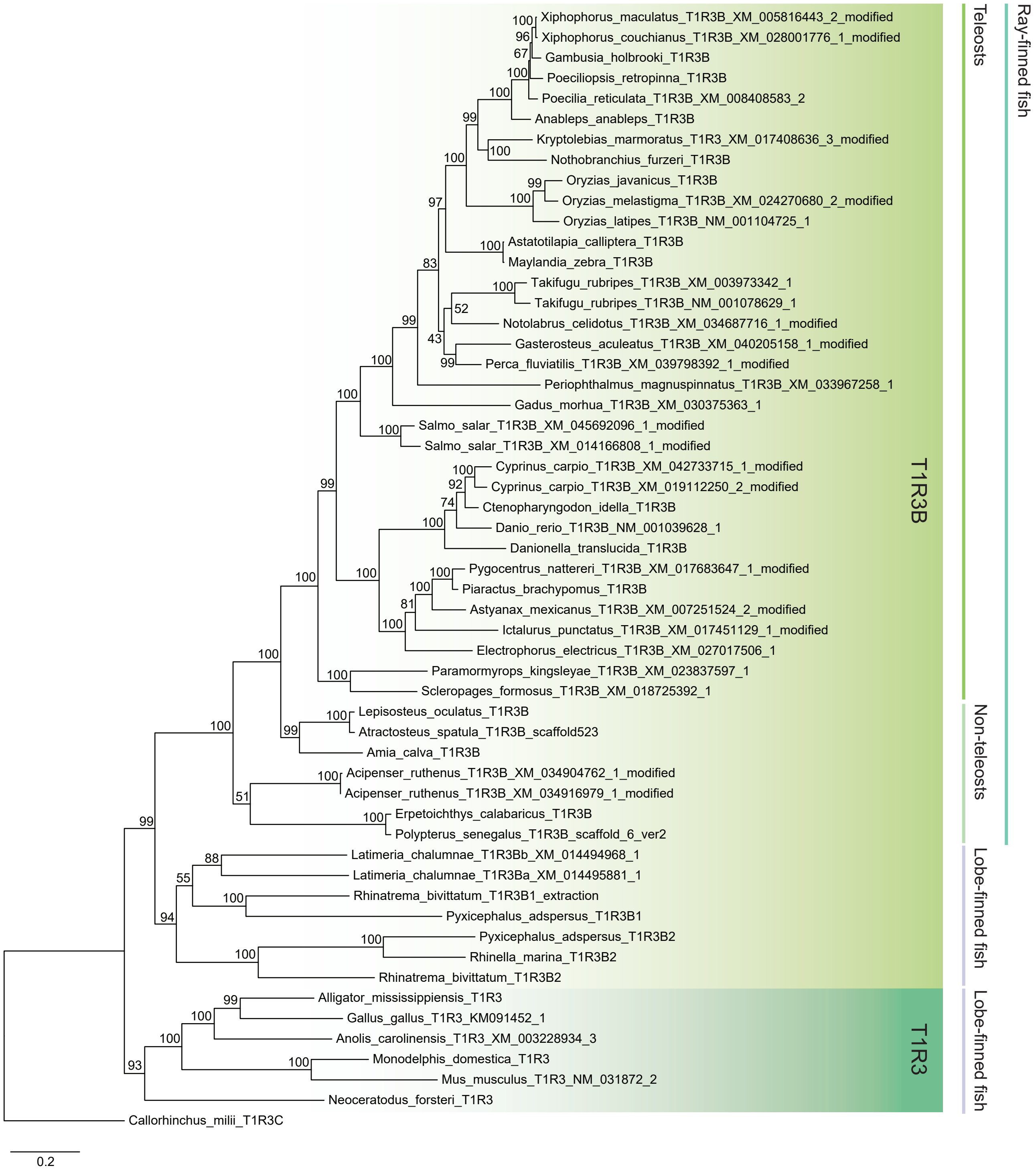

Xiphophorus_maculatus_T1R3B_XM_005816443_2_modified
Xiphophorus_couchianus_T1R3B_XM_028001776_1_modified
Gambusia_holbrooki_T1R3B
Poeciliopsis_retropinna_T1R3B
Poecilia_reticulata_T1R3B_XM_008408583_2
Anableps_anableps_T1R3B
Kryptolebias_marmoratus_T1R3_XM_017408636_3_modified
Nothobranchius_furzeri_T1R3B
Oryzias_javanicus_T1R3B
Oryzias_melastigma_T1R3B_XM_024270680_2_modified
Oryzias_latipes_T1R3B_NM_001104725_1
Astatotilapia_calliptera_T1R3B
Maylandia_zebra_T1R3B
Takifugu_rubripes_T1R3B_XM_003973342_1
Takifugu_rubripes_T1R3B_NM_001078629_1
Notolabrus_celidotus_T1R3B_XM_034687716_1_modified
Gasterosteus_aculeatus_T1R3B_XM_040205158_1_modified
Perca_fluviatilis_T1R3B_XM_039798392_1_modified
Periophthalmus_magnuspinnatus_T1R3B_XM_033967258_1
Gadus_morhua_T1R3B_XM_030375363_1
Salmo_salar_T1R3B_XM_045692096_1_modified
Salmo_salar_T1R3B_XM_014166808_1_modified
Cyprinus_carpio_T1R3B_XM_042733715_1_modified
Cyprinus_carpio_T1R3B_XM_019112250_2_modified
Ctenopharyngodon_idella_T1R3B
Danio_rerio_T1R3B_NM_001039628_1
Danionella_translucida_T1R3B
Pygocentrus_nattereri_T1R3B_XM_017683647_1_modified
Piaractus_brachypomus_T1R3B
Astyanax_mexicanus_T1R3B_XM_007251524_2_modified
Ictalurus_punctatus_T1R3B_XM_017451129_1_modified
Electrophorus_electricus_T1R3B_XM_027017506_1
Paramormyrops_kingsleyae_T1R3B_XM_023837597_1
Scleropages_formosus_T1R3B_XM_018725392_1
Lepisosteus_oculatus_T1R3B
Atractosteus_spatula_T1R3B_scaffold523
Amia_calva_T1R3B
Acipenser_ruthenus_T1R3B_XM_034904762_1_modified
Acipenser_ruthenus_T1R3B_XM_034916979_1_modified
Erpetoichthys_calabaricus_T1R3B
Polypterus_senegalus_T1R3B_scaffold_6_ver2
Latimeria_chalumnae_T1R3Bb_XM_014494968_1
Latimeria_chalumnae_T1R3Ba_XM_014495881_1
Rhinatrema_bivittatum_T1R3B1_extraction
Pyxicephalus_adspersus_T1R3B1
Pyxicephalus_adspersus_T1R3B2
Rhinella_marina_T1R3B2
Rhinatrema_bivittatum_T1R3B2
Alligator_mississippiensis_T1R3
Gallus_gallus_T1R3_KM091452_1
Anolis_carolinensis_T1R3_XM_003228934_3
Monodelphis_domestica_T1R3
Mus_musculus_T1R3_NM_031872_2
Neoceratodus_forsteri_T1R3
Callorhinchus_milii_T1R3C
T1R3B
T1R3
Teleosts
Non-teleosts
Lobe-finned fish
Lobe-finned fish
Ray-finned fish
0.2

**fig. S13. Maximum-likelihood tree of *T1R3, T1R3B* and *T1R3C* across jawed vertebrates.** The maximum-likelihood tree was inferred with 55 nucleotide sequences of lobe-finned fish *T1R3* (dark green shading), bony vertebrate *T1R3B* (light green shading) and shark *T1R3C* (no shading) from 49 species and the GTR+G+I model using RAxML. Node support values indicate bootstrap values with 100 replicates.

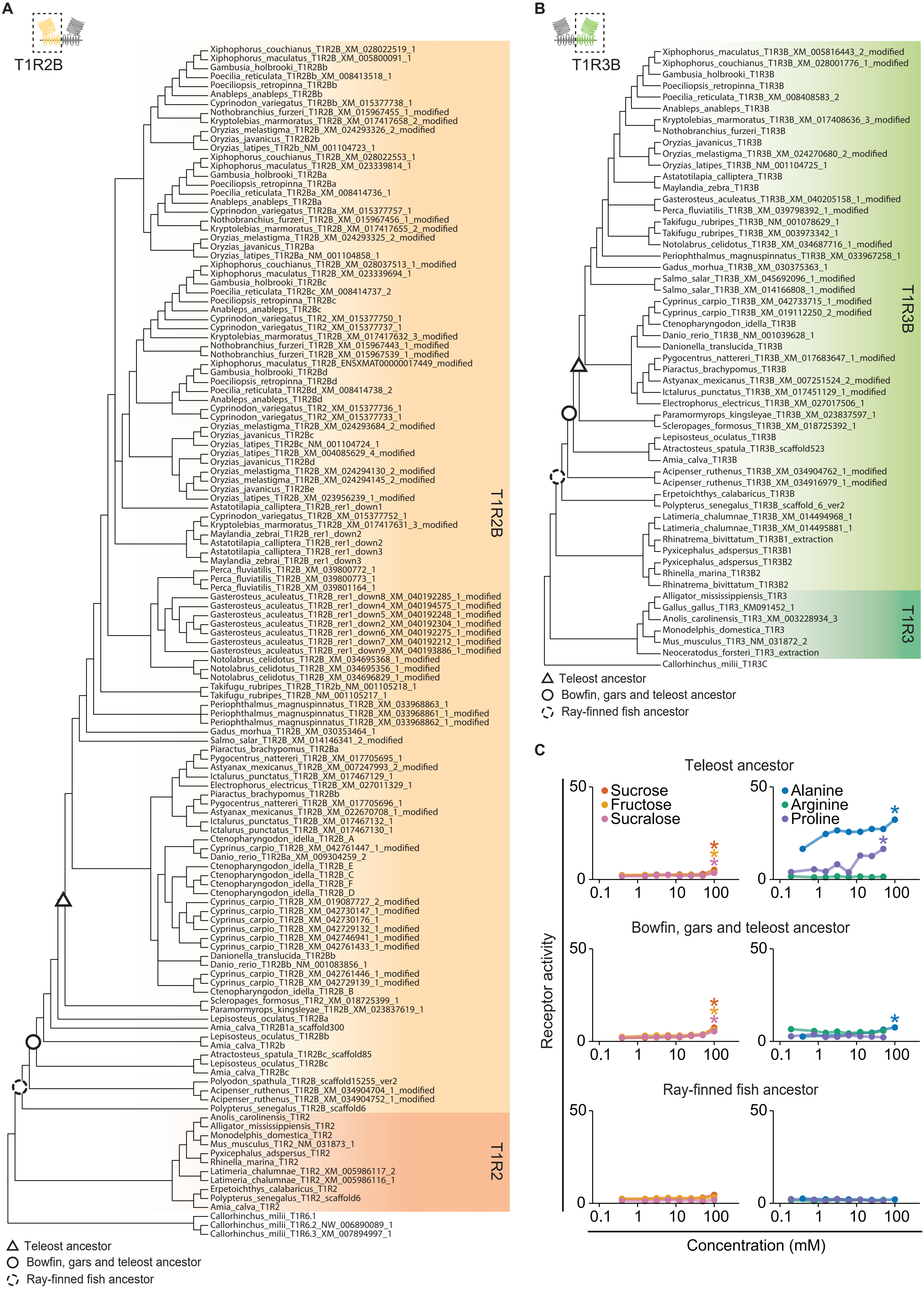

A
T1R2B
Xiphophorus_couchianus_T1R2B_XM_028022519_1
Xiphophorus_maculatus_T1R2B_XM_005800091_1
Gambusia_holbrooki_T1R2Bb
Poecilia_reticulata_T1R2Bb_XM_008413518_1
Poeciliopsis_retropinna_T1R2Bb
Anableps_anableps_T1R2Bb
Cyprinodon_variegatus_T1R2Bb_XM_015377738_1
Nothobranchius_furzeri_T1R2B_XM_015967455_1_modified
Kryptolebias_marmoratus_T1R2B_XM_017417658_2_modified
Oryzias_melastigma_T1R2B_XM_024293326_2_modified
Oryzias_javanicus_T1R2B2b
Oryzias_latipes_T1R2b_NM_001104723_1
Xiphophorus_couchianus_T1R2B_XM_028022553_1
Xiphophorus_maculatus_T1R2B_XM_023339814_1
Gambusia_holbrooki_T1R2Ba
Poeciliopsis_retropinna_T1R2Ba
Poecilia_reticulata_T1R2Ba_XM_008414736_1
Anableps_anableps_T1R2Ba
Cyprinodon_variegatus_T1R2Ba_XM_015377757_1
Nothobranchius_furzeri_T1R2B_XM_015967456_1_modified
Kryptolebias_marmoratus_T1R2B_XM_017417655_2_modified
Oryzias_melastigma_T1R2B_XM_024293325_2_modified
Oryzias_javanicus_T1R2Ba
Oryzias_latipes_T1R2Ba_NM_001104858_1
Xiphophorus_couchianus_T1R2B_XM_028037513_1_modified
Xiphophorus_maculatus_T1R2B_XM_023339694_1
Gambusia_holbrooki_T1R2Bc
Poecilia_reticulata_T1R2Bc_XM_008414737_2
Poeciliopsis_retropinna_T1R2Bc
Anableps_anableps_T1R2Bc
Cyprinodon_variegatus_T1R2_XM_015377750_1
Cyprinodon_variegatus_T1R2_XM_015377737_1
Kryptolebias_marmoratus_T1R2B_XM_017417632_3_modified
Nothobranchius_furzeri_T1R2B_XM_015967443_1_modified
Nothobranchius_furzeri_T1R2B_XM_015967539_1_modified
Xiphophorus_maculatus_T1R2B_ENSXMAT00000017449_modified
Gambusia_holbrooki_T1R2Bd
Poeciliopsis_retropinna_T1R2Bd
Poecilia_reticulata_T1R2Bd_XM_008414738_2
Anableps_anableps_T1R2Bd
Cyprinodon_variegatus_T1R2_XM_015377736_1
Cyprinodon_variegatus_T1R2_XM_015377733_1
Oryzias_melastigma_T1R2B_XM_024293684_2_modified
Oryzias_javanicus_T1R2Bc
Oryzias_latipes_T1R2Bc_NM_001104724_1
Oryzias_latipes_T1R2B_XM_004085629_4_modified
Oryzias_javanicus_T1R2Bd
Oryzias_melastigma_T1R2B_XM_024294130_2_modified
Oryzias_melastigma_T1R2B_XM_024294145_2_modified
Oryzias_javanicus_T1R2Be
Oryzias_latipes_T1R2B_XM_023956239_1_modified
Astatotilapia_calliptera_T1R2B_rer1_down1
Cyprinodon_variegatus_T1R2B_XM_015377752_1
Kryptolebias_marmoratus_T1R2B_XM_017417631_3_modified
Maylandia_zebrai_T1R2B_rer1_down2
Astatotilapia_calliptera_T1R2B_rer1_down2
Astatotilapia_calliptera_T1R2B_rer1_down3
Maylandia_zebrai_T1R2B_rer1_down3
Perca_fluviatilis_T1R2B_XM_039800772_1
Perca_fluviatilis_T1R2B_XM_039800773_1
Perca_fluviatilis_T1R2B_XM_039801164_1
Gasterosteus_aculeatus_T1R2B_rer1_down8_XM_040192285_1_modified
Gasterosteus_aculeatus_T1R2B_rer1_down4_XM_040194575_1_modified
Gasterosteus_aculeatus_T1R2B_rer1_down5_XM_040192248_1_modified
Gasterosteus_aculeatus_T1R2B_rer1_down2_XM_040192304_1_modified
Gasterosteus_aculeatus_T1R2B_rer1_down6_XM_040192275_1_modified
Gasterosteus_aculeatus_T1R2B_rer1_down7_XM_040192212_1_modified
Gasterosteus_aculeatus_T1R2B_rer1_down9_XM_040193886_1_modified
Notolabrus_celidotus_T1R2B_XM_034695368_1_modified
Notolabrus_celidotus_T1R2B_XM_034695356_1_modified
Notolabrus_celidotus_T1R2B_XM_034696829_1_modified
Takifugu_rubripes_T1R2B_T1R2b_NM_001105218_1
Takifugu_rubripes_T1R2B_NM_001105217_1
Periophthalmus_magnuspinnatus_T1R2B_XM_033968863_1
Periophthalmus_magnuspinnatus_T1R2B_XM_033968861_1_modified
Periophthalmus_magnuspinnatus_T1R2B_XM_033968862_1_modified
Gadus_morhua_T1R2B_XM_030353464_1
Salmo_salar_T1R2B_XM_014146341_2_modified
Piaractus_brachypomus_T1R2Ba
Pygocentrus_nattereri_T1R2B_XM_017705695_1
Astyanax_mexicanus_T1R2B_XM_007247993_2_modified
Ictalurus_punctatus_T1R2B_XM_017467129_1
Electrophorus_electricus_T1R2B_XM_027011329_1
Piaractus_brachypomus_T1R2Bb
Pygocentrus_nattereri_T1R2B_XM_017705696_1
Astyanax_mexicanus_T1R2B_XM_022670708_1_modified
Ictalurus_punctatus_T1R2B_XM_017467132_1
Ictalurus_punctatus_T1R2B_XM_017467130_1
Ctenopharyngodon_idella_T1R2B_A
Cyprinus_carpio_T1R2B_XM_042761447_1_modified
Danio_rerio_T1R2Ba_XM_009304259_2
Ctenopharyngodon_idella_T1R2B_E
Ctenopharyngodon_idella_T1R2B_C
Ctenopharyngodon_idella_T1R2B_F
Ctenopharyngodon_idella_T1R2B_D
Cyprinus_carpio_T1R2B_XM_019087727_2_modified
Cyprinus_carpio_T1R2B_XM_042730147_1_modified
Cyprinus_carpio_T1R2B_XM_042730176_1
Cyprinus_carpio_T1R2B_XM_042729132_1_modified
Cyprinus_carpio_T1R2B_XM_042746941_1_modified
Cyprinus_carpio_T1R2B_XM_042761433_1_modified
Danionella_translucida_T1R2Bb
Danio_rerio_T1R2Bb_NM_001083856_1
Cyprinus_carpio_T1R2B_XM_042761446_1_modified
Cyprinus_carpio_T1R2B_XM_042729139_1_modified
Ctenopharyngodon_idella_T1R2B_B
Scleropages_formosus_T1R2_XM_018725399_1
Paramormyrops_kingsleyae_T1R2B_XM_023837619_1
Lepisosteus_oculatus_T1R2Ba
Amia_calva_T1R2B1a_scaffold300
Lepisosteus_oculatus_T1R2Bb
Amia_calva_T1R2b
Atractosteus_spatula_T1R2Bc_scaffold85
Lepisosteus_oculatus_T1R2Bc
Amia_calva_T1R2Bc
Polyodon_spathula_T1R2B_scaffold15255_ver2
Acipenser_ruthenus_T1R2B_XM_034904704_1_modified
Acipenser_ruthenus_T1R2B_XM_034904752_1_modified
Polypterus_senegalus_T1R2B_scaffold6
Anolis_carolinensis_T1R2
Alligator_mississippiensis_T1R2
Monodelphis_domestica_T1R2
Mus_musculus_T1R2_NM_031873_1
Pyxicephalus_adspersus_T1R2
Rhinella_marina_T1R2
Latimeria_chalumnae_T1R2_XM_005986117_2
Latimeria_chalumnae_T1R2_XM_005986116_1
Erpetoichthys_calabaricus_T1R2
Polypterus_senegalus_T1R2_scaffold6
Amia_calva_T1R2
Callorhinchus_milii_T1R6.1
Callorhinchus_milii_T1R6.2_NW_006890089_1
Callorhinchus_milii_T1R6.3_XM_007894997_1
T1R2B
T1R2
Teleost ancestor
Bowfin, gars and teleost ancestor
Ray-finned fish ancestor
B
T1R3B
Xiphophorus_maculatus_T1R3B_XM_005816443_2_modified
Xiphophorus_couchianus_T1R3B_XM_028001776_1_modified
Gambusia_holbrooki_T1R3B
Poeciliopsis_retropinna_T1R3B
Poecilia_reticulata_T1R3B_XM_008408583_2
Anableps_anableps_T1R3B
Kryptolebias_marmoratus_T1R3B_XM_017408636_3_modified
Nothobranchius_furzeri_T1R3B
Oryzias_javanicus_T1R3B
Oryzias_melastigma_T1R3B_XM_024270680_2_modified
Oryzias_latipes_T1R3B_NM_001104725_1
Astatotilapia_calliptera_T1R3B
Maylandia_zebra_T1R3B
Gasterosteus_aculeatus_T1R3B_XM_040205158_1_modified
Perca_fluviatilis_T1R3B_XM_039798392_1_modified
Takifugu_rubripes_T1R3B_NM_001078629_1
Takifugu_rubripes_T1R3B_XM_003973342_1
Notolabrus_celidotus_T1R3B_XM_034687716_1_modified
Periophthalmus_magnuspinnatus_T1R3B_XM_033967258_1
Gadus_morhua_T1R3B_XM_030375363_1
Salmo_salar_T1R3B_XM_045692096_1_modified
Salmo_salar_T1R3B_XM_014166808_1_modified
Cyprinus_carpio_T1R3B_XM_042733715_1_modified
Cyprinus_carpio_T1R3B_XM_019112250_2_modified
Ctenopharyngodon_idella_T1R3B
Danio_rerio_T1R3B_NM_001039628_1
Danionella_translucida_T1R3B
Pygocentrus_nattereri_T1R3B_XM_017683647_1_modified
Piaractus_brachypomus_T1R3B
Astyanax_mexicanus_T1R3B_XM_007251524_2_modified
Ictalurus_punctatus_T1R3B_XM_017451129_1_modified
Electrophorus_electricus_T1R3B_XM_027017506_1
Paramormyrops_kingsleyae_T1R3B_XM_023837597_1
Scleropages_formosus_T1R3B_XM_018725392_1
Lepisosteus_oculatus_T1R3B
Atractosteus_spatula_T1R3B_scaffold523
Amia_calva_T1R3B
Acipenser_ruthenus_T1R3B_XM_034904762_1_modified
Acipenser_ruthenus_T1R3B_XM_034916979_1_modified
Erpetoichthys_calabaricus_T1R3B
Polypterus_senegalus_T1R3B_scaffold_6_ver2
Latimeria_chalumnae_T1R3B_XM_014494968_1
Latimeria_chalumnae_T1R3B_XM_014495881_1
Rhinatrema_bivittatum_T1R3B1_extraction
Pyxicephalus_adspersus_T1R3B1
Pyxicephalus_adspersus_T1R3B2
Rhinella_marina_T1R3B2
Rhinatrema_bivittatum_T1R3B2
Alligator_mississippiensis_T1R3
Gallus_gallus_T1R3_KM091452_1
Anolis_carolinensis_T1R3_XM_003228934_3
Monodelphis_domestica_T1R3
Mus_musculus_T1R3_NM_031872_2
Neoceratodus_forsteri_T1R3_extraction
Callorhinchus_milii_T1R3C
T1R3B
T1R3
Teleost ancestor
Bowfin, gars and teleost ancestor
Ray-finned fish ancestor
C
Teleost ancestor
Sucrose
Fructose
Sucralose
Alanine
Arginine
Proline
Bowfin, gars and teleost ancestor
Ray-finned fish ancestor
Receptor activity
Concentration (mM)
50
0
0.1
1
10
100

**fig. S14. Dose-dependent responses of ancestral T1R2B-T1R3B pairs inferred from three phylogenetic nodes.** (**A, B**) Tree topologies used to infer *T1R2B* (A) and *T1R3B* genes (B) from the ancestors of all teleosts (triangle), of bowfin, gars and teleosts (circle) and all ray-finned fishes (dotted circle) were modified from the maximum-likelihood trees (fig. S12-13; details see Methods). (**C**) T1R2B-T1R3B receptor pairs from the ancestors of all teleosts and of bowfin, gars and teleost ancestor show dose-dependent responses to sugars, sucralose and amino acids, but the pair from the ray-finned fish ancestor appeared nonfunctional. Receptor activity is measured as AUC (x$10^3$ RLU; mean ± s.e.m; n = 6-8; significance (* $p < 0.05$) estimated with one-tailed Welch's *t*-tests and BH corrections as in fig. S3; ligands and concentrations as fig. S3).

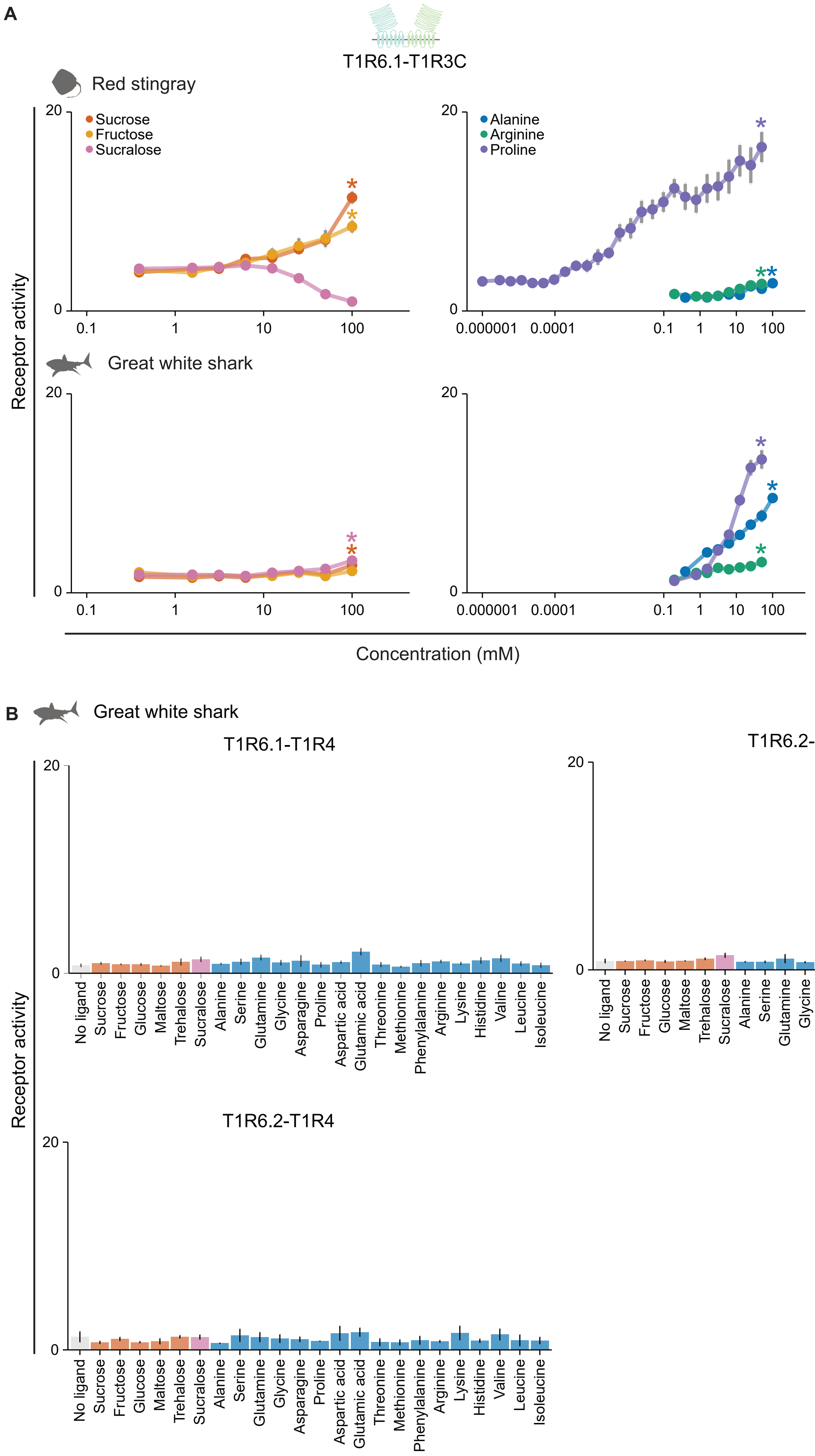
A
T1R6.1-T1R3C
Red stingray
Sucrose
Fructose
Sucralose
Alanine
Arginine
Proline
Great white shark
Receptor activity
Concentration (mM)
B
Great white shark
T1R6.1-T1R4
T1R6.2-T1R3C
T1R6.2-T1R4
No ligand
Sucrose
Fructose
Glucose
Maltose
Trehalose
Sucralose
Alanine
Serine
Glutamine
Glycine
Asparagine
Proline
Aspartic acid
Glutamic acid
Threonine
Methionine
Phenylalanine
Arginine
Lysine
Histidine
Valine
Leucine
Isoleucine

**fig. S15. Functional responses of red stingray (*Hemitrygon akajei*) and great white shark (*Carcharodon carcharias*) receptor pairs.** (**A**) The red stingray T1R6.1-T1R3C shows dose-dependent responses to sucrose, fructose and three amino acids, whereas the great white shark receptor pair is primarily proline and alanine, with weak responses to arginine, sucralose and sucrose. Sucralose may activate the red stingray T1R6.1-T1R3C at lower concentrations than what we measured or act as an inverse agonist, which remains to be verified in future studies. (**B**) Three other receptor pairs of the great white shark did not respond to any sugars or amino acids tested. Receptor activity is calculated as AUC ($x10^3$ RLU; mean ± s.e.m; (A) n = 6; significance (* $p < 0.05$) assessed with Welch's *t*-tests and BH corrections as in fig. S3; ligand concentrations as fig. S3; for the stingray receptor pair, 18 additional lower concentrations of proline were also tested (see Methods); (B) n = 3-6; significance assessed with one-tailed Welch's *t*-tests and BH corrections as in fig. S4; ligand concentrations as fig. S4).

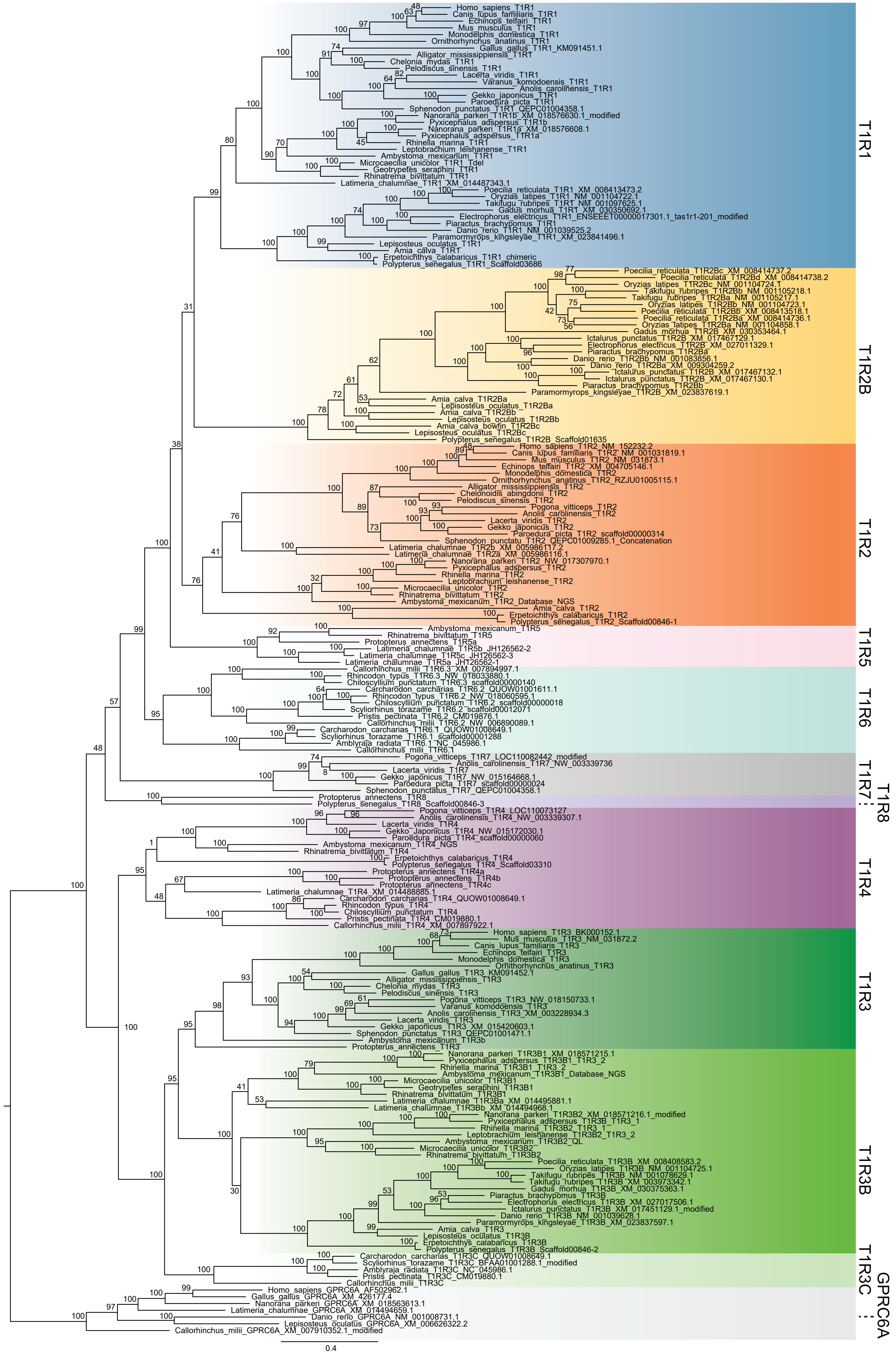
T1R1
Homo_sapiens_T1R1
Canis_lupus_familiaris_T1R1
Echinops_telfairi_T1R1
Mus_musculus_T1R1
Monodelphis_domestica_T1R1
Ornithorhynchus_anatinus_T1R1
Gallus_gallus_T1R1_KM091451.1
Alligator_mississippiensis_T1R1
Chelonia_mydas_T1R1
Pelodiscus_sinensis_T1R1
Lacerta_viridis_T1R1
Varanus_komodoensis_T1R1
Anolis_carolinensis_T1R1
Gekko_japonicus_T1R1
Paroedura_picta_T1R1
Sphenodon_punctatus_T1R1_QEPC01004358.1
Nanorana_parkeri_T1R1b_XM_018576630.1_modified
Pyxicephalus_adspersus_T1R1b
Nanorana_parkeri_T1R1a_XM_018576608.1
Pyxicephalus_adspersus_T1R1a
Rhinella_marina_T1R1
Leptobrachium_leishanense_T1R1
Ambystoma_mexicanum_T1R1
Microcaecilia_unicolor_T1R1_Tdel
Geotrypetes_seraphini_T1R1
Rhinatrema_bivittatum_T1R1
Latimeria_chalumnae_T1R1_XM_014487343.1
Poecilia_reticulata_T1R1_XM_008413473.2
Oryzias_latipes_T1R1_NM_001104722.1
Takifugu_rubripes_T1R1_NM_001097625.1
Gadus_morhua_T1R1_XM_030350692.1
Electrophorus_electricus_T1R1_ENSEEET00000017301.1_tas1r1-201_modified
Piaractus_brachypomus_T1R1
Danio_rerio_T1R1_NM_001039525.2
Paramormyrops_kingsleyae_T1R1_XM_023841496.1
Lepisosteus_oculatus_T1R1
Amia_calva_T1R1
Erpetoichthys_calabaricus_T1R1_chimeric
Polypterus_senegalus_T1R1_Scaffold03686
T1R2B
Poecilia_reticulata_T1R2Bc_XM_008414737.2
Poecilia_reticulata_T1R2Bd_XM_008414738.2
Oryzias_latipes_T1R2Bc_NM_001104724.1
Takifugu_rubripes_T1R2Bb_NM_001105218.1
Takifugu_rubripes_T1R2Ba_NM_001105217.1
Oryzias_latipes_T1R2Bb_NM_001104723.1
Poecilia_reticulata_T1R2Bb_XM_008413518.1
Poecilia_reticulata_T1R2Ba_XM_008414736.1
Oryzias_latipes_T1R2Ba_NM_001104858.1
Gadus_morhua_T1R2B_XM_030353464.1
Ictalurus_punctatus_T1R2B_XM_017467129.1
Electrophorus_electricus_T1R2B_XM_027011329.1
Piaractus_brachypomus_T1R2Ba
Danio_rerio_T1R2Bb_NM_001083856.1
Danio_rerio_T1R2Ba_XM_009304259.2
Ictalurus_punctatus_T1R2B_XM_017467132.1
Ictalurus_punctatus_T1R2B_XM_017467130.1
Piaractus_brachypomus_T1R2Bb
Paramormyrops_kingsleyae_T1R2B_XM_023837619.1
Amia_calva_T1R2Ba
Lepisosteus_oculatus_T1R2Ba
Amia_calva_T1R2Bb
Lepisosteus_oculatus_T1R2Bb
Amia_calva_bowfin_T1R2Bc
Lepisosteus_oculatus_T1R2Bc
Polypterus_senegalus_T1R2B_Scaffold01635
T1R2
Homo_sapiens_T1R2_NM_152232.2
Canis_lupus_familiaris_T1R2_NM_001031819.1
Mus_musculus_T1R2_NM_031873.1
Echinops_telfairi_T1R2_XM_004705146.1
Monodelphis_domestica_T1R2
Ornithorhynchus_anatinus_T1R2_RZJU01005115.1
Alligator_mississippiensis_T1R2
Chelonoidis_abingdonii_T1R2
Pelodiscus_sinensis_T1R2
Pogona_vitticeps_T1R2
Anolis_carolinensis_T1R2
Lacerta_viridis_T1R2
Gekko_japonicus_T1R2
Paroedura_picta_T1R2_scaffold00000314
Sphenodon_punctatu_T1R2_QEPC01009285.1_Concatenation
Latimeria_chalumnae_T1R2b_XM_005986117.2
Latimeria_chalumnae_T1R2a_XM_005986116.1
Nanorana_parkeri_T1R2_NW_017307970.1
Pyxicephalus_adspersus_T1R2
Rhinella_marina_T1R2
Leptobrachium_leishanense_T1R2
Microcaecilia_unicolor_T1R2
Rhinatrema_bivittatum_T1R2
Ambystoma_mexicanum_T1R2_Database_NGS
Amia_calva_T1R2
Erpetoichthys_calabaricus_T1R2
Polypterus_senegalus_T1R2_Scaffold00846-1
T1R5
Ambystoma_mexicanum_T1R5
Rhinatrema_bivittatum_T1R5
Protopterus_annectens_T1R5a
Latimeria_chalumnae_T1R5b_JH126562-2
Latimeria_chalumnae_T1R5c_JH126562-3
Latimeria_chalumnae_T1R5a_JH126562-1
T1R6
Callorhinchus_milii_T1R6.3_XM_007894997.1
Rhincodon_typus_T1R6.3_NW_018033880.1
Chiloscyllium_punctatum_T1R6.3_scaffold00000140
Carcharodon_carcharias_T1R6.2_QUOW01001611.1
Rhincodon_typus_T1R6.2_NW_018060595.1
Chiloscyllium_punctatum_T1R6.2_scaffold00000018
Scyliorhinus_torazame_T1R6.2_scaffold00012071
Pristis_pectinata_T1R6.2_CM019876.1
Callorhinchus_milii_T1R6.2_NW_006890089.1
Carcharodon_carcharias_T1R6.1_QUOW01008649.1
Scyliorhinus_torazame_T1R6.1_scaffold00001288
Amblyraja_radiata_T1R6.1_NC_045986.1
Callorhinchus_milii_T1R6.1
T1R7
Pogona_vitticeps_T1R7_LOC110082442_modified
Anolis_carolinensis_T1R7_NW_003339736
Lacerta_viridis_T1R7
Gekko_japonicus_T1R7_NW_015164668.1
Paroedura_picta_T1R7_scaffold00000024
Sphenodon_punctatus_T1R7_QEPC01004358.1
T1R8
Protopterus_annectens_T1R8
Polypterus_senegalus_T1R8_Scaffold00846-3
T1R4
Pogona_vitticeps_T1R4_LOC110073127
Anolis_carolinensis_T1R4_NW_003339307.1
Lacerta_viridis_T1R4
Gekko_Japonicus_T1R4_NW_015172030.1
Paroedura_picta_T1R4_scaffold00000060
Ambystoma_mexicanum_T1R4_NGS
Rhinatrema_bivittatum_T1R4
Erpetoichthys_calabaricus_T1R4
Polypterus_senegalus_T1R4_Scaffold03310
Protopterus_annectens_T1R4a
Protopterus_annectens_T1R4b
Protopterus_annectens_T1R4c
Latimeria_chalumnae_T1R4_XM_014488885.1
Carcharodon_carcharias_T1R4_QUOW01008649.1
Rhincodon_typus_T1R4
Chiloscyllium_punctatum_T1R4
Pristis_pectinata_T1R4_CM019880.1
Callorhinchus_milii_T1R4_XM_007897922.1
T1R3
Homo_sapiens_T1R3_BK000152.1
Mus_musculus_T1R3_NM_031872.2
Canis_lupus_familiaris_T1R3
Echinops_telfairi_T1R3
Monodelphis_domestica_T1R3
Ornithorhynchus_anatinus_T1R3
Gallus_gallus_T1R3_KM091452.1
Alligator_mississippiensis_T1R3
Chelonia_mydas_T1R3
Pelodiscus_sinensis_T1R3
Pogona_vitticeps_T1R3_NW_018150733.1
Varanus_komodoensis_T1R3
Anolis_carolinensis_T1R3_XM_003228934.3
Lacerta_viridis_T1R3
Gekko_japonicus_T1R3_XM_015420603.1
Sphenodon_punctatus_T1R3_QEPC01001471.1
Ambystoma_mexicanum_T1R3b
Protopterus_annectens_T1R3
T1R3B
Nanorana_parkeri_T1R3B1_XM_018571215.1
Pyxicephalus_adspersus_T1R3B1_T1R3_2
Rhinella_marina_T1R3B1_T1R3_2
Ambystoma_mexicanum_T1R3B1_Database_NGS
Microcaecilia_unicolor_T1R3B1
Geotrypetes_seraphini_T1R3B1
Rhinatrema_bivittatum_T1R3B1
Latimeria_chalumnae_T1R3Ba_XM_014495881.1
Latimeria_chalumnae_T1R3Bb_XM_014494968.1
Nanorana_parkeri_T1R3B2_XM_018571216.1_modified
Pyxicephalus_adspersus_T1R3B_T1R3_1
Rhinella_marina_T1R3B2_T1R3_1
Leptobrachium_leishanense_T1R3B2_T1R3_2
Ambystoma_mexicanum_T1R3B2_QL
Microcaecilia_unicolor_T1R3B2
Rhinatrema_bivittatum_T1R3B2
Poecilia_reticulata_T1R3B_XM_008408583.2
Oryzias_latipes_T1R3B_NM_001104725.1
Takifugu_rubripes_T1R3B_NM_001078629.1
Takifugu_rubripes_T1R3B_XM_003973342.1
Gadus_morhua_T1R3B_XM_030375363.1
Piaractus_brachypomus_T1R3B
Electrophorus_electricus_T1R3B_XM_027017506.1
Ictalurus_punctatus_T1R3B_XM_017451129.1_modified
Danio_rerio_T1R3B_NM_001039628.1
Paramormyrops_kingsleyae_T1R3B_XM_023837597.1
Amia_calva_T1R3
Lepisosteus_oculatus_T1R3B
Erpetoichthys_calabaricus_T1R3B
Polypterus_senegalus_T1R3B_Scaffold00846-2
T1R3C
Carcharodon_carcharias_T1R3C_QUOW01008649.1
Scyliorhinus_torazame_T1R3C_BFAA01001288.1_modified
Amblyraja_radiata_T1R3C_NC_045986.1
Pristis_pectinata_T1R3C_CM019880.1
Callorhinchus_milii_T1R3C
GPRC6A
Homo_sapiens_GPRC6A_AF502962.1
Gallus_gallus_GPRC6A_XM_426177.4
Nanorana_parkeri_GPRC6A_XM_018563613.1
Latimeria_chalumnae_GPRC6A_XM_014494659.1
Danio_rerio_GPRC6A_NM_001008731.1
Lepisosteus_oculatus_GPRC6A_XM_006626322.2
Callorhinchus_milii_GPRC6A_XM_007910352.1_modified
0.4

**fig. S16. Maximum-likelihood tree of 190 *T1R*s and seven *GPRC6A*s of jawed vertebrates.** The maximum-likelihood tree was constructed using a full set of 197 amino acid sequences of 11 clades of *T1R*s and *GPRC6A*s (as outgroups) of 44 jawed vertebrate species using RAxML with the JTT + CAT model. Branch supports represent bootstrap values calculated with 1,000 replicates. The *T1R* clades are colored and their names are shown on the right side. Our naming convention here followed our previous study (*29*) and used numbers to refer to ancient *T1R* duplicates and lower case letters to refer to copies within a species, not to orthologs across different lineages.

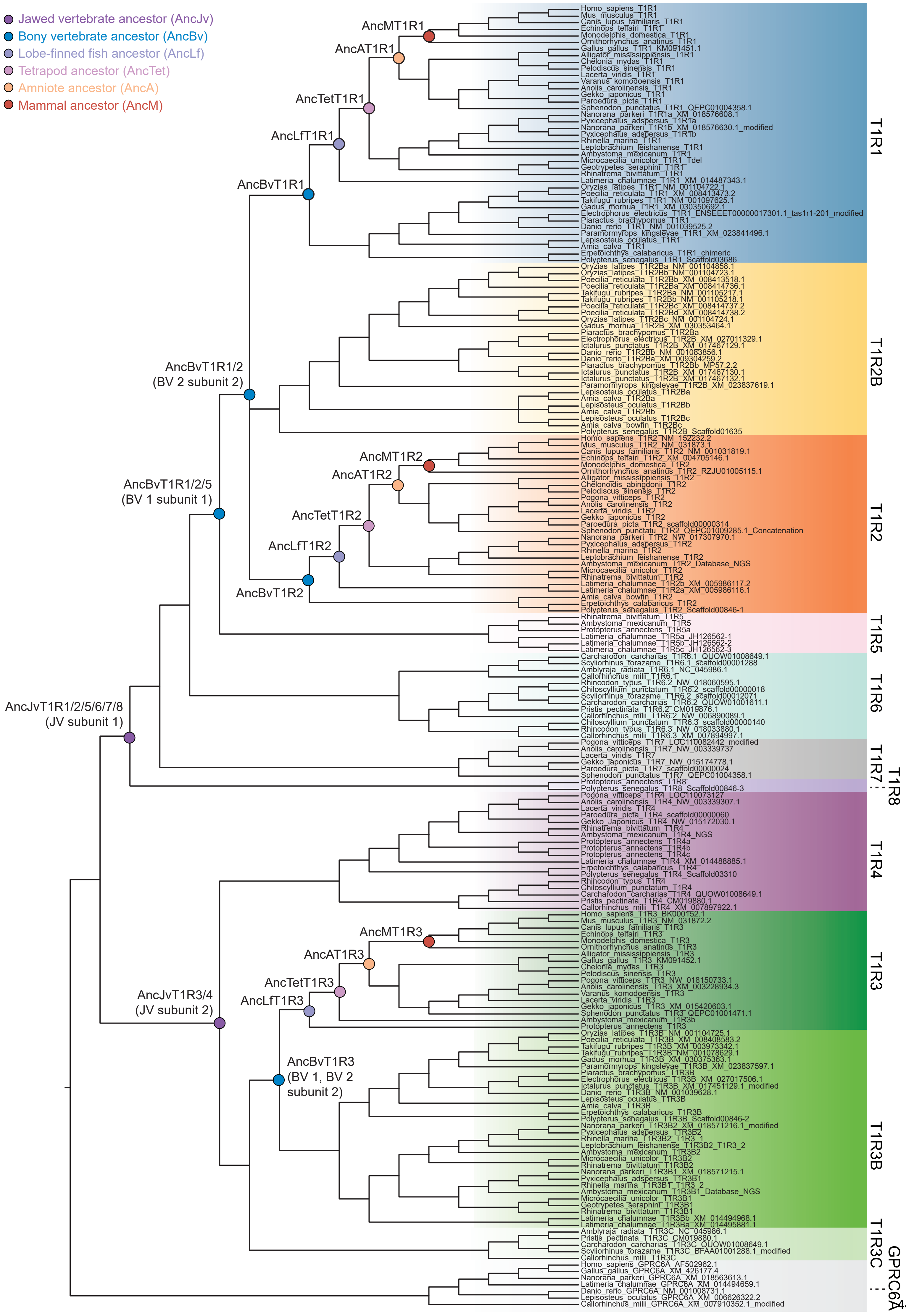

Jawed vertebrate ancestor (AncJv)
Bony vertebrate ancestor (AncBv)
Lobe-finned fish ancestor (AncLf)
Tetrapod ancestor (AncTet)
Amniote ancestor (AncA)
Mammal ancestor (AncM)
AncMT1R1
AncAT1R1
AncTetT1R1
AncLfT1R1
AncBvT1R1
AncBvT1R1/2
(BV 2 subunit 2)
AncBvT1R1/2/5
(BV 1 subunit 1)
AncMT1R2
AncAT1R2
AncTetT1R2
AncLfT1R2
AncBvT1R2
AncJvT1R1/2/5/6/7/8
(JV subunit 1)
AncMT1R3
AncAT1R3
AncTetT1R3
AncLfT1R3
AncJvT1R3/4
(JV subunit 2)
AncBvT1R3
(BV 1, BV 2
subunit 2)
Homo_sapiens_T1R1
Mus_musculus_T1R1
Canis_lupus_familiaris_T1R1
Echinops_telfairi_T1R1
Monodelphis_domestica_T1R1
Ornithorhynchus_anatinus_T1R1
Gallus_gallus_T1R1_KM091451.1
Alligator_mississippiensis_T1R1
Chelonia_mydas_T1R1
Pelodiscus_sinensis_T1R1
Lacerta_viridis_T1R1
Varanus_komodoensis_T1R1
Anolis_carolinensis_T1R1
Gekko_japonicus_T1R1
Paroedura_picta_T1R1
Sphenodon_punctatus_T1R1_QEPC01004358.1
Nanorana_parkeri_T1R1a_XM_018576608.1
Pyxicephalus_adspersus_T1R1a
Nanorana_parkeri_T1R1b_XM_018576630.1_modified
Pyxicephalus_adspersus_T1R1b
Rhinella_marina_T1R1
Leptobrachium_leishanense_T1R1
Ambystoma_mexicanum_T1R1
Microcaecilia_unicolor_T1R1_Tdel
Geotrypetes_seraphini_T1R1
Rhinatrema_bivittatum_T1R1
Latimeria_chalumnae_T1R1_XM_014487343.1
Oryzias_latipes_T1R1_NM_001104722.1
Poecilia_reticulata_T1R1_XM_008413473.2
Takifugu_rubripes_T1R1_NM_001097625.1
Gadus_morhua_T1R1_XM_030350692.1
Electrophorus_electricus_T1R1_ENSEEET00000017301.1_tas1r1-201_modified
Piaractus_brachypomus_T1R1
Danio_rerio_T1R1_NM_001039525.2
Paramormyrops_kingsleyae_T1R1_XM_023841496.1
Lepisosteus_oculatus_T1R1
Amia_calva_T1R1
Erpetoichthys_calabaricus_T1R1_chimeric
Polypterus_senegalus_T1R1_Scaffold03686
Oryzias_latipes_T1R2Ba_NM_001104858.1
Oryzias_latipes_T1R2Bb_NM_001104723.1
Poecilia_reticulata_T1R2Bb_XM_008413518.1
Poecilia_reticulata_T1R2Ba_XM_008414736.1
Takifugu_rubripes_T1R2Ba_NM_001105217.1
Takifugu_rubripes_T1R2Bb_NM_001105218.1
Poecilia_reticulata_T1R2Bc_XM_008414737.2
Poecilia_reticulata_T1R2Bd_XM_008414738.2
Oryzias_latipes_T1R2Bc_NM_001104724.1
Gadus_morhua_T1R2B_XM_030353464.1
Piaractus_brachypomus_T1R2Ba
Electrophorus_electricus_T1R2B_XM_027011329.1
Ictalurus_punctatus_T1R2B_XM_017467129.1
Danio_rerio_T1R2Bb_NM_001083856.1
Danio_rerio_T1R2Ba_XM_009304259.2
Piaractus_brachypomus_T1R2Bb_MP57.2.2
Ictalurus_punctatus_T1R2B_XM_017467130.1
Ictalurus_punctatus_T1R2B_XM_017467132.1
Paramormyrops_kingsleyae_T1R2B_XM_023837619.1
Lepisosteus_oculatus_T1R2Ba
Amia_calva_T1R2Ba
Lepisosteus_oculatus_T1R2Bb
Amia_calva_T1R2Bb
Lepisosteus_oculatus_T1R2Bc
Amia_calva_bowfin_T1R2Bc
Polypterus_senegalus_T1R2B_Scaffold01635
Homo_sapiens_T1R2_NM_152232.2
Mus_musculus_T1R2_NM_031873.1
Canis_lupus_familiaris_T1R2_NM_001031819.1
Echinops_telfairi_T1R2_XM_004705146.1
Monodelphis_domestica_T1R2
Ornithorhynchus_anatinus_T1R2_RZJU01005115.1
Alligator_mississippiensis_T1R2
Chelonoidis_abingdonii_T1R2
Pelodiscus_sinensis_T1R2
Pogona_vitticeps_T1R2
Anolis_carolinensis_T1R2
Lacerta_viridis_T1R2
Gekko_japonicus_T1R2
Paroedura_picta_T1R2_scaffold00000314
Sphenodon_punctatu_T1R2_QEPC01009285.1_Concatenation
Nanorana_parkeri_T1R2_NW_017307970.1
Pyxicephalus_adspersus_T1R2
Rhinella_marina_T1R2
Leptobrachium_leishanense_T1R2
Ambystoma_mexicanum_T1R2_Database_NGS
Microcaecilia_unicolor_T1R2
Rhinatrema_bivittatum_T1R2
Latimeria_chalumnae_T1R2b_XM_005986117.2
Latimeria_chalumnae_T1R2a_XM_005986116.1
Amia_calva_bowfin_T1R2
Erpetoichthys_calabaricus_T1R2
Polypterus_senegalus_T1R2_Scaffold00846-1
Rhinatrema_bivittatum_T1R5
Ambystoma_mexicanum_T1R5
Protopterus_annectens_T1R5a
Latimeria_chalumnae_T1R5a_JH126562-1
Latimeria_chalumnae_T1R5b_JH126562-2
Latimeria_chalumnae_T1R5c_JH126562-3
Carcharodon_carcharias_T1R6.1_QUOW01008649.1
Scyliorhinus_torazame_T1R6.1_scaffold00001288
Amblyraja_radiata_T1R6.1_NC_045986.1
Callorhinchus_milii_T1R6.1
Rhincodon_typus_T1R6.2_NW_018060595.1
Chiloscyllium_punctatum_T1R6.2_scaffold00000018
Scyliorhinus_torazame_T1R6.2_scaffold00012071
Carcharodon_carcharias_T1R6.2_QUOW01001611.1
Pristis_pectinata_T1R6.2_CM019876.1
Callorhinchus_milii_T1R6.2_NW_006890089.1
Chiloscyllium_punctatum_T1R6.3_scaffold00000140
Rhincodon_typus_T1R6.3_NW_018033880.1
Callorhinchus_milii_T1R6.3_XM_007894997.1
Pogona_vitticeps_T1R7_LOC110082442_modified
Anolis_carolinensis_T1R7_NW_003339737
Lacerta_viridis_T1R7
Gekko_japonicus_T1R7_NW_015174778.1
Paroedura_picta_T1R7_scaffold00000024
Sphenodon_punctatus_T1R7_QEPC01004358.1
Protopterus_annectens_T1R8
Polypterus_senegalus_T1R8_Scaffold00846-3
Pogona_vitticeps_T1R4_LOC110073127
Anolis_carolinensis_T1R4_NW_003339307.1
Lacerta_viridis_T1R4
Paroedura_picta_T1R4_scaffold00000060
Gekko_Japonicus_T1R4_NW_015172030.1
Rhinatrema_bivittatum_T1R4
Ambystoma_mexicanum_T1R4_NGS
Protopterus_annectens_T1R4a
Protopterus_annectens_T1R4b
Protopterus_annectens_T1R4c
Latimeria_chalumnae_T1R4_XM_014488885.1
Erpetoichthys_calabaricus_T1R4
Polypterus_senegalus_T1R4_Scaffold03310
Rhincodon_typus_T1R4
Chiloscyllium_punctatum_T1R4
Carcharodon_carcharias_T1R4_QUOW01008649.1
Pristis_pectinata_T1R4_CM019880.1
Callorhinchus_milii_T1R4_XM_007897922.1
Homo_sapiens_T1R3_BK000152.1
Mus_musculus_T1R3_NM_031872.2
Canis_lupus_familiaris_T1R3
Echinops_telfairi_T1R3
Monodelphis_domestica_T1R3
Ornithorhynchus_anatinus_T1R3
Alligator_mississippiensis_T1R3
Gallus_gallus_T1R3_KM091452.1
Chelonia_mydas_T1R3
Pelodiscus_sinensis_T1R3
Pogona_vitticeps_T1R3_NW_018150733.1
Anolis_carolinensis_T1R3_XM_003228934.3
Varanus_komodoensis_T1R3
Lacerta_viridis_T1R3
Gekko_japonicus_T1R3_XM_015420603.1
Sphenodon_punctatus_T1R3_QEPC01001471.1
Ambystoma_mexicanum_T1R3b
Protopterus_annectens_T1R3
Oryzias_latipes_T1R3B_NM_001104725.1
Poecilia_reticulata_T1R3B_XM_008408583.2
Takifugu_rubripes_T1R3B_XM_003973342.1
Takifugu_rubripes_T1R3B_NM_001078629.1
Gadus_morhua_T1R3B_XM_030375363.1
Paramormyrops_kingsleyae_T1R3B_XM_023837597.1
Piaractus_brachypomus_T1R3B
Electrophorus_electricus_T1R3B_XM_027017506.1
Ictalurus_punctatus_T1R3B_XM_017451129.1_modified
Danio_rerio_T1R3B_NM_001039628.1
Lepisosteus_oculatus_T1R3B
Amia_calva_T1R3B
Erpetoichthys_calabaricus_T1R3B
Polypterus_senegalus_T1R3B_Scaffold00846-2
Nanorana_parkeri_T1R3B2_XM_018571216.1_modified
Pyxicephalus_adspersus_T1R3B2
Rhinella_marina_T1R3B2_T1R3_1
Leptobrachium_leishanense_T1R3B2_T1R3_2
Ambystoma_mexicanum_T1R3B2
Microcaecilia_unicolor_T1R3B2
Rhinatrema_bivittatum_T1R3B2
Nanorana_parkeri_T1R3B1_XM_018571215.1
Pyxicephalus_adspersus_T1R3B1
Rhinella_marina_T1R3B1_T1R3_2
Ambystoma_mexicanum_T1R3B1_Database_NGS
Microcaecilia_unicolor_T1R3B1
Geotrypetes_seraphini_T1R3B1
Rhinatrema_bivittatum_T1R3B1
Latimeria_chalumnae_T1R3Bb_XM_014494968.1
Latimeria_chalumnae_T1R3Ba_XM_014495881.1
Amblyraja_radiata_T1R3C_NC_045986.1
Pristis_pectinata_T1R3C_CM019880.1
Carcharodon_carcharias_T1R3C_QUOW01008649.1
Scyliorhinus_torazame_T1R3C_BFAA01001288.1_modified
Callorhinchus_milii_T1R3C
Homo_sapiens_GPRC6A_AF502962.1
Gallus_gallus_GPRC6A_XM_426177.4
Nanorana_parkeri_GPRC6A_XM_018563613.1
Latimeria_chalumnae_GPRC6A_XM_014494659.1
Danio_rerio_GPRC6A_NM_001008731.1
Lepisosteus_oculatus_GPRC6A_XM_006626322.2
Callorhinchus_milii_GPRC6A_XM_007910352.1_modified
T1R1
T1R2B
T1R2
T1R5
T1R6
T1R7
T1R8
T1R4
T1R3
T1R3B
T1R3C
GPRC6A

**fig. S17. *T1R* tree topology (species tree with the full alignment) and phylogenetic nodes for ancestral sequence reconstruction across jawed vertebrates.** The *T1R* tree topology with *GPRC6A*s as the outgroup was adjusted from the maximum-likelihood tree in fig. S16 following the most parsimonious gain and loss scenarios and known species relationships (details in Methods). *T1R1*, *T1R2* and *T1R2B* were depicted as a polytomy due to low bootstrap support values (<50). Ancestral receptors across jawed vertebrates marked as colored circles were reconstructed and tested as heterodimers in functional assays. Names in brackets correspond to those in Fig. 4B.

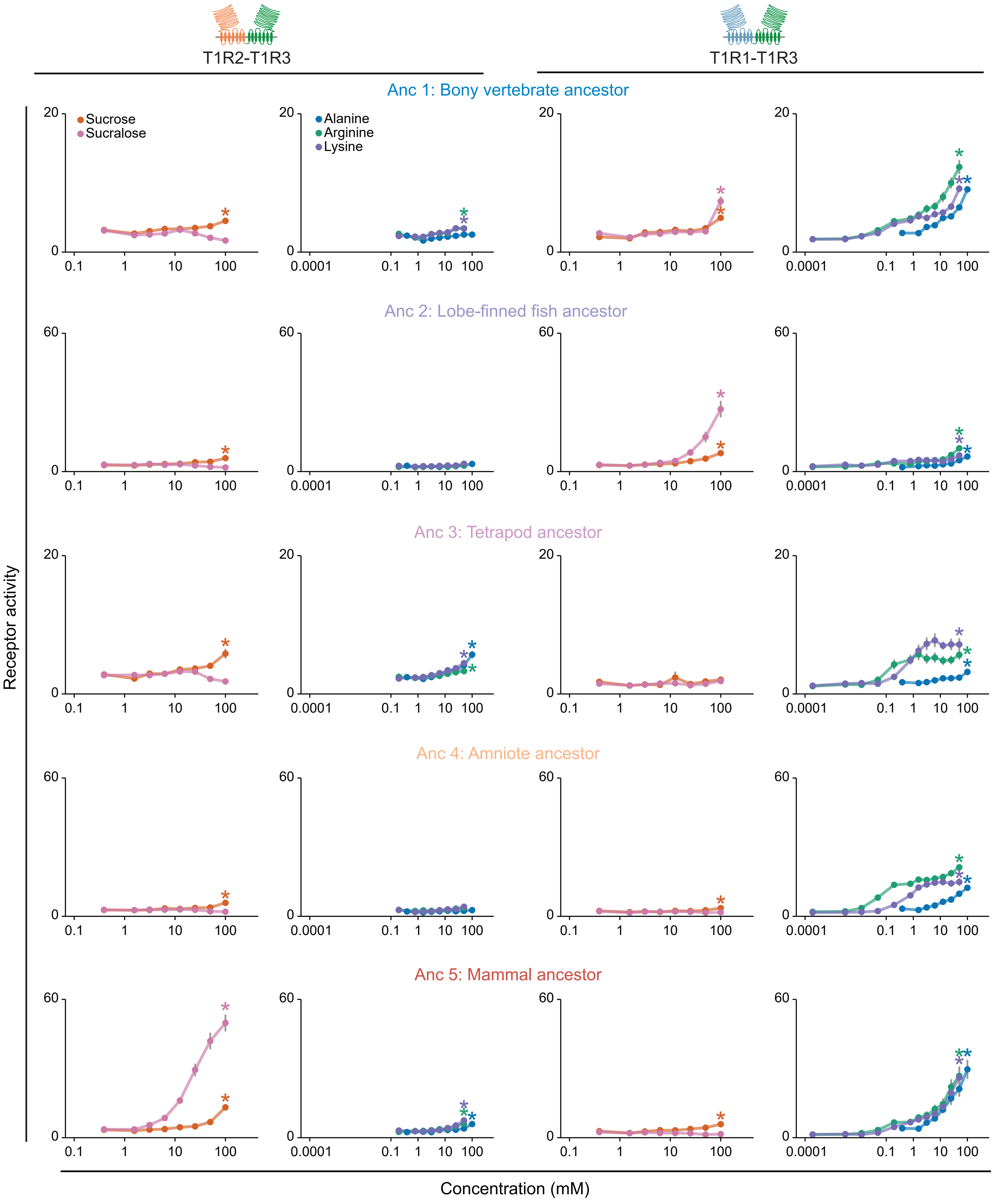

T1R2-T1R3
T1R1-T1R3
Anc 1: Bony vertebrate ancestor
Anc 2: Lobe-finned fish ancestor
Anc 3: Tetrapod ancestor
Anc 4: Amniote ancestor
Anc 5: Mammal ancestor
Sucrose
Sucralose
Alanine
Arginine
Lysine
Receptor activity
Concentration (mM)

**fig. S18. Dose-dependent responses of ancestral receptor pairs across bony vertebrates.** Most ancestral heterodimers show bifunctional responses: T1R2-T1R3 shows a gradual heightened tuning to sucrose (left), whereas T1R1-T1R3 displays an elevated tuning to amino acids across bony vertebrates (right). Notably, T1R2-T1R3 of the mammal ancestor displays the highest relative responses to sucralose and sucrose compared to amino acids and T1R1-T1R3 of all mammals also shows most specialization towards amino acid detection. Receptor activity is measured as AUC (x$10^3$ RLU; mean ± s.e.m; n = 6-10; * $p < 0.05$ assessed with one-tailed Welch's *t*-tests and BH corrections as in fig. S3; ligands and concentrations as in fig. S3, except for five ancestral T1R1-T1R3 pairs which were also tested additionally with four lower concentrations (0.049, 0.012, 0.0031, 0.00019 mM) of arginine and proline). The ancestor names are colored according to the corresponding nodes in fig. S17.

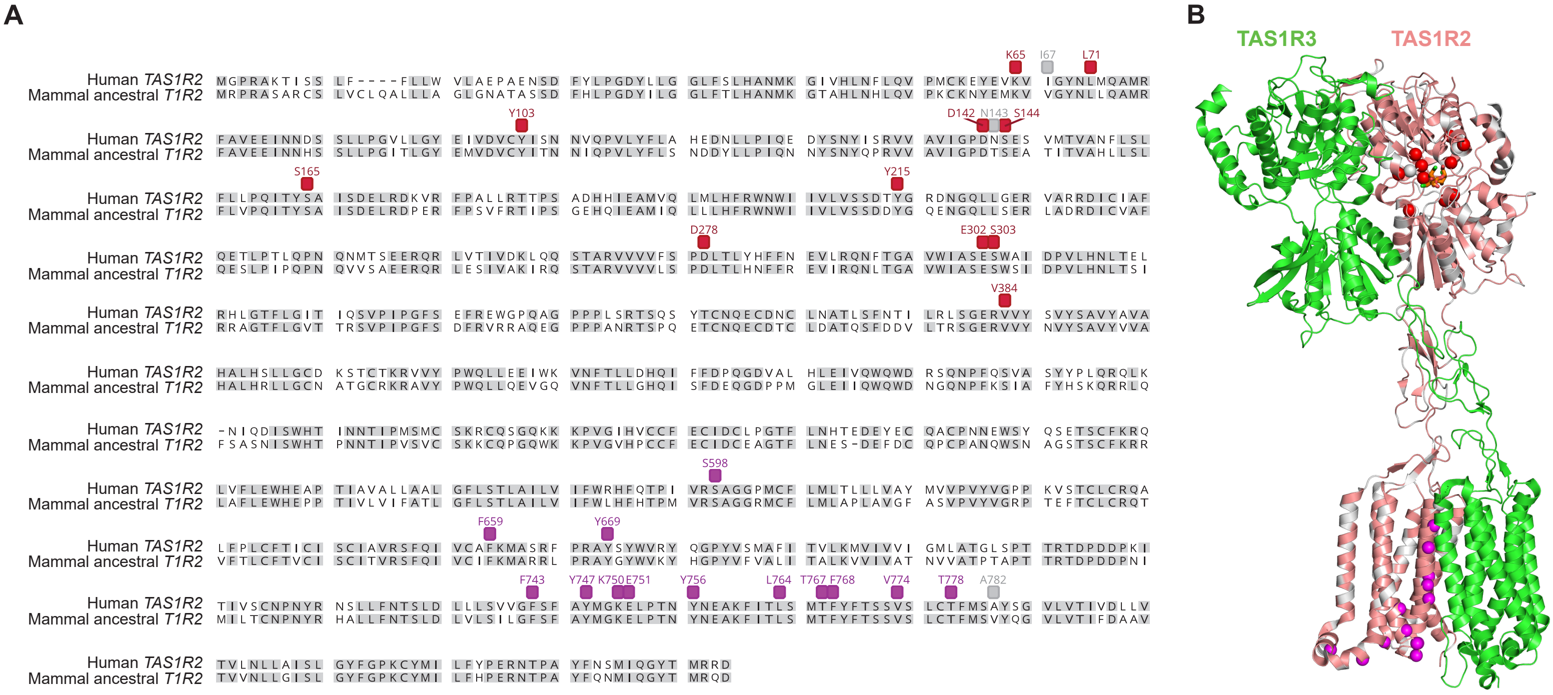
A
Human TAS1R2
Mammal ancestral T1R2
K65
I67
L71
Y103
D142
N143
S144
S165
Y215
D278
E302
S303
V384
S598
F659
Y669
F743
Y747
K750
E751
Y756
L764
T767
F768
V774
T778
A782
B
TAS1R3
TAS1R2

**fig. S19. Amino acid sequence comparison between the human *TAS1R2* (UniProtKB: Q8TE23) and ancestral *T1R2* of all mammals.** (**A**) Human *TAS1R2* and mammal ancestral *T1R2* amino acid alignment and (**B**) sucralose-bounded human sweet taste receptor structure (PDB ID: 9NOT; TAS1R2: salmon; TAS1R3: green; sucralose: red stick model; visualized in Pymol (v3.1.6.1) (*186*)). Most key residues for sucralose binding as well as for heterodimer and G protein interactions (magenta) identified in human *TAS1R2* from previous studies (*79*, *80*) are also present in the inferred mammal ancestral *T1R2*, except for three residues (gray) that differ between human and the mammal ancestral sequences (human *TAS1R2* numbering). Pairwise alignment was conducted using MUSCLE (v3.8.425) (*187*) (implemented in Geneious (v11.1.5; https://www.geneious.com) (*188*). Gray and white shading in (A) signify amino acid residues that are identical or distinct between the two sequences.

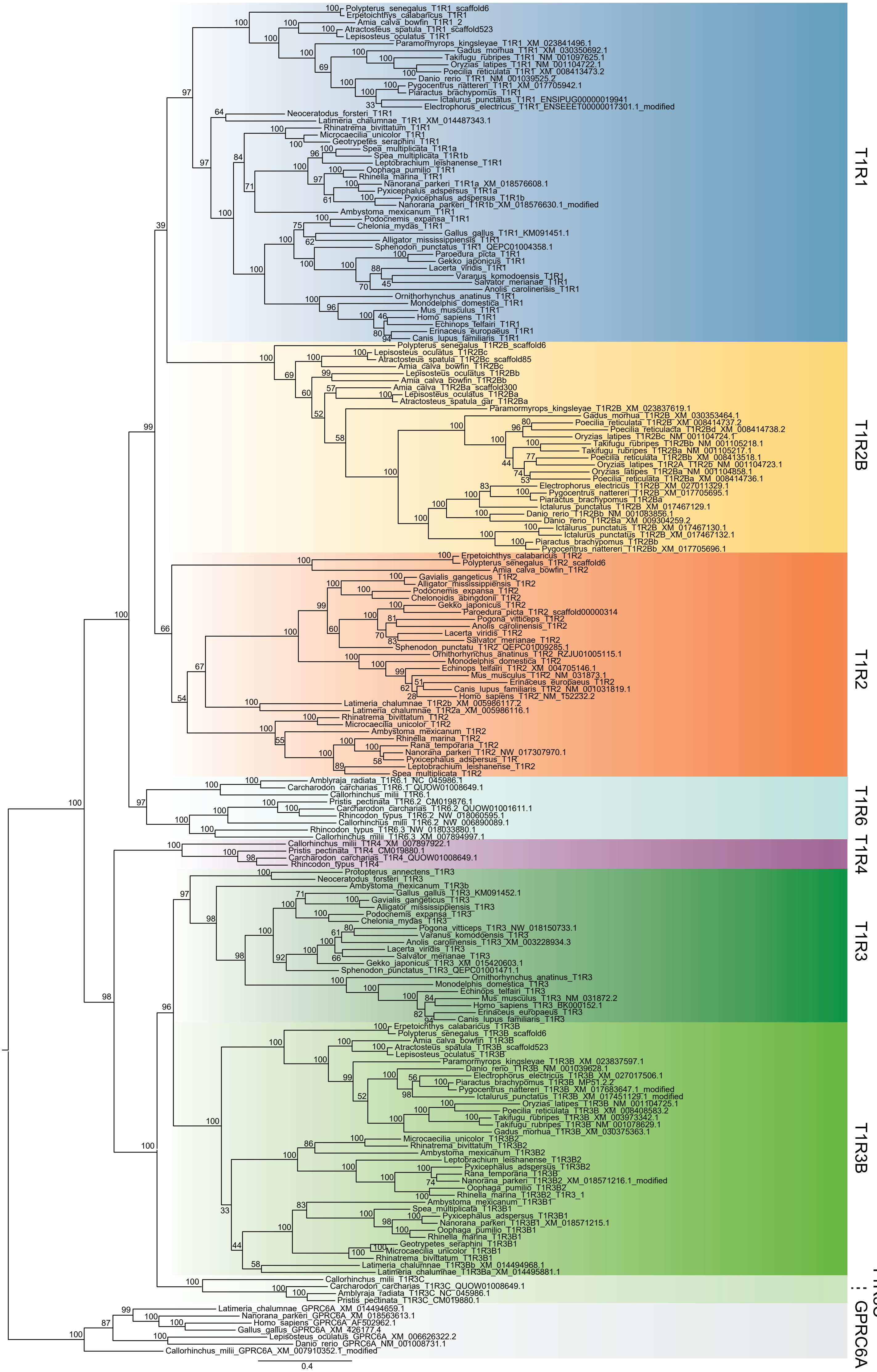

Polypterus_senegalus_T1R1_scaffold6
Erpetoichthys_calabaricus_T1R1
Amia_calva_bowfin_T1R1_2
Atractosteus_spatula_T1R1_scaffold523
Lepisosteus_oculatus_T1R1
Paramormyrops_kingsleyae_T1R1_XM_023841496.1
Gadus_morhua_T1R1_XM_030350692.1
Takifugu_rubripes_T1R1_NM_001097625.1
Oryzias_latipes_T1R1_NM_001104722.1
Poecilia_reticulata_T1R1_XM_008413473.2
Danio_rerio_T1R1_NM_001039525.2
Pygocentrus_nattereri_T1R1_XM_017705942.1
Piaractus_brachypomus_T1R1
Ictalurus_punctatus_T1R1_ENSIPUG00000019941
Electrophorus_electricus_T1R1_ENSEEET00000017301.1_modified
Neoceratodus_forsteri_T1R1
Latimeria_chalumnae_T1R1_XM_014487343.1
Rhinatrema_bivittatum_T1R1
Microcaecilia_unicolor_T1R1
Geotrypetes_seraphini_T1R1
Spea_multiplicata_T1R1a
Spea_multiplicata_T1R1b
Leptobrachium_leishanense_T1R1
Oophaga_pumilio_T1R1
Rhinella_marina_T1R1
Nanorana_parkeri_T1R1a_XM_018576608.1
Pyxicephalus_adspersus_T1R1a
Pyxicephalus_adspersus_T1R1b
Nanorana_parkeri_T1R1b_XM_018576630.1_modified
Ambystoma_mexicanum_T1R1
Podocnemis_expansa_T1R1
Chelonia_mydas_T1R1
Gallus_gallus_T1R1_KM091451.1
Alligator_mississippiensis_T1R1
Sphenodon_punctatus_T1R1_QEPC01004358.1
Paroedura_picta_T1R1
Gekko_japonicus_T1R1
Lacerta_viridis_T1R1
Varanus_komodoensis_T1R1
Salvator_merianae_T1R1
Anolis_carolinensis_T1R1
Ornithorhynchus_anatinus_T1R1
Monodelphis_domestica_T1R1
Mus_musculus_T1R1
Homo_sapiens_T1R1
Echinops_telfairi_T1R1
Erinaceus_europaeus_T1R1
Canis_lupus_familiaris_T1R1
Polypterus_senegalus_T1R2B_scaffold6
Lepisosteus_oculatus_T1R2Bc
Atractosteus_spatula_T1R2Bc_scaffold85
Amia_calva_bowfin_T1R2Bc
Lepisosteus_oculatus_T1R2Bb
Amia_calva_bowfin_T1R2Bb
Amia_calva_T1R2Ba_scaffold300
Lepisosteus_oculatus_T1R2Ba
Atractosteus_spatula_gar_T1R2Ba
Paramormyrops_kingsleyae_T1R2B_XM_023837619.1
Gadus_morhua_T1R2B_XM_030353464.1
Poecilia_reticulata_T1R2B_XM_008414737.2
Poecilia_reticulacta_T1R2Bd_XM_008414738.2
Oryzias_latipes_T1R2Bc_NM_001104724.1
Takifugu_rubripes_T1R2Bb_NM_001105218.1
Takifugu_rubripes_T1R2Ba_NM_001105217.1
Poecilia_reticulata_T1R2Bb_XM_008413518.1
Oryzias_latipes_T1R2A_T1R25_NM_001104723.1
Oryzias_latipes_T1R2Ba_NM_001104858.1
Poecilia_reticulata_T1R2Ba_XM_008414736.1
Electrophorus_electricus_T1R2B_XM_027011329.1
Pygocentrus_nattereri_T1R2B_XM_017705695.1
Piaractus_brachypomus_T1R2Ba
Ictalurus_punctatus_T1R2B_XM_017467129.1
Danio_rerio_T1R2Bb_NM_001083856.1
Danio_rerio_T1R2Ba_XM_009304259.2
Ictalurus_punctatus_T1R2B_XM_017467130.1
Ictalurus_punctatus_T1R2B_XM_017467132.1
Piaractus_brachypomus_T1R2Bb
Pygocentrus_nattereri_T1R2Bb_XM_017705696.1
Erpetoichthys_calabaricus_T1R2
Polypterus_senegalus_T1R2_scaffold6
Amia_calva_bowfin_T1R2
Gavialis_gangeticus_T1R2
Alligator_mississippiensis_T1R2
Podocnemis_expansa_T1R2
Chelonoidis_abingdonii_T1R2
Gekko_japonicus_T1R2
Paroedura_picta_T1R2_scaffold00000314
Pogona_vitticeps_T1R2
Anolis_carolinensis_T1R2
Lacerta_viridis_T1R2
Salvator_merianae_T1R2
Sphenodon_punctatu_T1R2_QEPC01009285.1
Ornithorhynchus_anatinus_T1R2_RZJU01005115.1
Monodelphis_domestica_T1R2
Echinops_telfairi_T1R2_XM_004705146.1
Mus_musculus_T1R2_NM_031873.1
Erinaceus_europaeus_T1R2
Canis_lupus_familiaris_T1R2_NM_001031819.1
Homo_sapiens_T1R2_NM_152232.2
Latimeria_chalumnae_T1R2b_XM_005986117.2
Latimeria_chalumnae_T1R2a_XM_005986116.1
Rhinatrema_bivittatum_T1R2
Microcaecilia_unicolor_T1R2
Ambystoma_mexicanum_T1R2
Rhinella_marina_T1R2
Rana_temporaria_T1R2
Nanorana_parkeri_T1R2_NW_017307970.1
Pyxicephalus_adspersus_T1R
Leptobrachium_leishanense_T1R2
Spea_multiplicata_T1R2
Amblyraja_radiata_T1R6.1_NC_045986.1
Carcharodon_carcharias_T1R6.1_QUOW01008649.1
Callorhinchus_milii_T1R6.1
Pristis_pectinata_T1R6.2_CM019876.1
Carcharodon_carcharias_T1R6.2_QUOW01001611.1
Rhincodon_typus_T1R6.2_NW_018060595.1
Callorhinchus_milii_T1R6.2_NW_006890089.1
Rhincodon_typus_T1R6.3_NW_018033880.1
Callorhinchus_milii_T1R6.3_XM_007894997.1
Callorhinchus_milii_T1R4_XM_007897922.1
Pristis_pectinata_T1R4_CM019880.1
Carcharodon_carcharias_T1R4_QUOW01008649.1
Rhincodon_typus_T1R4
Protopterus_annectens_T1R3
Neoceratodus_forsteri_T1R3
Ambystoma_mexicanum_T1R3b
Gallus_gallus_T1R3_KM091452.1
Gavialis_gangeticus_T1R3
Alligator_mississippiensis_T1R3
Podocnemis_expansa_T1R3
Chelonia_mydas_T1R3
Pogona_vitticeps_T1R3_NW_018150733.1
Varanus_komodoensis_T1R3
Anolis_carolinensis_T1R3_XM_003228934.3
Lacerta_viridis_T1R3
Salvator_merianae_T1R3
Gekko_japonicus_T1R3_XM_015420603.1
Sphenodon_punctatus_T1R3_QEPC01001471.1
Ornithorhynchus_anatinus_T1R3
Monodelphis_domestica_T1R3
Echinops_telfairi_T1R3
Mus_musculus_T1R3_NM_031872.2
Homo_sapiens_T1R3_BK000152.1
Erinaceus_europaeus_T1R3
Canis_lupus_familiaris_T1R3
Erpetoichthys_calabaricus_T1R3B
Polypterus_senegalus_T1R3B_scaffold6
Amia_calva_bowfin_T1R3B
Atractosteus_spatula_T1R3B_scaffold523
Lepisosteus_oculatus_T1R3B
Paramormyrops_kingsleyae_T1R3B_XM_023837597.1
Danio_rerio_T1R3B_NM_001039628.1
Electrophorus_electricus_T1R3B_XM_027017506.1
Piaractus_brachypomus_T1R3B_MP51.2.2
Pygocentrus_nattereri_T1R3B_XM_017683647.1_modified
Ictalurus_punctatus_T1R3B_XM_017451129.1_modified
Oryzias_latipes_T1R3B_NM_001104725.1
Poecilia_reticulata_T1R3B_XM_008408583.2
Takifugu_rubripes_T1R3B_XM_003973342.1
Takifugu_rubripes_T1R3B_NM_001078629.1
Gadus_morhua_T1R3B_XM_030375363.1
Microcaecilia_unicolor_T1R3B2
Rhinatrema_bivittatum_T1R3B2
Ambystoma_mexicanum_T1R3B2
Leptobrachium_leishanense_T1R3B2
Pyxicephalus_adspersus_T1R3B2
Rana_temporaria_T1R3B
Nanorana_parkeri_T1R3B2_XM_018571216.1_modified
Oophaga_pumilio_T1R3B2
Rhinella_marina_T1R3B2_T1R3_1
Ambystoma_mexicanum_T1R3B1
Spea_multiplicata_T1R3B1
Pyxicephalus_adspersus_T1R3B1
Nanorana_parkeri_T1R3B1_XM_018571215.1
Oophaga_pumilio_T1R3B1
Rhinella_marina_T1R3B1
Geotrypetes_seraphini_T1R3B1
Microcaecilia_unicolor_T1R3B1
Rhinatrema_bivittatum_T1R3B1
Latimeria_chalumnae_T1R3Bb_XM_014494968.1
Latimeria_chalumnae_T1R3Ba_XM_014495881.1
Callorhinchus_milii_T1R3C
Carcharodon_carcharias_T1R3C_QUOW01008649.1
Amblyraja_radiata_T1R3C_NC_045986.1
Pristis_pectinata_T1R3C_CM019880.1
Latimeria_chalumnae_GPRC6A_XM_014494659.1
Nanorana_parkeri_GPRC6A_XM_018563613.1
Homo_sapiens_GPRC6A_AF502962.1
Gallus_gallus_GPRC6A_XM_426177.4
Lepisosteus_oculatus_GPRC6A_XM_006626322.2
Danio_rerio_GPRC6A_NM_001008731.1
Callorhinchus_milii_GPRC6A_XM_007910352.1_modified
0.4
T1R1
T1R2B
T1R2
T1R6
T1R4
T1R3
T1R3B
T1R3C
GPRC6A

**fig. S20. Maximum-likelihood tree inferred with a reduced alignment set of 175 *T1R*s and seven *GPRC6A*s across jawed vertebrates for ancestral reconstruction.** The maximum-likelihood tree was inferred using a reduced set of 175 *T1R* (from eight *T1R* clades) and seven *GPRC6A* (as outgroup) amino acid sequences across 51 jawed vertebrates with the JTT + CAT model in RAxML. The bootstrap values were calculated with 1,000 replicates.

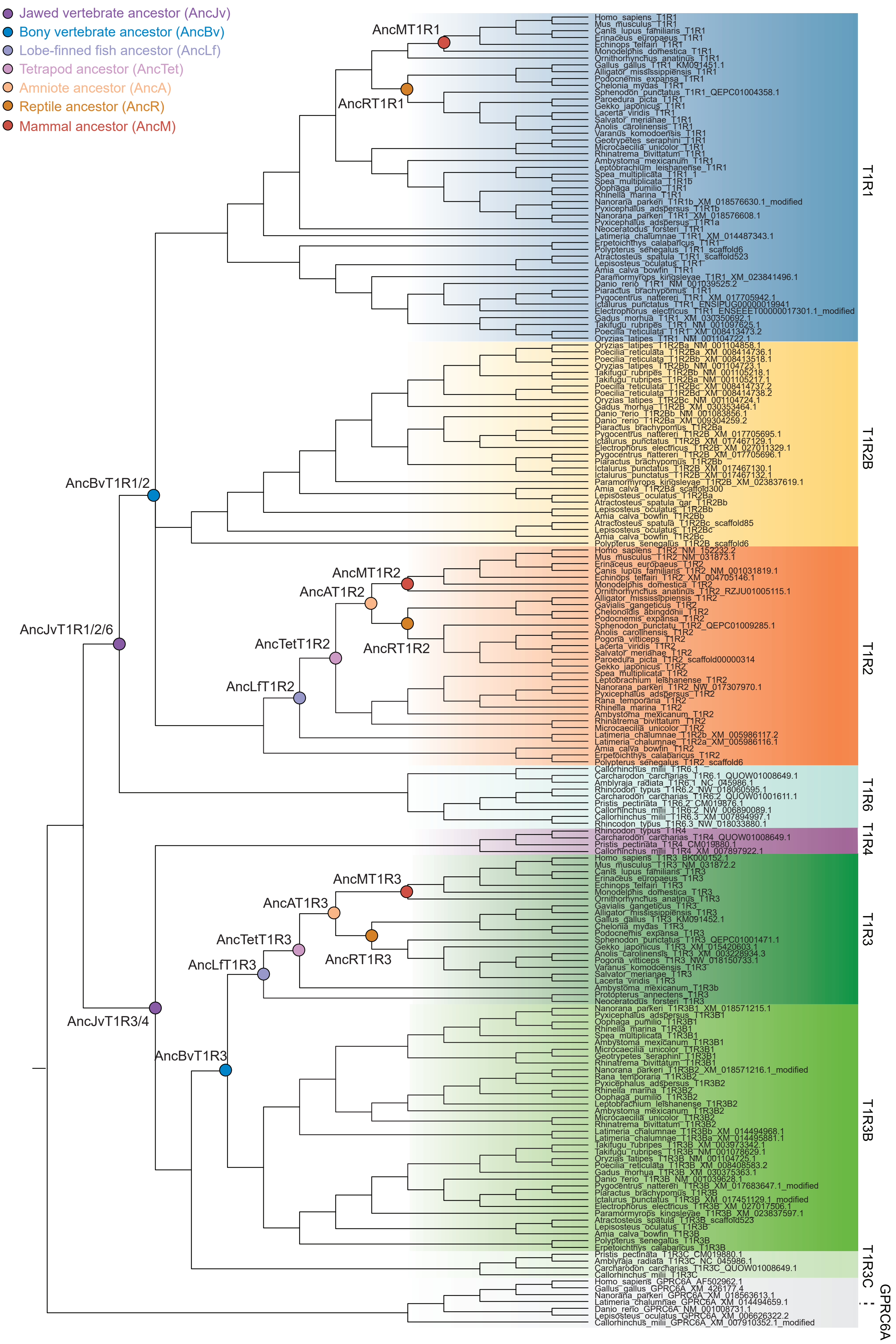

Jawed vertebrate ancestor (AncJv)
Bony vertebrate ancestor (AncBv)
Lobe-finned fish ancestor (AncLf)
Tetrapod ancestor (AncTet)
Amniote ancestor (AncA)
Reptile ancestor (AncR)
Mammal ancestor (AncM)
AncMT1R1
AncRT1R1
AncBvT1R1/2
AncJvT1R1/2/6
AncMT1R2
AncAT1R2
AncTetT1R2
AncRT1R2
AncLfT1R2
AncMT1R3
AncAT1R3
AncTetT1R3
AncRT1R3
AncLfT1R3
AncJvT1R3/4
AncBvT1R3
Homo_sapiens_T1R1
Mus_musculus_T1R1
Canis_lupus_familiaris_T1R1
Erinaceus_europaeus_T1R1
Echinops_telfairi_T1R1
Monodelphis_domestica_T1R1
Ornithorhynchus_anatinus_T1R1
Gallus_gallus_T1R1_KM091451.1
Alligator_mississippiensis_T1R1
Podocnemis_expansa_T1R1
Chelonia_mydas_T1R1
Sphenodon_punctatus_T1R1_QEPC01004358.1
Paroedura_picta_T1R1
Gekko_japonicus_T1R1
Lacerta_viridis_T1R1
Salvator_merianae_T1R1
Anolis_carolinensis_T1R1
Varanus_komodoensis_T1R1
Geotrypetes_seraphini_T1R1
Microcaecilia_unicolor_T1R1
Rhinatrema_bivittatum_T1R1
Ambystoma_mexicanum_T1R1
Leptobrachium_leishanense_T1R1
Spea_multiplicata_T1R1_1
Spea_multiplicata_T1R1b
Oophaga_pumilio_T1R1
Rhinella_marina_T1R1
Nanorana_parkeri_T1R1b_XM_018576630.1_modified
Pyxicephalus_adspersus_T1R1b
Nanorana_parkeri_T1R1_XM_018576608.1
Pyxicephalus_adspersus_T1R1a
Neoceratodus_forsteri_T1R1
Latimeria_chalumnae_T1R1_XM_014487343.1
Erpetoichthys_calabaricus_T1R1
Polypterus_senegalus_T1R1_scaffold6
Atractosteus_spatula_T1R1_scaffold523
Lepisosteus_oculatus_T1R1
Amia_calva_bowfin_T1R1
Paramormyrops_kingsleyae_T1R1_XM_023841496.1
Danio_rerio_T1R1_NM_001039525.2
Piaractus_brachypomus_T1R1
Pygocentrus_nattereri_T1R1_XM_017705942.1
Ictalurus_punctatus_T1R1_ENSIPUG00000019941
Electrophorus_electricus_T1R1_ENSEEET00000017301.1_modified
Gadus_morhua_T1R1_XM_030350692.1
Takifugu_rubripes_T1R1_NM_001097625.1
Poecilia_reticulata_T1R1_XM_008413473.2
Oryzias_latipes_T1R1_NM_001104722.1
Oryzias_latipes_T1R2Ba_NM_001104858.1
Poecilia_reticulata_T1R2Ba_XM_008414736.1
Poecilia_reticulata_T1R2Bb_XM_008413518.1
Oryzias_latipes_T1R2Bb_NM_001104723.1
Takifugu_rubripes_T1R2Bb_NM_001105218.1
Takifugu_rubripes_T1R2Ba_NM_001105217.1
Poecilia_reticulata_T1R2Bc_XM_008414737.2
Poecilia_reticulata_T1R2Bd_XM_008414738.2
Oryzias_latipes_T1R2Bc_NM_001104724.1
Gadus_morhua_T1R2B_XM_030353464.1
Danio_rerio_T1R2Bb_NM_001083856.1
Danio_rerio_T1R2Ba_XM_009304259.2
Piaractus_brachypomus_T1R2Ba
Pygocentrus_nattereri_T1R2B_XM_017705695.1
Ictalurus_punctatus_T1R2B_XM_017467129.1
Electrophorus_electricus_T1R2B_XM_027011329.1
Pygocentrus_nattereri_T1R2B_XM_017705696.1
Piaractus_brachypomus_T1R2Bb
Ictalurus_punctatus_T1R2B_XM_017467130.1
Ictalurus_punctatus_T1R2B_XM_017467132.1
Paramormyrops_kingsleyae_T1R2B_XM_023837619.1
Amia_calva_T1R2Ba_scaffold300
Lepisosteus_oculatus_T1R2Ba
Atractosteus_spatula_gar_T1R2Bb
Lepisosteus_oculatus_T1R2Bb
Amia_calva_bowfin_T1R2Bb
Atractosteus_spatula_T1R2Bc_scaffold85
Lepisosteus_oculatus_T1R2Bc
Amia_calva_bowfin_T1R2Bc
Polypterus_senegalus_T1R2B_scaffold6
Homo_sapiens_T1R2_NM_152232.2
Mus_musculus_T1R2_NM_031873.1
Erinaceus_europaeus_T1R2
Canis_lupus_familiaris_T1R2_NM_001031819.1
Echinops_telfairi_T1R2_XM_004705146.1
Monodelphis_domestica_T1R2
Ornithorhynchus_anatinus_T1R2_RZJU01005115.1
Alligator_mississippiensis_T1R2
Gavialis_gangeticus_T1R2
Chelonoidis_abingdonii_T1R2
Podocnemis_expansa_T1R2
Sphenodon_punctatu_T1R2_QEPC01009285.1
Anolis_carolinensis_T1R2
Pogona_vitticeps_T1R2
Lacerta_viridis_T1R2
Salvator_merianae_T1R2
Paroedura_picta_T1R2_scaffold00000314
Gekko_japonicus_T1R2
Spea_multiplicata_T1R2
Leptobrachium_leishanense_T1R2
Nanorana_parkeri_T1R2_NW_017307970.1
Pyxicephalus_adspersus_T1R2
Rana_temporaria_T1R2
Rhinella_marina_T1R2
Ambystoma_mexicanum_T1R2
Rhinatrema_bivittatum_T1R2
Microcaecilia_unicolor_T1R2
Latimeria_chalumnae_T1R2b_XM_005986117.2
Latimeria_chalumnae_T1R2a_XM_005986116.1
Amia_calva_bowfin_T1R2
Erpetoichthys_calabaricus_T1R2
Polypterus_senegalus_T1R2_scaffold6
Callorhinchus_milii_T1R6.1
Carcharodon_carcharias_T1R6.1_QUOW01008649.1
Amblyraja_radiata_T1R6.1_NC_045986.1
Rhincodon_typus_T1R6.2_NW_018060595.1
Carcharodon_carcharias_T1R6.2_QUOW01001611.1
Pristis_pectinata_T1R6.2_CM019876.1
Callorhinchus_milii_T1R6.2_NW_006890089.1
Callorhinchus_milii_T1R6.3_XM_007894997.1
Rhincodon_typus_T1R6.3_NW_018033880.1
Rhincodon_typus_T1R4
Carcharodon_carcharias_T1R4_QUOW01008649.1
Pristis_pectinata_T1R4_CM019880.1
Callorhinchus_milii_T1R4_XM_007897922.1
Homo_sapiens_T1R3_BK000152.1
Mus_musculus_T1R3_NM_031872.2
Canis_lupus_familiaris_T1R3
Erinaceus_europaeus_T1R3
Echinops_telfairi_T1R3
Monodelphis_domestica_T1R3
Ornithorhynchus_anatinus_T1R3
Gavialis_gangeticus_T1R3
Alligator_mississippiensis_T1R3
Gallus_gallus_T1R3_KM091452.1
Chelonia_mydas_T1R3
Podocnemis_expansa_T1R3
Sphenodon_punctatus_T1R3_QEPC01001471.1
Gekko_japonicus_T1R3_XM_015420603.1
Anolis_carolinensis_T1R3_XM_003228934.3
Pogona_vitticeps_T1R3_NW_018150733.1
Varanus_komodoensis_T1R3
Salvator_merianae_T1R3
Lacerta_viridis_T1R3
Ambystoma_mexicanum_T1R3b
Protopterus_annectens_T1R3
Neoceratodus_forsteri_T1R3
Nanorana_parkeri_T1R3B1_XM_018571215.1
Pyxicephalus_adspersus_T1R3B1
Oophaga_pumilio_T1R3B1
Rhinella_marina_T1R3B1
Spea_multiplicata_T1R3B1
Ambystoma_mexicanum_T1R3B1
Microcaecilia_unicolor_T1R3B1
Geotrypetes_seraphini_T1R3B1
Rhinatrema_bivittatum_T1R3B1
Nanorana_parkeri_T1R3B2_XM_018571216.1_modified
Rana_temporaria_T1R3B2
Pyxicephalus_adspersus_T1R3B2
Rhinella_marina_T1R3B2
Oophaga_pumilio_T1R3B2
Leptobrachium_leishanense_T1R3B2
Ambystoma_mexicanum_T1R3B2
Microcaecilia_unicolor_T1R3B2
Rhinatrema_bivittatum_T1R3B2
Latimeria_chalumnae_T1R3Bb_XM_014494968.1
Latimeria_chalumnae_T1R3Ba_XM_014495881.1
Takifugu_rubripes_T1R3B_XM_003973342.1
Takifugu_rubripes_T1R3B_NM_001078629.1
Oryzias_latipes_T1R3B_NM_001104725.1
Poecilia_reticulata_T1R3B_XM_008408583.2
Gadus_morhua_T1R3B_XM_030375363.1
Danio_rerio_T1R3B_NM_001039628.1
Pygocentrus_nattereri_T1R3B_XM_017683647.1_modified
Piaractus_brachypomus_T1R3B
Ictalurus_punctatus_T1R3B_XM_017451129.1_modified
Electrophorus_electricus_T1R3B_XM_027017506.1
Paramormyrops_kingsleyae_T1R3B_XM_023837597.1
Atractosteus_spatula_T1R3B_scaffold523
Lepisosteus_oculatus_T1R3B
Amia_calva_bowfin_T1R3B
Polypterus_senegalus_T1R3B
Erpetoichthys_calabaricus_T1R3B
Pristis_pectinata_T1R3C_CM019880.1
Amblyraja_radiata_T1R3C_NC_045986.1
Carcharodon_carcharias_T1R3C_QUOW01008649.1
Callorhinchus_milii_T1R3C
Homo_sapiens_GPRC6A_AF502962.1
Gallus_gallus_GPRC6A_XM_426177.4
Nanorana_parkeri_GPRC6A_XM_018563613.1
Latimeria_chalumnae_GPRC6A_XM_014494659.1
Danio_rerio_GPRC6A_NM_001008731.1
Lepisosteus_oculatus_GPRC6A_XM_006626322.2
Callorhinchus_milii_GPRC6A_XM_007910352.1_modified
T1R1
T1R2B
T1R2
T1R6
T1R4
T1R3
T1R3B
T1R3C
GPRC6A

**fig. S21. Species tree topology and phylogenetic nodes for ancestral reconstruction across jawed vertebrates using a reduced alignment.** The species tree topology was modified from the maximum-likelihood tree inferred using a reduced alignment set (also see fig. S20; different from fig. S17 in the number of *T1R* clades). Ancestral T1R subunits marked on each tree were inferred and functionally examined.

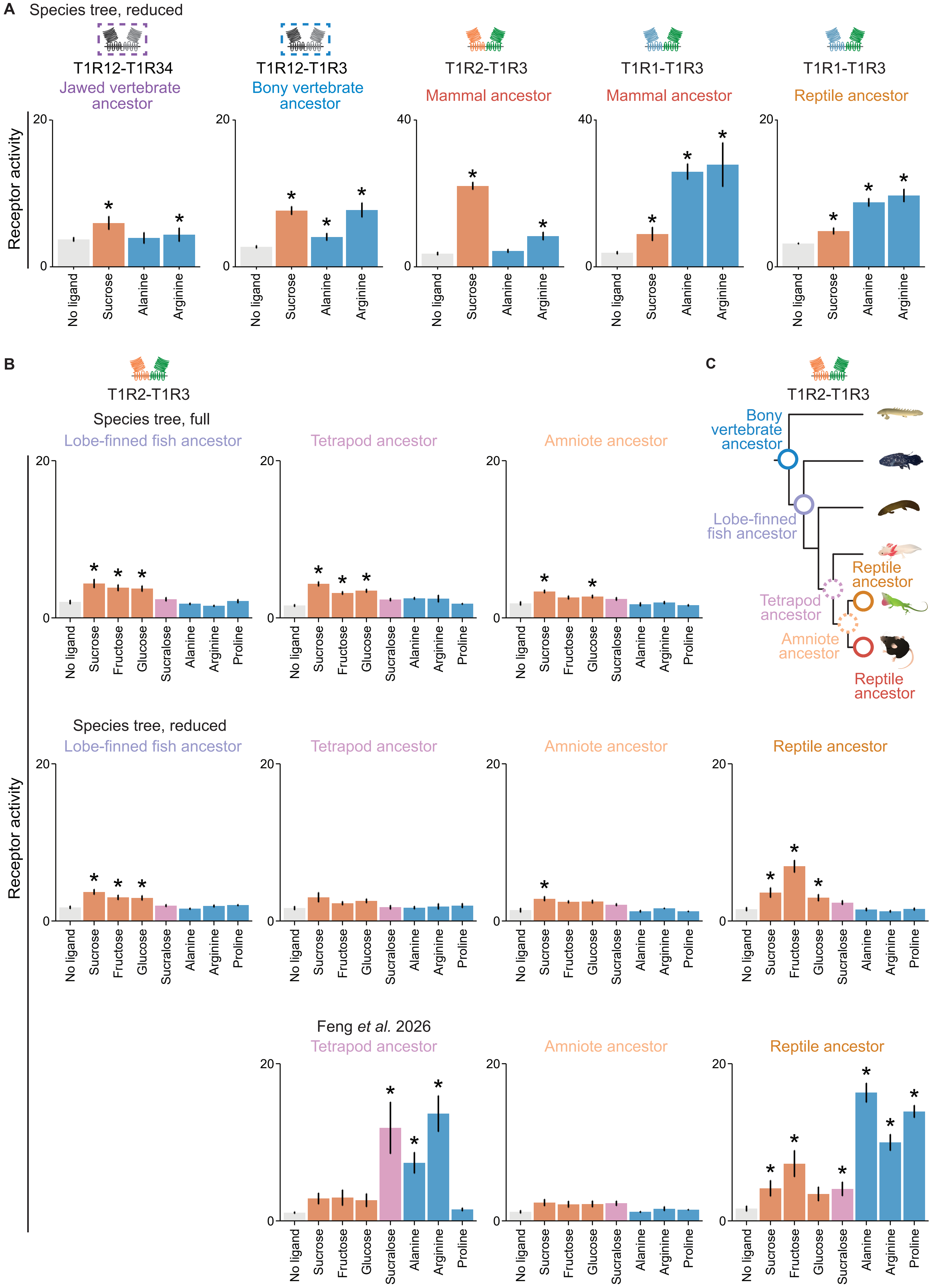
A
Species tree, reduced
T1R12-T1R34
Jawed vertebrate ancestor
T1R12-T1R3
Bony vertebrate ancestor
T1R2-T1R3
Mammal ancestor
T1R1-T1R3
Mammal ancestor
T1R1-T1R3
Reptile ancestor
Receptor activity
20
40
0
No ligand
Sucrose
Alanine
Arginine
*
B
T1R2-T1R3
Species tree, full
Lobe-finned fish ancestor
Tetrapod ancestor
Amniote ancestor
Species tree, reduced
Lobe-finned fish ancestor
Tetrapod ancestor
Amniote ancestor
Reptile ancestor
Feng et al. 2026
Tetrapod ancestor
Amniote ancestor
Reptile ancestor
Receptor activity
No ligand
Sucrose
Fructose
Glucose
Sucralose
Alanine
Arginine
Proline
C
T1R2-T1R3
Bony vertebrate ancestor
Lobe-finned fish ancestor
Reptile ancestor
Tetrapod ancestor
Amniote ancestor
Reptile ancestor

**fig. S22. Ancestral receptor pairs inferred with three different datasets show consistent responses to sucrose.** (**A**) Consistent with the ancestral receptor responses inferred from the full alignment and a species tree topology (species tree, full) (Fig. 4 and fig. S18), ancestral heterodimers inferred from the jawed vertebrate ancestor, the bony vertebrate ancestor and the mammal ancestor inferred with the reduced alignment with a species tree topology (species tree, reduced) show responses to sucrose and some also respond to alanine and arginine. In particular, mammal ancestral T1R2-T1R3 is more sensitive to sucrose than to arginine, whereas its T1R1-T1R3 is more tuned to alanine and arginine than to sucrose (additionally, bifunctional responses are also found in ancestral T1R1-T1R3 of all reptiles). Thus, in this study, the numbers of *T1R* clades included in the alignment sets did not affect the functions of reconstructed ancestral receptor pairs. (**B-C**) Luminescence-based receptor responses and summary of ancestral receptor pairs inferred from two sets in this study and a recent study (Feng *et al.* 2026) (*27*) across bony vertebrates. (B) Ancestral T1R2-T1R3 receptor pairs of all lobe-finned fish inferred from two sets in this study and receptor pairs of all reptiles from the reduced set (with a species tree topology) and from Feng *et al.* (*27*) show significant responses to sugars. The clear presence of sugar sensing in the ancestral receptor pairs of the earlier nodes (bony vertebrate and lobe-finned fish ancestors) as well as in the later nodes (the reptile and mammal ancestors) (solid line circles on the topology in (C)) suggest an early origin of sugar sensing no later than the emergence of bony vertebrates, despite the small or no response to sugars from the ancestral T1R2-T1R3 of all tetrapods and of all amniotes (dotted line circles in (C)). Receptor activity is calculated as AUC (x$10^3$ RLU) ($n$ = 5-12; mean ± s.e.m; significance (* $p < 0.05$) assessed with one-tailed Welch's *t*-tests and BH corrections as in fig. S4; ligand concentrations as fig. S4).
