## Supplementary tables for "Early origin of sugar sensing in jawed vertebrates"

**table S1**. Numbers and percentages of fruit- and seed-taking species across major clades of vertebrates. Related to Fig. 1A and fig. S1A.

| Class | # of total species | # of fruit- & seed-taking species | % of fruit- & seed-taking species | # of fruit-taking species | % of fruit-taking species |
|---|---|---|---|---|---|
| Aves | 11,131 | 5,128 | 46.07 | 3,719 | 33.41 |
| Crocodilia | 27 | 10 | 37.04 | 6 | 22.22 |
| Testudines | 366 | 79 | 21.58 | 76 | 20.77 |
| Lepidosauria | 12,250 | 498 | 4.07 | 482 | 3.93 |
| Mammalia | 6,723 | 2,813 | 41.84 | 2,218 | 32.99 |
| Amphibia | 8,923 | 5 | 0.06 | 2 | 0.02 |
| Dipnoi | 6 | 2 | 33.33 | 1 | 16.67 |
| Coelacanth | 2 | 0 | 0.00 | 0 | 0.00 |
| Actinopterygii | 34,635 | 274 | 0.79 | 215 | 0.62 |
| Chondrichthyes | 1,344 | 0 | 0.00 | 0 | 0.00 |
| Agnatha | 141 | 0 | 0.00 | 0 | 0.00 |

**table S2**. Numbers and percentages of fruit- and seed-taking species across 17 orders of ray-finned fish. Related to Fig. 3A and fig. S1B.

| Order | # of total species | # of fruit- & seed-taking species | % of fruit- & seed-taking species | # of fruit-taking species | % of fruit-taking species |
|---|---|---|---|---|---|
| Cyprinodontiformes | 1,418 | 3 | 0.21 | 3 | 0.21 |
| Beloniformes | 294 | 0 | 0.00 | 0 | 0.00 |
| Atheriniformes | 383 | 1 | 0.26 | 0 | 0.00 |
| Cichliformes | 1,792 | 16 | 0.89 | 12 | 0.67 |
| Anabantiformes | 275 | 4 | 1.45 | 4 | 1.45 |
| Gobiiformes | 2,366 | 2 | 0.08 | 2 | 0.08 |
| Centrarchiformes | 298 | 4 | 1.34 | 3 | 1.01 |
| Tetraodontiformes | 450 | 1 | 0.22 | 1 | 0.22 |
| Characiformes | 2,269 | 124 | 5.46 | 110 | 4.85 |
| Siluriformes | 4,025 | 59 | 1.47 | 48 | 1.19 |
| Gymnotiformes | 256 | 2 | 0.78 | 2 | 0.78 |
| Cypriniformes | 4,732 | 49 | 1.04 | 27 | 0.57 |
| Osteoglossiformes | 256 | 7 | 2.73 | 2 | 0.78 |
| Elopiformes | 9 | 1 | 11.11 | 1 | 11.11 |
| Lepisosteiformes | 7 | 0 | 0.00 | 0 | 0.00 |
| Amiiformes | 2 | 0 | 0.00 | 0 | 0.00 |
| Polypteriformes | 14 | 1 | 7.14 | 0 | 0.00 |

**table S3**. Statistical output from linear mixed models assessing the behavioral sugar preference of tortoises and anoles. Related to Fig. 1C-D and fig S2.

| **Species** | **Response variable** | **# of individuals** | **# of trials** | **Stimulus** | **Time window (min)** | **Estimate** | **Std.error** | **Statistic** | ***p* value** |
|---|---|---|---|---|---|---|---|---|---|
| Red-footed tortoise | Number of bites | 5 | 7 | Sucrose 1 M | 5 | 0.255 | 0.081 | 3.147 | **0.002** |
| | | | | | 10 | 0.364 | 0.076 | 4.809 | **0.000** |
| | | | | | 20 | 0.539 | 0.073 | 7.369 | **0.000** |
| | | | | | 60 | 0.654 | 0.061 | 10.797 | **0.000** |
| | | | | | 150 | 0.741 | 0.055 | 13.509 | **0.000** |
| Green anole | Normalized time drinking | 6 | 34 | Sucrose 0.75 M | 5 | 0.456 | 0.232 | 1.966 | **0.049** |
| | | | | | 10 | 0.545 | 0.221 | 2.462 | **0.014** |
| | | | | | 20 | 0.552 | 0.221 | 2.494 | **0.013** |
| | | | | | 60 | 0.526 | 0.221 | 2.378 | **0.017** |
| | | 2 | 10 | Sucrose 0.5 M | 5 | 1.140 | 0.371 | 3.076 | **0.002** |
| | | | | | 10 | 1.006 | 0.391 | 2.572 | **0.010** |
| | | | | | 20 | 0.916 | 0.406 | 2.258 | **0.024** |
| | | | | | 60 | 1.062 | 0.380 | 2.798 | **0.005** |
| | | 3 | 20 | Sucrose 0.3 M | 5 | 0.808 | 0.292 | 2.766 | **0.006** |
| | | | | | 10 | 0.774 | 0.292 | 2.649 | **0.008** |
| | | | | | 20 | 0.753 | 0.290 | 2.592 | **0.010** |
| | | | | | 60 | 0.874 | 0.279 | 3.138 | **0.002** |
| | | 3 | 15 | Glucose 0.75 M | 5 | 0.829 | 0.336 | 2.468 | **0.014** |
| | | | | | 10 | 0.829 | 0.336 | 2.468 | **0.014** |
| | | | | | 20 | 0.829 | 0.336 | 2.468 | **0.014** |
| | | | | | 60 | 0.776 | 0.337 | 2.302 | **0.021** |
| | | 4 | 25 | Fructose 0.75 M | 5 | 0.301 | 0.282 | 1.065 | 0.287 |
| | | | | | 10 | 0.355 | 0.281 | 1.264 | 0.206 |
| | | | | | 20 | 0.442 | 0.277 | 1.594 | 0.111 |
| | | | | | 60 | 0.536 | 0.275 | 1.951 | 0.051 |

**table S4**. Genome and transcriptome data used for *T1R* curation.

| Species | Data version and accession | Reference |
|---|---|---|
| Human (*Homo sapiens* ) | GRCh38.p13 (NCBI: GCF_000001405.39) | 1 |
| Mouse (*Mus musculus* ) | GRCm38.p6 (NCBI: GCF_000001635.26) | 2 |
| Opossum (*Monodelphis domestica* ) | monDom5 (NCBI: GCF_000002295.1) | 3 |
| Zebra finch (*Taeniopygia guttata* ) | bTaeGut1.4 (NCBI: GCF_003957565.2) | 4 |
| Chicken (*Gallus gallus* ) | GRCg6a (NCBI: GCF_000002315.6) | 5 |
| Kiwi (*Apteryx australis mantelli* ) | AptMant0 (NCBI: GCA_001039765.2) | 6 |
| American alligator (*Alligator mississippiensis* ) | ASM28112v4 (NCBI: GCF_000281125.3) | 7 |
| Chinese alligator (*Alligator sinensis* ) | ASM45574v1 (NCBI: GCF_000455745.1) | 8 |
| Crocodylus porosus (*Australian saltwater crocodile* ) | CroPor_comp1 (NCBI: GCA_001723895.1) | 9 |
| Green sea turtle (*Chelonia mydas* ) | rCheMyd1.pri.v2 (NCBI: GCF_015237465.2) | 10 |
| Western painted turtle (*Chrysemys picta* ) | Chrysemys_picta_BioNano-3.0.4 (NCBI: GCF_000241765.5) | 11 |
| Chinese softshell turtle (*Pelodiscus sinensis* ) | PelSin_1.0 (NCBI: GCF_000230535.1) | 12 |
| Green anole (*Anolis carolinensis* ) | AnoCar2.0 (NCBI: GCF_000090745.1) | 13 |
| Central bearded dragon (*Pogona vitticeps* ) | pvi1.1 (NCBI: GCF_900067755.1) | 14 |
| Japanese gecko (*Gekko japonicas* ) | Gekko_japonicus_V1.1 (NCBI: GCF_001447785.1) | 15 |
| Tuatara (*Sphenodon punctatus* ) | ASM311381v1 (NCBI: GCA_003113815.1) | 16 |
| African bullfrog (*Pyxicephalus adspersus* ) | Pads_1.0 (NCBI: GCA_004786255.1) | 17 |
| Tibetan frog (*Nanorana parkeri* ) | ASM93562v1 (NCBI: GCF_000935625.1) | 18 |
| Axolotl (*Ambystoma mexicanum* ) | AmbMex60DD (NCBI: GCA_002915635.3) | 19 |
| | RNA-seq (NCBI: PRJEB22921) | 20 |
| | RNA-seq (NCBI: PRJNA312389) | 21 |
| | RNA-seq (NCBI: PRJNA354434) | 22 |
| Two-lined caecilian (*Rhinatrema bivittatum* ) | aRhiBiv1.1 (NCBI: GCF_901001135.1) | 4 |
| West African lungfish (*Protopterus annectens* ) | National Genomic Data Center : GWHANVS00000000 | 23 |
| | RNA-seq (NCBI: PRJNA282925) | 24 |
| Australian lungfish (*Neoceratodus forsteri* ) | neoFor_v3 (NCBI: GCA_016271365.1) | 25 |
| Coelacanth (*Latimeria chalumnae* ) | LatCha1 (NCBI: GCF_000225785.1) | 26, 27 |
| | RNA-seq (DDBJ: DRP000627) | |
| Fugu (*Takifugu rubripes* ) | FUGU5 (NCBI: GCF_000180615.1) | 28 |
| Nile tilapia (*Oreochromis niloticus* ) | O_niloticus_UMD_NMBU (NCBI: GCF_001858045.2) | 29 |
| Japanese rice fish (medaka) (*Oryzias latipes* ) | ASM223467v1 (NCBI: GCF_002234675.1) | 30 |
| Guppy (*Poecilia reticulata* ) | Guppy_female_1.0+MT (NCBI: GCF_000633615.1) | 31 |
| Silver salmon (*Oncorhynchus kisutch* ) | Okis_V2 (NCBI: GCF_002021735.2) | 32 |
| Zebrafish (*Danio rerio* ) | GRCz11 (NCBI: GCF_000002035.6) | 33 |
| Mexican tetra (*Astyanax mexicanu* ) | AstMex3_surface (NCBI: GCF_023375975.1) | 34 |
| Red-bellied piranha (*Pygocentrus nattereri* ) | Pygocentrus_nattereri-1.0.2 (NCBI: GCA_001682695.1) | 35 |
| Spotted gar (*Lepisosteus oculatus)* | LepOcu1 (NCBI: GCF_000242695.1) | 36 |
| Alligator gar (*Atractosteus spatula* ) | BGI_Aspa_1.0 (GCA_016984175.1) | 37 |
| Bowfin (*Amia calva* ) | AmiCal1 (NCBI: GCA_017591415.1) | 37 |
| Sterlet (*Acipenser_ruthenus* ) | ASM1064508v1 (NCBI: GCF_010645085.1) | 38 |
| Reedfish (*Erpetoichthys calabaricus* ) | fErpCal1.3 (NCBI: GCA_900747795.4) | 4 |
| Bichir (*Polypterus senegalus* ) | AS1683550v1 (NCBI: GCF_016835505.1) | 37 |
| | polypterus_v0.62 | Okabe et al. (unpublished) |
| Elephant shark (*Callorhinchus milii* ) | Callorhinchus_milii-6.1.3 (NCBI: GCF_000165045.1) | 39 |
| | RNA-seq (NCBI: PRJNA135005) | 40 |
| White shark (*Carcharodon carcharias* ) | ASM360424v1 (NCBI: GCA_003604245.1) | 41 |
| Whale shark (*Rhincodon typus* ) | ASM164234v2 (NCBI: GCF_001642345.1) | 42 |
| Cloudy catshark (*Scyliorhinus torazame* ) | Storazame_v1.0 (NCBI: GCA_003427355.1) | 43 |
| | RNA-seq (NCBI: PRJNA371391) | 44 |
| Brownbanded bamboo shark (*Chiloscyllium punctatum* ) | Cpunctatum_v1.0 (NCBI: GCA_003427335.1) | 43 |
| Atlantic stingray (*Hypanus sabinus* ) | sHypSab1 (NCBI: GCA_030144785.1) | 4 |
| Red stingray (*Hemitrygon akajei* ) | sHemAka1.1 | 45 |
| | sHemAka1.3 (NCBI: GCA_048418815.1) | 46 |
